# Accumulation of Rare Coding Variants in Genes Implicated in Risk of Human Cleft Lip with or without Cleft Palate

**DOI:** 10.1101/533208

**Authors:** Nicholas J. Marini, Kripa Asrani, Wei Yang, Jasper Rine, Gary M Shaw

## Abstract

Cleft lip with/without cleft palate (CLP) is a common craniofacial malformation with complex etiologies, reflecting both genetic and environmental factors. Most of the suspected genetic risk for CLP has yet to be identified. To further classify risk loci and estimate the contribution of rare variants, we sequenced the exons in 49 candidate genes in 323 CLP cases and 211 non-malformed controls. Our findings indicated that rare, protein-altering variants displayed markedly higher burdens in CLP cases at relevant loci. First, putative loss-of-function mutations (nonsense, frameshift) were significantly enriched among cases: 13 of 323 cases (~4%) harbored such alleles within these 49 genes, versus one such change in controls (*p* = 0.01). Second, in gene-level analyses, the burden of rare alleles showed greater case-association for several genes previously implicated in cleft risk. For example, *BHMT* displayed a 10-fold increase in protein-altering variants in CLP cases (*p* = 0.03), including multiple case occurrences of a rare frameshift mutation (K400fs). Other loci with greater rare, coding allele burdens in cases were in signaling pathways relevant to craniofacial development (*WNT9B*, *BMP4*, *BMPR1B*) as well as the methionine cycle (*MTRR*). We conclude that rare coding variants may confer risk for isolated CLP.

## INTRODUCTION

Although their causes are largely unknown, orofacial clefts are suspected of being etiologically heterogeneous with both genetic and non-genetic risk factors [Dixon et al., 2011]. Clinical observations have long indicated that orofacial clefts should be classified into at least two distinct phenotypic groups: cleft lip with or without cleft palate (CLP), and cleft palate alone (CPO) [Fogh-Anderson 1967]. Collectively, these are among the most common birth defects with a worldwide prevalence of approximately 1 in 700 live births [WHO, 2002].

Genetic associations for CLP have been observed for several genes, including IRF6, *FGFR2, FOXE1, MSX1, NECTIN1, BMP4, TBX22*, and *TGFa* [Dixon et al., 2011] from either linkage analysis or candidate gene studies. In addition, genome-wide association studies (GWAS) have identified approximately 15 additional risk loci [Beaty et al., 2010; Birnbaum et al., 2009; Grant et al., 2009; Leslie et al., 2016; Ludwig et al., 2012; Mangold et al., 2010] with some of these replicating in subsequent studies [e.g. – Jia et al., 2015]. For some of these loci, such as *IRF6* and *NOG,* follow-up sequencing efforts have identified compelling functional variants as putative risk alleles [Leslie et al., 2015; Rahimov et al., 2008]. Interestingly, there is apparently little overlap between the most significantly-associated GWAS regions and genes previously implicated in CLP risk.

This is probably indicative of the complex genetic etiologies underlying the CLP phenotype and the difficulties detecting associated loci at genome-wide significance. Thus, there are likely to be numerous additional genetic risk factors yet to be identified. Indeed, the combined efforts of all these genetic approaches have been extremely fruitful, yet still explain only a fraction of the population burden of these human birth defects.

Several non-genetic factors also appear to contribute to cleft phenotypes [Mossey et al., 2009]. Of particular relevance to the current study is that maternal use of multivitamin supplements containing folic acid in early pregnancy has been associated with decreased risk of CLP [Shaw et al., 1995; Wilcox et al., 2007]. Although some reports do not observe a reduction in risk associated with maternal folate supplementation [see Little et al., 2008], a recent meta-analysis indicated a significant reduction in risk of CLP with maternal folic acid use [Jahanbin et al., 2018]. Because of this folic acid link, folate/one-carbon pathway genes have been a logical place to look for risk variants [Boyles et al., 2009; Blanton et al., 2011; Marini et al., 2016]. However, many of the known common pathway SNPs have been evaluated, yet the results are still somewhat unconvincing. The exception to this seems to be the fairly consistent association of the *BHMT/BHMT2/DMGDH* locus on Chromosome 5 with CLP observed in a number of studies [Marini et al., 2016]. Interestingly, while the locus often shows association, the local SNPs driving the association can vary between studies. Thus, the nature of the risk at this locus is still unclear.

In this study we hypothesized that some genetic risk for CLP lies in rare variants and this has contributed to the lack of consistent findings and attendant difficulty in unraveling risk alleles. Thus, we sequenced the exons in 30 genes involved in folate metabolism in a 534-member case-control population to determine if rare variant allele burden (in individual genes, subsets of genes, or the entire pathway) correlated with phenotype. We also sequenced the coding region of an additional 19 candidate genes (not related to folate metabolism) that are involved in lip/palate development or previously associated with CLP. Our results showed that, while there were no folate pathway-wide trends in rare variant allele burden, two genes involved in methionine synthesis (*BHMT* and *MTRR*) showed convincing case-associated trends for protein-altering variants. Likewise, genes involved in BMP/WNT signaling (*BMP4*, *BMPR1B*, *WNT9B*), which have been previously implicated in orofacial cleft development [Zhang et al., 2002; Suzuki et al., 2009; Juriloff et al., 2014], displayed higher allele burdens in cases. CLP cases also presented a greater number of loss-of-function alleles (frameshifts, truncations) in this candidate gene set.

## METHODS

### Study Population

This case-control study included data on deliveries that had estimated due dates from 1995–2003. The study included live-born infants with isolated CLP (N=681), isolated CPO (N=157), or without any structural malformation (controls; N = 706). Because the goal of the study was to determine whether rare coding variants in candidate genes underlie risk for CLP, variant discovery by gene sequencing was performed on an ethnically-mixed subset of this population comprising 323 CLP cases and 211 controls. A second group (358 CLP cases, 157 CPO cases, 485 controls) was used to further validate the CLP case-association of a frameshift mutation in BHMT. See Table I for race-ethnic breakdown of the study population.

**Table I.** Populations Used in Study.

**A. Discovery population**
| **Race/Ethnicity** | **CLP Cases**<br>**(Collective Percentage)** | **Controls**<br>**(Collective Percentage)** |
| --- | --- | --- |
| Asian | 35 (10.8) | 25 (11.8) |
| Black | 13 (4.0) | 8 (3.8) |
| White, Hispanic | 190 (58.8) | 126 (59.7) |
| White, Non-Hispanic | 79 (24.5) | 48 (22.7) |
| Other | 6 (1.9) | 4 (1.9) |
| **TOTAL** | **323** | **211** |

**B. Validation population**
| **Race/Ethnicity** | **CLP Cases**<br>**(Collective Percentage)** | **CPO Cases**<br>**(Collective Percentage)** | **Controls**<br>**(Collective Percentage)** |
| --- | --- | --- | --- |
| Asian | 41 (11.5) | 18 (11.5) | 41 (8.5) |
| Black | 9 (2.5) | 9 (5.7) | 29 (6.0) |
| White, Hispanic | 188 (52.5) | 58 (36.9) | 231 (47.6) |
| White, Non-Hispanic | 54 (15.1) | 32 (20.4) | 83 (17.1) |
| Other | 6 (1.7) | 1 (0.6) | 5 (1.0) |
| No Race Reported | 59 (16.5) | 39 (24.8) | 96 (19.8) |
| Native American | 1 (0.3) |  |  |
| **TOTAL** | **358** | **157** | **485** |

Case information was abstracted from hospital reports and medical records following established procedures by the California Birth Defects Monitoring Program [Croen et al., 1991]. Each medical record was further reviewed by a medical geneticist (Edward J. Lammer, MD). Infants with trisomies were ineligible. Non-malformed control infants were selected randomly to represent the birth population from which the cases were derived in selected counties and birth periods. This study, including the collection and use of archived newborn bloodspots, was approved by the California State Committee for the Protection of Human Subjects as well as Institutional Review Boards at Stanford University and the University of California, Berkeley.

### Genomic DNA Isolation

Genomic DNA (gDNA) was available from newborn screening dried blood spots obtained from linkage efforts made by the California Birth Defects Monitoring Program. gDNA was extracted from dried blood spots and whole-genome amplified using QIAmp DNA Mini kits and Replica-G Midi amplification kits (Qiagen), respectively, as described previously [Marini et al., 2011].

### Illumina Paired-End Sequencing of Enriched Target Exons

3-5 μg of whole-genome amplified DNA was used as starting material for Illumina library construction and target exon capture with Sure Select (Agilent) probes according to the Agilent SureSelect^XT^ protocol (https://www.agilent.com/cs/library/usermanuals/public/G7530-90000.pdf). Briefly, DNA was fragmented with a Covaris sonicator, followed by end-repair and A-tailing, then ligation of sequencing adaptors and PCR amplification. Amplified libraries were then used to perform target exon capture by hybridization against a custom-made capture library of 120 bp biotinylated oligonucleotide probes covering the 555 coding exons in 49 candidate genes of interest (Table S1). Probes (Agilent SureSelect) were designed via the SureDesign web tool using the genomic coordinates of target exons (genome build GRCh37/hg19). To ensure capture and coverage of the entire coding region and splice sites, probes were designed with an additional 50 bp “pad” that covered intronic regions adjacent to coding exons or, in the case of the first or last coding exon, covered the 5’- or 3’- untranslated regions. Following hybridization and streptavidin capture of enriched exons, captured libraries were barcoded with 6 bp tags with an additional round of PCR, quantified, and 12-16 sample libraries were pooled for 100 bp paired-end sequencing on the Illumina HiSeq (v3 chemistry).

### Read Mapping and Variant Filtering

Raw sequence files (FASTQ) were imported into CLC Genomics Workbench (v.6; Qiagen) and mapped onto the human genome (GRCh37/hg19). Only full-length, paired, uniquely-mapping reads with <= 3 mismatches were used for variant calling. Variants with depth of coverage <30, or with genotype quality scores <25, were excluded. In addition, alleles not observed on both strands (with a minimum frequency ≥ 0.1) or with allelic imbalances (heterozygote genotype calls with one allele observed with frequency <0.25 or homozygote alleles observed with frequency < 0.85) were also excluded. After compilation of the dataset, positions not covered in at least 95% of the cases or controls were then removed.

### Variant Calling Accuracy

The accuracy of the variant calls in the filtered dataset was evaluated in two ways: 1) the concordance of genotype calls by Illumina sequencing and by TaqMan allelic discrimination assays (described in Marini et al., 2016) was determined for 8,537 variant calls (mostly common alleles) in 476 samples, and 2) confirmation by Sanger sequencing for the subset of rare variants in genes with striking case-related trends (~15% of all protein-altering variants identified in the study). Verification sequencing was performed on original bloodspot gDNA preparations, prior to whole-genome amplification, using amplicon-specific M13-capped PCR primers designed by VariantSEQr methodology (ThermoFisher) and deposited into the NCBI probe database (https://www.ncbi.nlm.nih.gov/probe/docs/projvariantseqr/). Each sample for verification had a corresponding negative control for comparison.

### Analyses

To estimate the contribution of rare (MAF < 1%) coding variants (for which there may only be a single occurrence) to CLP risk, a simple allele burden test was applied on a gene-by-gene basis to highlight genes with prominent case-control skews. Alleles were summed by gene (or groups of genes) and case-control associations were tested by Fisher’s exact test based on genotype distributions (i.e. carrier frequency). Although, this cannot be used as a strict test, statistical differences in case or control representation are readily observed with this metric. For common polymorphisms, variants were treated as categorical variables, i.e., homozygous wild-type as referent versus heterozygous or homozygous variant. Odds ratios and 95% confidence intervals were used to estimate risks.

## RESULTS

### Dataset Definition, Variant Calling and Quality Control

We took a candidate gene approach, with a focus on folate pathway genes, to investigate the potential contribution of rare coding variants to CLP risk by determining whether putative loss-of-function alleles or rare allele burdens in certain genes (or pathways) correlated with phenotype. Thus we sequenced the coding exons and exon/intron boundaries in 30 genes related to folate metabolism and 19 additional genes involved in craniofacial development in a case-control study (Table I). The full list of genes queried, including exon annotations and coverage statistics are in Table S1. 532 coding exons out of the 555 targeted for enrichment in these 49 genes (96%) passed quality control for adequate coverage and sequence quality. Mean coverage for each of these 532 exons was similar between the case group and the control group, indicating each group was sequenced with equal efficiency (Figure S1).

Variants were called in exons and in intron regions within 10 bp from exon/intron junctions as described in Methods. To evaluate the overall quality and accuracy of the variant calls, we performed 2 types of independent assessments on different subsets of variants. In the first, we established the concordance between the genotypes called from Illumina sequencing with those obtained from TaqMan allelic discrimination assays for many multiply-occurring loci (i.e. not singletons) using TaqMan assays previously described [Marini et al., 2016]. 91 variant positions in 24 genes were compared in 476 samples. Out of 8,537 variant calls, we observed 99.7% concordance between the two methods (Table S2).

In a second quality-control check, we evaluated all putative loss-of-function alleles (frameshifts and truncations), and all rare and singleton missense variant calls from genes whose alleles showed higher case representation, by exon-specific Sanger resequencing (see Methods). Out of 95 tested variants, all were confirmed except one frameshift in a low complexity region, which was subsequently removed (Table S3). Based on these analyses, we are confident in the sequencing accuracy of the data, necessary for rare variant genotyping.

The final analytical dataset contained data for 773 variant positions: 285 synonymous, 380 protein-altering, and 108 non-coding variants. One hundred fifty-five (54%) of the synonymous alleles were singletons, whereas 243 (64%) of the protein-altering alleles were singletons. The complete list of annotated variants, allele frequencies and case-control distributions are in Table S4.

### Common Allele Associations

Although our focus was on the distribution of rare coding alleles, sequence data also provided information on associations of more common variants. We observed a small number of nominally significant associations that are in agreement with previous studies. For example, association of nonsynonymous SNPs in *DMGDH* (S279P, homozygous OR = 0.5 (0.3-0.9), *p* = 0.022) and *ALDH1L1* (I812V, heterozygous OR = 2.7 (1.3-5.5), *p* = 0.008) have been reported previously [Boyles et al., 2009; Marini et al., 2016].

By far, the strongest statistically significant signal of any variant with high enough frequency to calculate was the nonsynonymous SNP V274I in *IRF6* (heterozygous OR = 0.5 (0.3-0.7), *p* = 4E-04; homozygous OR = 0.5 (0.2-0.9), *p* = 0.046), a well-studied polymorphism in orofacial cleft studies with numerous reported associations [Beaty et al., 2016]. A second *IRF6* synonymous SNP (S153S) was also significantly associated in homozygotes (OR = 0.6 (0.4-0.9), *p*=.02). Finally, there were 2 linked synonymous SNPs in the *VANGL2* gene (G445G and P467P; heterozygous OR = 1.8 (1.2-2.7), *p*=.004), which has not previously been associated with CLP.

Odds ratios were calculated only from genotype frequencies in the entire, ethnically diverse population, and not in ethnically stratified sub-populations, because this study was not sufficiently powered for such analyses. Our focus was on rare variants which are often impervious to meaningful ethnic stratification analyses owing to the infrequency of occurrence. However, we have ensured that the race-ethnic breakdown was approximately the same in case and control groups (Table I).

### Association of Putative Loss-Of-Function Alleles

The candidate gene set interrogated in this study is a mix of both folate pathway genes and genes implicated in cleft occurrence because of their roles in craniofacial development. Thus, this set was enriched for potential cleft risk genes over what might be considered in a broader sampling of the exome. Therefore, it is noteworthy that we observed a significant enrichment of putative loss-of-function alleles (frameshifts, truncations) in CLP cases: 13 cases harbored one such allele from this gene set versus only 1 control (Fisher’s exact *p* = 0.01; Table II). All alleles in Table II were seen only once in this population except for the deletion/frameshift at Lysine 400 in BHMT (K400fs), which was seen in 4 cases of differing ethnicity. It is further noteworthy that several of the genes represented in Table II (*BHMT, DMGDH, MTRR, IRF6*) have displayed significant CLP associations in multiple studies (see Discussion). It should be emphasized that functional impact of these alleles is only inferred, though they represent good candidates for mutations of consequence for cleft risk.

**Table II.** Putative Loss-Of-Function Alleles.

| **Gene** | **Variant** | **Type<sup>#</sup>** | **Case Appearance<sup>\*</sup>** | **Control Appearance<sup>\*</sup>** |
| --- | --- | --- | --- | --- |
| *BHMT* | K400fs | deletion/frameshift | Hispanic<br>Hispanic<br>non-Hispanic, White<br>non-Hispanic, White |  |
| *DMGDH* | W495* | SNV/stop-gain | Hispanic |  |
| *GGH* | L190fs | insertion/frameshift | non-Hispanic, White |  |
| *IRF6* | Q112* | SNV/stop-gain | non-Hispanic, White |  |
| *MTRR* | R386* | SNV/stop-gain | non-Hispanic, White |  |
| *SARDH* | R457* | SNV/stop-gain | Hispanic |  |
| *SARDH* | R514* | SNV/stop-gain | Hispanic |  |
| *SP8* | S261* | SNV/stop-gain | Hispanic |  |
| *SHMT1* | R99* | SNV/stop-gain | Hispanic |  |
| *SHMT1* | R207* | SNV/stop-gain | Asian |  |
| *SHMT1* | Y457* | SNV/stop-gain |  | Asian |
| **TOTAL** |  |  | **13** | **1** |

<sup>#</sup> SNV=Single Nucleotide Variation
<sup>\*</sup> Ethnicity of newborn

### Folate Pathway Variation

Despite numerous epidemiological studies suggesting that maternal folic acid supplementation may modify risk for CLP, common SNP association studies with folate/one-carbon pathway variants have not been definitive in linking pathway loci to phenotype. Suggestive evidence exists for several loci from this group [e.g. – Boyles et al., 2009; Blanton et al., 2011; Marini et al., 2016], though reproducibility across populations and small effect sizes remain problematic. The role of rare variants in folate pathway genes has not been adequately queried and may account for genetic risk yet to be identified. Indeed, the folate/one-carbon pathway may be particularly susceptible to the aggregate mutation burden throughout the pathway, since there are many co-dependent and interacting components. Thus, we hypothesized that the aggregate burden of rare variants in the folate pathway (or related genes in pathway sub-compartments) may confer risk for CLP and help explain the folate link. We have found such pathway analyses useful in understanding the role of folate pathway variation, common and rare, with respect to Neural Tube Defect risk [Marini et al., 2011].

By allele summing of rare variants (defined here as MAF < 1%), the group of folate/one-carbon pathway genes (N=30) did not show an increased aggregate burden of either synonymous mutations (case aggregate allele frequency = 0.27, control aggregate allele frequency = 0.28; *p* = 0.57, Mann Whitney test) or intron mutations within 10 bp of the exon boundary (case aggregate allele frequency = 0.12, control aggregate allele frequency = 0.10; *p* = 0.34, Mann Whitney test) that might distinguish cases from controls. There was as slight increase in aggregated protein-altering allele frequency for cases (case aggregate allele frequency = 0.38, control aggregate allele frequency = 0.31); though the distributions were not significantly different (*p* = 0.08).

Upon sub-dividing pathway genes into compartments with metabolically related genes (e.g. – purine synthesis, see [Marini et al., 2011]), it became obvious that the strongest case-associated signals for rare, protein-altering variants were restricted to 2 genes involved in methionine synthesis: *BHMT* (betaine-homocysteine methyl transferase) and *MTRR* (methionine synthase reductase), both of which are necessary for the remethylation of homocysteine to methionine (Fig. 1, Table III). In addition to the 4 case occurrences of the K400fs allele in *BHMT*, nonsynonymous variants were found in an additional 7 cases and only a single control (Fisher’s exact *p* = 0.03). Similarly, protein altering variants for *MTRR* were found in 13 cases versus 2 controls (*p* = 0.06).

**Figure 1.**
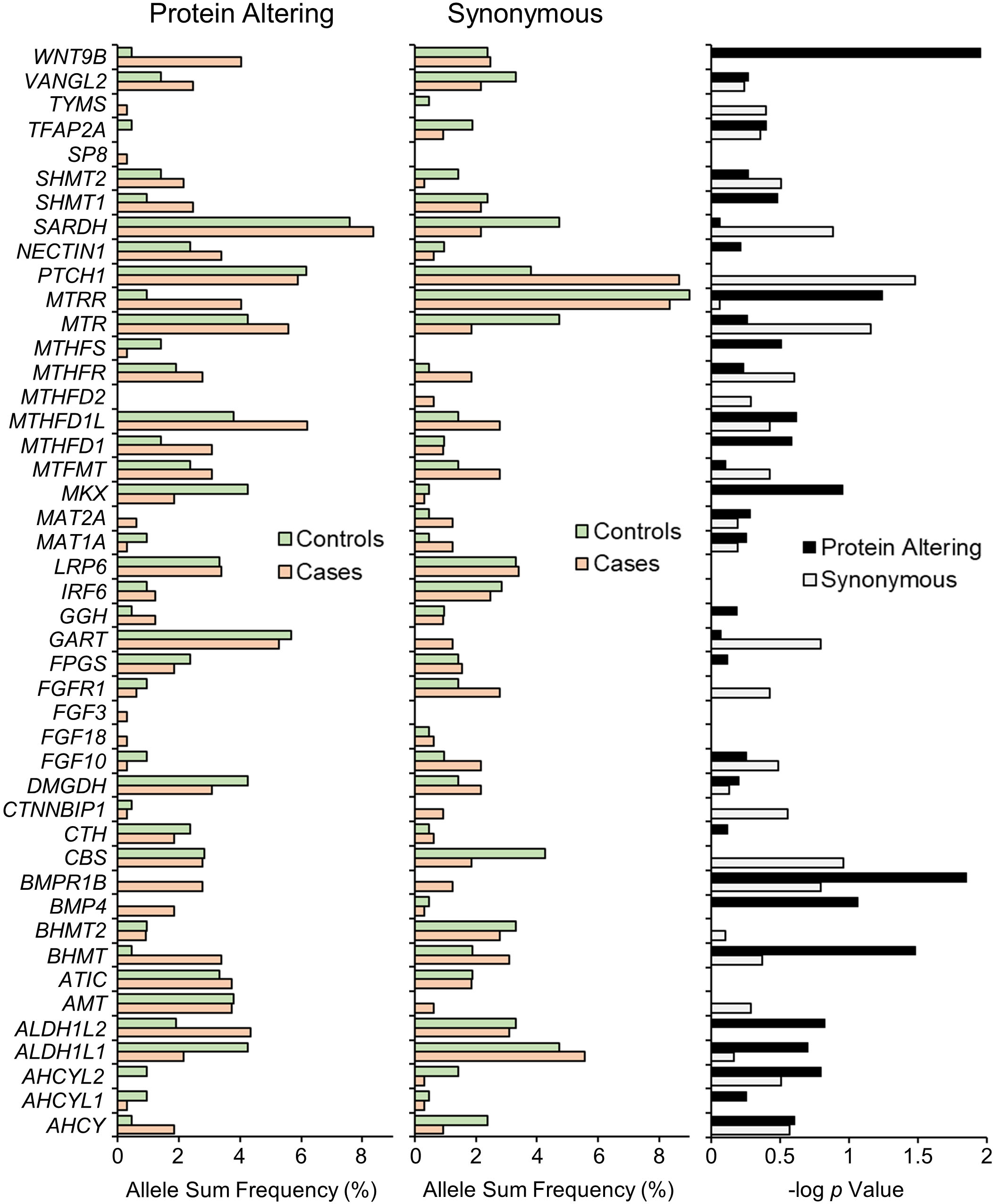
The occurrences of all rare coding region variants (MAF <1.0%) were summed on a gene-by-gene basis to yield the total allele burden for synonymous (left panel) and protein-altering (middle panel) variants. Protein altering variants encompass frameshifts, truncations and missense alleles. Allele Sum Frequency is the carrier frequency (% heterozygote carriers) for each gene. *p* values were calculated (right panel) based on 2×2 Fisher’s exact test of the summed carrier frequency for synonymous and protein-altering changes and expressed as −log *p*.

**Table III.** Genes with Strongest Case Representations of Rare, Protein-Altering Variants.

| Gene | Case Appearances* |  | Control Appearances* |  |
| --- | --- | --- | --- | --- |
|  | Variant | # | Variant | # |
| *BHMT* | D105N | 1 | V122M | 1 |
|  | E136K | 1 |  |  |
|  | P197S | 3 |  |  |
|  | G199S | 2 |  |  |
|  | K400fs | 4 |  |  |
| *BMP4* | D56E | 1 |  |  |
|  | R139H | 1 |  |  |
|  | R162Q | 1 |  |  |
|  | R223H | 1 |  |  |
|  | R226Q | 1 |  |  |
|  | R269Q | 1 |  |  |
| *BMPR1B* | P23S | 1 |  |  |
|  | V67D | 1 |  |  |
|  | V140I | 2 |  |  |
|  | R149W | 1 |  |  |
|  | Q153R | 1 |  |  |
|  | M301V | 1 |  |  |
|  | R371Q | 1 |  |  |
|  | R413S | 1 |  |  |
| *MTRR* | H36Y | 1 |  |  |
|  | N218K | 1 |  |  |
|  | V265G | 1 |  |  |
|  | T286M | 3 |  |  |
|  | I290T | 1 |  |  |
|  | K352N | 1 |  |  |
|  | R386* | 1 |  |  |
|  | R418Q | 1 |  |  |
|  | T490A | 2 | T490A | 1 |
|  | V674A | 1 | V607I | 1 |
| *WNT9B* | A42V | 1 |  |  |
|  | Q47R | 2 |  |  |
|  | R94Q | 2 | R94Q | 1 |
|  | N99S | 1 |  |  |
|  | A126T | 3 |  |  |
|  | K184N | 1 |  |  |
|  | R222H | 1 |  |  |
|  | S244L | 2 |  |  |

\*All variants were seen as heterozygotes; variants shaded in gray were seen in both cases and controls

Using a minor allele frequency cutoff of 5%, rather than 1%, yielded similar folate pathway-wide trends: there were no significant differences in the aggregate burden of non-coding variants (*p* = 0.38) or synonymous variants (*p* = 0.81), but the aggregate burden of protein-altering variants was significantly higher in cases (case aggregate allele frequency = 0.63, control aggregate allele frequency = 0.50; *p* = 0.02). No additional folate pathway genes, besides *BHMT* and *MTRR*, displayed significant case-control differences.

### Individual Gene Analysis

For each of the 49 genes in this study, rare synonymous or protein-altering variants were collapsed by summing and genotype frequencies for this allele burden metric were subject to Fisher’s exact testing (Fig. 1). Consistent with the absence of a signal seen from synonymous variation in the folate pathway, *PTCH1* was the only gene with a statistically significant altered case-control distribution of rare synonymous variants: case carrier frequency (0.20) was approximately 2-fold higher than that in controls (0.12; *p* = 0.03).

In contrast, there were several genes with higher carrier frequency of protein-altering variants in cases with more striking case-control differentials (Fig. 1 and Table III). In addition to the *BHMT* and *MTRR* allele distributions discussed above, there were 13 case appearances (all heterozygotes) of missense alleles in the *WNT9B* gene, versus only one missense occurrence in controls (*p* = 0.01). One missense change, A126T, was found exclusively in 3 cases. Likewise, in the *BMPR1B* gene, which encodes a receptor for BMP (Bone Morphogenetic Protein) secreted ligands, there were 9 occurrences of missense alleles in cases without a single observation in controls (*p* = 0.01). Also noteworthy is the *BMP4* gene, which encodes a specific, secreted Bone Morphogenetic ligand involved in embryonic head development. We observed 6 different missense alleles of *BMP4* in case heterozygotes versus zero protein-altering variants in controls (*p* = 0.09). These last 3 genes were included based on their roles in craniofacial development (and, in some cases, isolated cleft risk [Juriloff et al., 2014; Suzuki et al., 2009; Yang et al., 2017]). Thus, it was interesting to see striking accumulation of rare variation in CLP cases.

### Replication of the case association of *BHMT* K400fs in a second population

As described above, we observed multiple case occurrences of a frameshift (delA) at lysine 400 in *BHMT*. This lesion occurs only 7 amino acids from the C-terminus and is inferred (by nucleotide sequence) to replace these last 7 residues (KQKFKSQ) with 13 different amino acids (NKNSNHSSLDRSYF). Thus, the functional impact of this mutation is difficult to confidently discern. We took the same set of Exon 8-specific resequencing primers used for variant verification (see Methods) to genotype-by-sequence this variant in a second, case-control population (validation population in Table I). We observed this variant as a heterozygote 3 additional times in the validation population: 2 additional CLP cases (one Hispanic, one non-Hispanic, White) and one control (non-Hispanic, White). We did not observe this variant in the cleft palate only (CPO) subset of this population (N = 157).

## DISCUSSION

In this study we took a candidate gene approach to investigate the potential contribution of rare coding variants to CLP risk, with the reasoning that at least some of the genetic risk not yet identified in common allele studies may reside in rare variation. The contribution of rare, coding mutations in CLP risk has been previously suggested in multiple sequencing studies [Suzuki et al., 2009; Leslie and Murray, 2013; Al Chawa et al., 2014], though this has only recently begun to be systematically addressed [Aylward et al., 2016; Leslie et al., 2017]. We focused on 49 candidate genes with an emphasis on nearly complete coverage of folate metabolism (N = 30 genes), since rare folate pathway variants have not been explored for CLP risk, particularly when considered in aggregate across the pathway or within related genesets. We queried 19 additional genes unrelated to folate metabolism, that have been implicated in disease risk by vertebrate models of clefting (e.g. *WNT9B*) or craniofacial development, or by human genetic association studies (e.g. *BMP4*, *IRF6*). From a catalog of the full spectrum of allelic variation within the coding regions of this target-enriched gene set, we found two lines of evidence for the contribution of rare variants to CLP risk: 1) cases displayed an overabundance of putative loss-of-function alleles at these loci, and 2) several mechanistically relevant genes showed similar case over-representation for protein-altering variants, in general.

### Folate pathway variation: Methionine cycle genes

We sequenced the majority of folate/one-carbon pathway genes, which enabled us to interpret rare variation in the context of pathway function. This approach was useful in identifying folate-related risk signatures for neural tube defects [Marini et al., 2011]. While we did not see any pathway-wide trends in allele burdens or distributions that significantly differentiated CLP cases from controls, we did observe 2 genes directly involved in methionine biosynthesis with a considerable over-abundance of protein altering variants in cases: *BHMT* and *MTRR* (Table III). Betaine-homocysteine methyltransferase (BHMT) and Methionine synthase reductase (MTRR) are both involved in the methylation of homocysteine to yield methionine. They operate in different pathways, however, and while BHMT utilizes methyl groups generated from choline catabolism, the reaction involving MTRR utilizes 5-methyl-tetrahydrofolate.

The CLP-association of rare variants at *BHMT* is intriguing for two reasons: First, this locus, which is part of the metabolically-related *BHMT/BHMT2/DMGDH* gene cluster on chromosome 5, is one of the most consistently associated loci in one-carbon pathway gene-CLP association studies [reviewed in Marini et al., 2016]. However, while the locus is often associated, the regional SNPs that drive the association can be different, as can the directionality of risk (i.e., whether odds ratios are < or > 1). We postulated that these phenomena may reflect the contribution of underlying rare variants at this locus (which may occur on the haplotype of the major or minor allele of a nearby common variant) as has been previously suggested for clefts [Leslie and Murray 2013] and as has been seen for regions highlighted by GWAS studies [Leslie et al., 2015]. The distribution of rare variants at *BHMT* is also intriguing because of the appearance of a frameshift mutation (K400fs) that was observed in multiple CLP cases (discussed below).

The increased mutation burden in these 2 genes brings attention to homocysteine clearance as relevant for modifying CLP risk, which has been previously suggested [Wong et al., 1999; Kumari et al., 2013]. Furthermore, interference of methionine synthesis suggests perturbations in cellular methylation reactions (via S-adenosylmethionine levels), which may manifest in DNA methylation changes associated with CLP, as has been observed [Sharp et al., 2016; Alvizi et al., 2017].

### Putative loss-of-function alleles

Within this target gene set, CLP cases displayed a significantly greater frequency of indel frameshift and truncation alleles (*p* = 0.01; Table II). One of the most intriguing alleles is the K400fs deletion allele of *BHMT*, which was observed in four cases (both Hispanic and non-Hispanic, white) but not in controls in the discovery population. A second, validation population revealed 2 additional CLP case occurrences and one control occurrence. Although the functional impact of this allele is yet to be determined, its observance in multiple cases is noteworthy particularly because global population frequencies of this variant are estimated at 2.2E-04 in the Genome Aggregation Database (www.gnomad.broadinstitute.org), substantially lower than our overall case frequency (MAF = 0.004 in all cases queried). Although this variant is described as low-confidence in this database, the signal was strong in our data and all instances clearly replicated in verification Sanger sequencing.

In addition to *BHMT*, we also observed case-associated truncations in several other genes relevant to methyl donor production (*MTRR* R386*, *DMGDH* W495* and *SARDH* R457*, R514*). Based on protein lengths (MTRR 698aa, DMGDH 866aa, SARDH 918aa), these changes are likely to be consequential. BHMT, DMGDH (Dimethyl-glycine dehydrogenase) and SARDH (Sarcosine dehydrogenase) are sequential enzymatic steps in the catabolism of choline to glycine, which generates methyl donors for cellular methylation reactions as noted above. Interestingly, increased intake of choline in the maternal diet has been associated with decreased CLP risk [Shaw et al., 2006].

Outside of folate/one-carbon pathway genes, 2 case truncations were observed in genes previously implicated in orofacial cleft risk: *IRF6* Q112* and *SP8* S261*. Significant associations of *IRF6* with CLP have been reported in multiple populations [Beaty et al., 2016] and may account for up to 12% of the genetic component of CLP in some populations [Zucchero et al., 2004]. SP8 is a transcription factor in which mutations display severe craniofacial malformations including cleft palate [Kasberg et al., 2013].

### Genes unrelated to the folate metabolism

Three additional genes involved in craniofacial development also displayed striking case-overrepresentation of missense variants: *BMP4*, *BMPR1B* and *WNT9B* (Table III). WNT9B and BMP4 are secreted signaling proteins regulating critical interacting signaling pathways in craniofacial development [Alexander et al., 2014]. BMPR1B encodes a receptor for BMP4 and other bone morphogens. Significantly, all 3 genes have been implicated in orofacial cleft risk.

*BMP4* is one of the most promising candidate genes for CLP, with evidence shown in animal experiments [e.g. Liu et al., 2005] and association studies [e.g. Chen et al., 2014]. Furthermore, a gene sequencing study [Suzuki et al., 2009] demonstrated that rare *BMP4* coding mutations were enriched in CLP cases at similar frequencies to those shown here. Only the R162Q case-specific nonsynonymous mutation was identified in both studies.

Although rare coding variants in *WNT9B* or *BMPR1B* have yet to be reported as contributing to CLP risk, these genes have been implicated by other means. *WNT9B* is at a locus identified in a genome-wide association study [Yu et al., 2017] and a murine model demonstrated that transcriptional repression of Wnt9b via epigenetic mechanisms leads to the development of cleft lip in mouse embryos [Juriloff et al., 2014]. *BMPR1B* is at a locus associated with CLP by linkage analysis [Schultz et al., 2004] and has been identified as a genetic cause of Pierre Robin syndrome, in which cleft palate is a prominent component [Yang et al., 2017].

### Rare versus private mutations

Of all the variants described in Tables II and III, most have been seen before in the population, as measured by their annotation in the Genome Aggregation Database (www.gnomad.broadinstitute.org). Novel variants in this study are restricted to 2 missense variants in *BMPR1B* (P23S and V67D), and 3 putative loss-of-function truncations (*DMGDH* W495*, *IRF6* Q112*, *SP8* S261*). The frameshift allele in *GGH* (insT at L190) was previously observed once in over 250,000 alleles. All others have multiple occurrences, though at significantly lower frequencies in the general population than seen here in CLP cases. However, most of our observations are singletons (with a notable exception being the K400fs allele of *BHMT*), precluding true case frequency estimates.

If rare alleles such as these contribute to CLP risk, it is likely that they will be uncovered only by sequencing/discovery approaches rather than array-based genotyping, even for large array sizes. Furthermore, the degree to which risk loci identified by traditional association studies will reveal loci with underlying rare variant burdens (as we have suggested for the *BHMT*/*BHMT2*/*DMGDH* gene cluster) is still unknown. For example, Leslie et al.,[2015] failed to find convincing evidence of CLP-related rare allele burdens at GWAS loci in 1500 trios. It may be that the underlying genetic heterogeneity for CLP risk is sufficiently complex that many such burdened loci do not display genome-wide significance in traditional association studies. Indeed, nearly all variants discussed here were heterozygous and none co-occurred with any others. Thus, if they are involved in CLP risk, they must be part of a more complex etiology.

### Strengths/Limitations

Strengths of this study lie in the case-control design, which allows for a broader sampling of rare alleles than family-based studies, and the assurance of accuracy in the sequencing data, which is essential for rare variant discovery. Although CLP risk loci identified in this study are consistent with previous studies and with biological mechanisms, the study is limited by both sample size and the ability to draw reliable associations from rare variants. Thus, statistical measures used here are not a true measure of association but of highlighting genes with differential case burdens. Therefore, replication of these findings in additional, larger populations is needed to afford significance of the potential associations observed here. Another limitation is that the functional impact of variants is unknown. All mutations were collapsed and weighted equally. Attempts to weigh variants based on predicted functional impact yielded inconsistent classifications, thus we await empirical determination of functional impact to enable more robust mechanistic conclusions with respect to phenotype. Nevertheless, these results are consistent with several reports implicating rare coding variants in CLP risk. The clustering of risk genes reinforces focus on one-carbon metabolism and the BMP/WNT signaling pathways.

## Acknowledgments

This project was supported by NIH R01 HD074695 (N.J.M), CDC 6U01-DD-000489 (G.M.S.) and 1U01-DD-0006982 (G.M.S.). This work used the Vincent J. Coates Genomics Sequencing Laboratory at UC Berkeley. We thank the California Department of Public Health Maternal Child and Adolescent Health Division for providing data. The findings and conclusions in this report are those of the authors and do not necessarily represent the official position of the California Department of Public Health. The authors declare no conflicts of interest.

## Supplementary

Table S1. Gene/exon annotations; Sequencing coverage/failures.

Figure S1. Target exon coverage in cases vs. controls.

Table S2. Genotype concordance between exon sequencing and TaqMan allelic discrimination assay.

Table S3. Rare variant verification by resequencing gDNA prior to whole genome amplification.

Table S4. Annotations and genotype distributions for 773 variants scored in this study.

