## Supplementary figures and images for "Accumulation of Rare Coding Variants in Genes Implicated in Risk of Human Cleft Lip with or without Cleft Palate"

### Figure S1

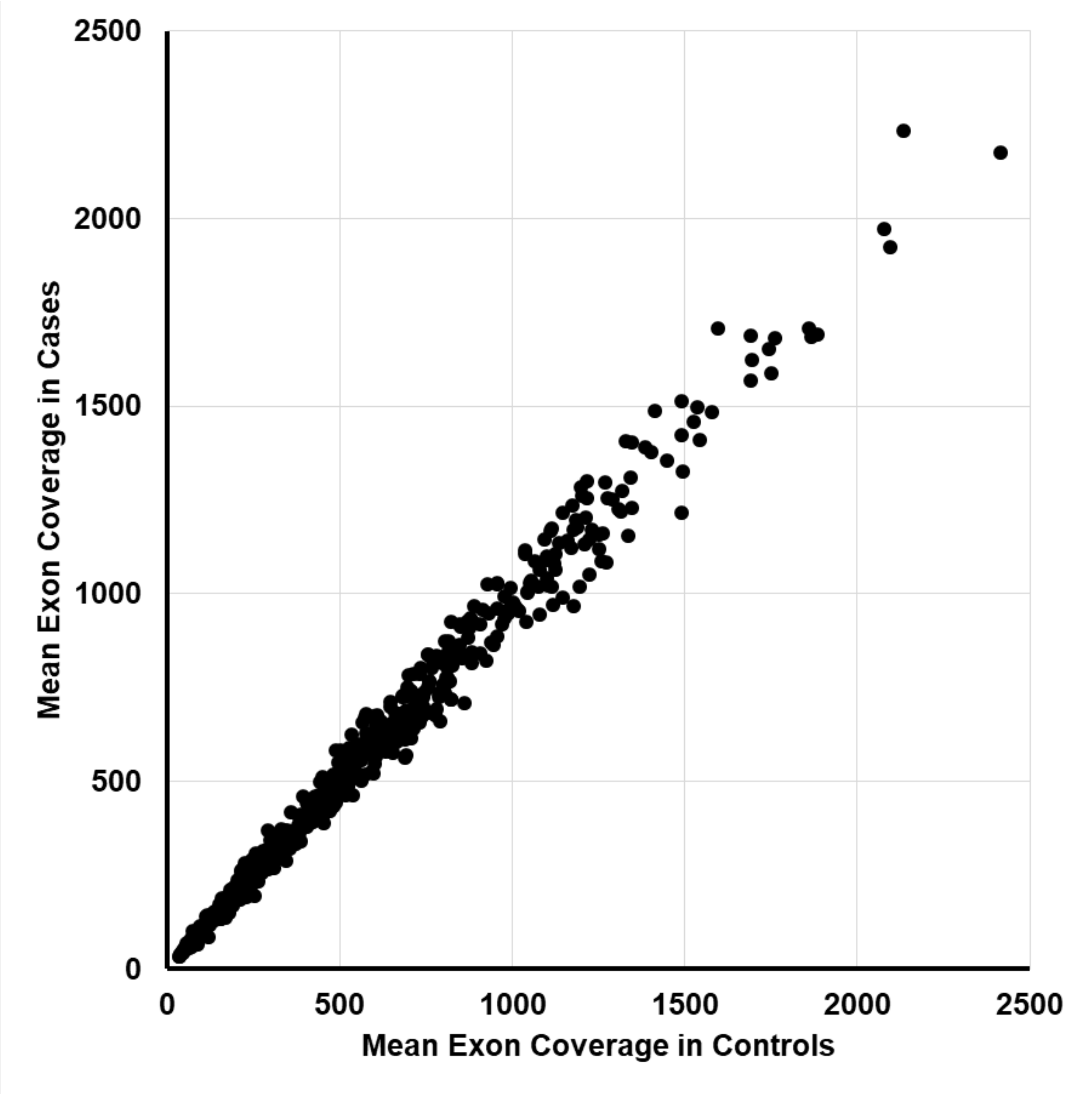


Figure S1. The mean exon coverage in cases versus controls for the 532 exons that passed QC filtering.
