## Supplementary material for "Accumulation of Rare Coding Variants in Genes Implicated in Risk of Human Cleft Lip with or without Cleft Palate": Table S1

**Table S1**. Annotation of the coding regions sequenced in this study by gene/exon.
First and/or last exons in each gene highligted in gray contain 5'-UTR and 3'-UTR sequences.
Exons discarded due to poor quality are indicated in the last column.
* Exon coordinates are according the Genomic Locus RefSeq ID (NG#)
** 90% of case or control samples had mean exon coverage greater than value indicated
*** % case or control samples with >30X coverage across entire CDS region of exon

| Gene | Genomic Locus RefSeq ID | mRNA RefSeq ID | CCDS RefSeq ID | Exon | Exon Coordinate 1* | Exon Coordinate 2* | mRNA Length | CDS length | CTRL Coverage of CDS regions (SAMPLE 10%ILE)** | CASE Coverage of CDS regions (SAMPLE 10%ILE)** | % CTRL SAMPLES with CDS region coverage >30X *** | % CASE SAMPLES with CDS region coverage >30X *** | QC COMMENT |
|---|---|---|---|---|---|---|---|---|---|---|---|---|---|
| AHCY | NG_012630 | NM_001161766.1 | CCDS54457.1 | 2 | 21218 | 21408 | 191 | 135 | 359 | 417 | 100 | 98 | |
| | | | | 3 | 22647 | 22722 | 76 | 76 | 273 | 256 | 100 | 98 | |
| | | | | 4 | 24296 | 24445 | 150 | 150 | 248 | 262 | 100 | 99 | |
| | | | | 5 | 25272 | 25384 | 113 | 113 | 437 | 454 | 100 | 100 | |
| | | | | 6 | 25865 | 26072 | 208 | 208 | 665 | 668 | 100 | 99 | |
| | | | | 7 | 26165 | 26252 | 88 | 88 | 673 | 680 | 100 | 100 | |
| | | | | 8 | 26354 | 26471 | 118 | 118 | 742 | 725 | 100 | 100 | |
| | | | | 9 | 31169 | 31363 | 195 | 195 | 456 | 487 | 100 | 99 | |
| | | | | 10 | 35638 | 36538 | 901 | 132 | 162 | 160 | 100 | 98 | |
| AHCYL1 | NG_029182 | NM_006621.5 | CCDS818.1 | 1 | 5001 | 5409 | 409 | 120 | 12 | 14 | 69 | 63 | poor quality/discard |
| | | | | 2 | 29271 | 29382 | 112 | 112 | 970 | 918 | 100 | 100 | |
| | | | | 3 | 31451 | 31594 | 144 | 144 | 787 | 724 | 100 | 100 | |
| | | | | 4 | 32599 | 32699 | 101 | 101 | 798 | 830 | 100 | 100 | |
| | | | | 5 | 33134 | 33236 | 103 | 103 | 715 | 640 | 100 | 100 | |
| | | | | 6 | 35000 | 35094 | 95 | 95 | 484 | 479 | 100 | 100 | |
| | | | | 7 | 35662 | 35768 | 107 | 107 | 503 | 475 | 100 | 100 | |
| | | | | 8 | 36581 | 36697 | 117 | 117 | 620 | 612 | 100 | 100 | |
| | | | | 9 | 36904 | 36967 | 64 | 64 | 676 | 662 | 100 | 100 | |
| | | | | 10 | 37732 | 37820 | 89 | 89 | 809 | 776 | 100 | 100 | |
| | | | | 11 | 38183 | 38253 | 71 | 71 | 740 | 673 | 100 | 100 | |
| | | | | 12 | 38610 | 38704 | 95 | 95 | 632 | 593 | 100 | 100 | |
| | | | | 13 | 38789 | 38887 | 99 | 99 | 528 | 523 | 100 | 100 | |
| | | | | 14 | 39288 | 39356 | 69 | 69 | 786 | 736 | 100 | 100 | |
| | | | | 15 | 39785 | 39863 | 79 | 79 | 637 | 601 | 100 | 100 | |
| | | | | 16 | 40968 | 41088 | 121 | 121 | 717 | 677 | 100 | 100 | |
| | | | | 17 | 41911 | 43979 | 2069 | 7 | 91 | 80 | 99 | 97 | |
| AHCYL2 | NG_029180 | NM_015328.3 | CCDS5812.1 | 1 | 5001 | 5426 | 426 | 363 | 20 | 23 | 78 | 83 | poor quality/discard |
| | | | | 2 | 159625 | 159736 | 112 | 112 | 1527 | 1458 | 100 | 100 | |
| | | | | 3 | 169043 | 169186 | 144 | 144 | 660 | 682 | 100 | 100 | |
| | | | | 4 | 169621 | 169721 | 101 | 101 | 755 | 838 | 100 | 100 | |
| | | | | 5 | 177209 | 177311 | 103 | 103 | 314 | 319 | 100 | 99 | |
| | | | | 6 | 180277 | 180371 | 95 | 95 | 292 | 367 | 100 | 100 | |
| | | | | 7 | 183366 | 183472 | 107 | 107 | 504 | 583 | 100 | 100 | |
| | | | | 8 | 185084 | 185200 | 117 | 117 | 817 | 849 | 100 | 100 | |
| | | | | 9 | 185823 | 185886 | 64 | 64 | 620 | 659 | 100 | 100 | |
| | | | | 10 | 186365 | 186453 | 89 | 89 | 639 | 605 | 100 | 100 | |
| | | | | 11 | 189463 | 189533 | 71 | 71 | 1271 | 1298 | 100 | 100 | |
| | | | | 12 | 193581 | 193675 | 95 | 95 | 703 | 741 | 100 | 100 | |
| | | | | 13 | 202827 | 202925 | 99 | 99 | 1123 | 1079 | 100 | 100 | |
| | | | | 14 | 204861 | 204929 | 69 | 69 | 1044 | 1002 | 100 | 100 | |
| | | | | 15 | 205050 | 205128 | 79 | 79 | 1147 | 989 | 100 | 100 | |
| | | | | 16 | 206430 | 206550 | 121 | 121 | 1119 | 969 | 100 | 100 | |
| | | | | 17 | 207026 | 210198 | 3173 | 7 | 100 | 89 | 100 | 98 | |
| ALDH1L1 | NG_012260 | NM_012190.3 | CCDS3034.1 | 2 | 24641 | 24790 | 150 | 127 | 564 | 506 | 100 | 100 | |
| | | | | 3 | 27004 | 27238 | 235 | 235 | 691 | 570 | 100 | 100 | |
| | | | | 4 | 28135 | 28300 | 166 | 166 | 563 | 503 | 100 | 100 | |
| | | | | 5 | 30140 | 30241 | 102 | 102 | 490 | 453 | 100 | 100 | |
| | | | | 6 | 31000 | 31089 | 90 | 90 | 419 | 402 | 100 | 100 | |
| | | | | 7 | 32062 | 32199 | 138 | 138 | 458 | 417 | 100 | 100 | |
| | | | | 8 | 35112 | 35237 | 126 | 126 | 419 | 394 | 100 | 100 | |
| | | | | 9 | 38717 | 38808 | 92 | 92 | 597 | 522 | 100 | 100 | |
| | | | | 10 | 47683 | 47830 | 148 | 148 | 408 | 396 | 100 | 100 | |
| | | | | 11 | 48760 | 48879 | 120 | 120 | 388 | 341 | 100 | 100 | |
| | | | | 12 | 49981 | 50108 | 128 | 128 | 346 | 287 | 100 | 98 | |
| | | | | 13 | 54109 | 54259 | 151 | 151 | 493 | 483 | 100 | 99 | |
| | | | | 14 | 55350 | 55420 | 71 | 71 | 508 | 465 | 100 | 99 | |
| | | | | 15 | 59922 | 60027 | 106 | 106 | 697 | 681 | 100 | 100 | |
| | | | | 16 | 61192 | 61279 | 88 | 88 | 525 | 508 | 100 | 100 | |
| | | | | 17 | 67545 | 67638 | 94 | 94 | 203 | 191 | 100 | 98 | |
| | | | | 18 | 70987 | 71086 | 100 | 100 | 428 | 429 | 100 | 100 | |
| | | | | 19 | 72763 | 72861 | 99 | 99 | 407 | 391 | 100 | 99 | |
| | | | | 20 | 75534 | 75699 | 166 | 166 | 649 | 580 | 100 | 100 | |
| | | | | 21 | 78397 | 78502 | 106 | 106 | 645 | 638 | 100 | 100 | |
| | | | | 22 | 79718 | 79917 | 200 | 200 | 459 | 454 | 100 | 100 | |
| | | | | 23 | 81797 | 82082 | 286 | 56 | 225 | 230 | 100 | 97 | |
| ALDH1L2 | NG_012748 | NM_001034173.3 | CCDS31891.1 | 1 | 5001 | 5175 | 175 | 48 | 3 | 4 | 42 | 44 | poor quality/discard |
| | | | | 2 | 15559 | 15703 | 145 | 145 | 781 | 692 | 100 | 100 | |
| | | | | 3 | 18760 | 18994 | 235 | 235 | 1080 | 1063 | 100 | 100 | |
| | | | | 4 | 20680 | 20845 | 166 | 166 | 884 | 815 | 100 | 100 | |
| | | | | 5 | 22895 | 22996 | 102 | 102 | 1197 | 1019 | 100 | 100 | |
| | | | | 6 | 24208 | 24297 | 90 | 90 | 804 | 736 | 100 | 100 | |
| | | | | 7 | 26542 | 26676 | 135 | 135 | 716 | 677 | 100 | 100 | |
| | | | | 8 | 27812 | 27937 | 126 | 126 | 707 | 615 | 100 | 100 | |
| | | | | 9 | 28492 | 28583 | 92 | 92 | 687 | 563 | 100 | 100 | |
| | | | | 10 | 31344 | 31491 | 148 | 148 | 776 | 676 | 100 | 100 | |
| | | | | 11 | 36633 | 36752 | 120 | 120 | 472 | 421 | 100 | 100 | |
| | | | | 12 | 37347 | 37474 | 128 | 128 | 323 | 326 | 100 | 100 | |
| | | | | 13 | 39506 | 39656 | 151 | 151 | 529 | 499 | 100 | 100 | |
| | | | | 14 | 42595 | 42665 | 71 | 71 | 416 | 393 | 100 | 100 | |
| | | | | 15 | 42749 | 42854 | 106 | 106 | 490 | 445 | 100 | 100 | |
| | | | | 16 | 48870 | 48957 | 88 | 88 | 791 | 659 | 100 | 100 | |
| | | | | 17 | 49758 | 49851 | 94 | 94 | 592 | 523 | 100 | 100 | |
| | | | | 18 | 51362 | 51461 | 100 | 100 | 715 | 647 | 100 | 100 | |

| | | | | | | | | | | | | | |
|---|---|---|---|---|---|---|---|---|---|---|---|---|---|
| | | | | 19 | 55166 | 55264 | 99 | 99 | 938 | 869 | 100 | 100 | |
| | | | | 20 | 57630 | 57795 | 166 | 166 | 859 | 709 | 100 | 100 | |
| | | | | 21 | 59135 | 59240 | 106 | 106 | 1272 | 1083 | 100 | 100 | |
| | | | | 22 | 62820 | 63019 | 200 | 200 | 2416 | 2176 | 100 | 100 | |
| | | | | 23 | 65085 | 69780 | 4696 | 56 | 251 | 261 | 100 | 99 | |
| AMT | NG_015986 | NM_000481.3 | CCDS2797.1 | 1 | 5001 | 5318 | 318 | 90 | 78 | 71 | 99 | 94 | |
| | | | | 2 | 5408 | 5575 | 168 | 168 | 154 | 134 | 100 | 97 | |
| | | | | 3 | 6107 | 6187 | 81 | 81 | 223 | 239 | 100 | 98 | |
| | | | | 4 | 7337 | 7468 | 132 | 132 | 176 | 196 | 100 | 97 | |
| | | | | 5 | 7891 | 7969 | 79 | 79 | 220 | 216 | 100 | 99 | |
| | | | | 6 | 8274 | 8419 | 146 | 146 | 196 | 182 | 100 | 97 | |
| | | | | 7 | 8528 | 8708 | 181 | 181 | 265 | 235 | 100 | 97 | |
| | | | | 8 | 9706 | 9861 | 156 | 156 | 330 | 306 | 100 | 98 | |
| | | | | 9 | 9961 | 10901 | 941 | 179 | 189 | 197 | 100 | 98 | |
| ATIC | NG_013002 | NM_004044.6 | CCDS2398.1 | 1 | 5001 | 5206 | 206 | 19 | 9 | 12 | 62 | 65 | poor quality/discard |
| | | | | 2 | 5543 | 5669 | 127 | 127 | 361 | 352 | 100 | 100 | |
| | | | | 3 | 11202 | 11278 | 77 | 77 | 547 | 519 | 100 | 100 | |
| | | | | 4 | 12710 | 12776 | 67 | 67 | 604 | 674 | 100 | 100 | |
| | | | | 5 | 18286 | 18374 | 89 | 89 | 765 | 802 | 100 | 100 | |
| | | | | 6 | 19032 | 19183 | 152 | 152 | 846 | 865 | 100 | 100 | |
| | | | | 7 | 19867 | 20023 | 157 | 157 | 1329 | 1406 | 100 | 100 | |
| | | | | 8 | 25427 | 25552 | 126 | 126 | 647 | 700 | 100 | 100 | |
| | | | | 9 | 26395 | 26502 | 108 | 108 | 956 | 885 | 100 | 100 | |
| | | | | 10 | 27964 | 28049 | 86 | 86 | 888 | 968 | 100 | 100 | |
| | | | | 11 | 29080 | 29169 | 90 | 90 | 834 | 835 | 100 | 100 | |
| | | | | 12 | 31824 | 31952 | 129 | 129 | 1100 | 1042 | 100 | 100 | |
| | | | | 13 | 37824 | 37916 | 93 | 93 | 1041 | 926 | 100 | 100 | |
| | | | | 14 | 39804 | 39986 | 183 | 183 | 1223 | 1052 | 100 | 100 | |
| | | | | 15 | 42139 | 42294 | 156 | 156 | 1263 | 1162 | 100 | 100 | |
| | | | | 16 | 42581 | 42818 | 238 | 120 | 1252 | 1156 | 100 | 100 | |
| BHMT | NG_029156 | NM_001713.2 | CCDS4046.1 | 1 | 5001 | 5138 | 138 | 33 | 227 | 215 | 100 | 98 | |
| | | | | 2 | 8987 | 9119 | 133 | 133 | 525 | 485 | 100 | 100 | |
| | | | | 3 | 12479 | 12597 | 119 | 119 | 461 | 468 | 100 | 100 | |
| | | | | 4 | 13570 | 13761 | 192 | 192 | 452 | 390 | 100 | 100 | |
| | | | | 5 | 14438 | 14585 | 148 | 148 | 340 | 349 | 100 | 100 | |
| | | | | 6 | 19266 | 19448 | 183 | 183 | 731 | 657 | 100 | 100 | |
| | | | | 7 | 20975 | 21203 | 229 | 229 | 494 | 550 | 100 | 100 | |
| | | | | 8 | 24153 | 25510 | 1358 | 184 | 383 | 363 | 100 | 100 | |
| BHMT2 | NG_029157 | NM_017614.4 | CCDS4045.1 | 1 | 5001 | 5092 | 92 | 33 | 0 | 0 | 0 | 0 | exon fail/discard |
| | | | | 2 | 12757 | 12889 | 133 | 133 | 1101 | 1099 | 100 | 100 | |
| | | | | 3 | 14646 | 14737 | 92 | 92 | 1447 | 1355 | 100 | 100 | |
| | | | | 4 | 15964 | 16155 | 192 | 192 | 1866 | 1683 | 100 | 100 | |
| | | | | 5 | 18086 | 18233 | 148 | 148 | 1496 | 1325 | 100 | 100 | |
| | | | | 6 | 18469 | 18651 | 183 | 183 | 1176 | 1236 | 100 | 100 | |
| | | | | 7 | 18905 | 19133 | 229 | 229 | 1218 | 1255 | 100 | 100 | |
| | | | | 8 | 23770 | 25351 | 1582 | 82 | 356 | 319 | 100 | 100 | |
| BMP4 | NG_009215 | NM_001202.5 | CCDS9715.1 | 3 | 9608 | 9984 | 377 | 370 | 611 | 581 | 100 | 100 | |
| | | | | 4 | 10949 | 12101 | 1153 | 857 | 605 | 613 | 100 | 100 | |
| BMPR1B | NG_009245 | NM_001203.2 | CCDS3642.1 | 4 | 351432 | 351591 | 160 | 143 | 907 | 840 | 100 | 100 | |
| | | | | 5 | 361744 | 361846 | 103 | 103 | 1126 | 1107 | 100 | 100 | |
| | | | | 6 | 362709 | 362811 | 103 | 103 | 1346 | 1228 | 100 | 100 | |
| | | | | 7 | 370834 | 370930 | 97 | 97 | 1216 | 1300 | 100 | 100 | |
| | | | | 8 | 372007 | 372145 | 139 | 139 | 1763 | 1680 | 100 | 100 | |
| | | | | 9 | 376886 | 377078 | 193 | 193 | 1316 | 1220 | 100 | 100 | |
| | | | | 10 | 378239 | 378536 | 298 | 298 | 2094 | 1925 | 100 | 100 | |
| | | | | 11 | 395772 | 395947 | 176 | 176 | 579 | 592 | 100 | 100 | |
| | | | | 12 | 399667 | 399797 | 131 | 131 | 508 | 523 | 100 | 100 | |
| | | | | 13 | 401572 | 405474 | 3903 | 126 | 179 | 151 | 99 | 98 | |
| CBS | NG_008938 | NM_000071.2 | CCDS13693.1 | 3 | 8730 | 8946 | 217 | 209 | 66 | 61 | 97 | 94 | |
| | | | | 4 | 12316 | 12422 | 107 | 107 | 102 | 103 | 100 | 94 | |
| | | | | 5 | 14554 | 14688 | 135 | 135 | 259 | 237 | 100 | 97 | |
| | | | | 6 | 15236 | 15315 | 80 | 80 | 182 | 210 | 99 | 97 | |
| | | | | 7 | 15410 | 15544 | 135 | 135 | 235 | 260 | 100 | 97 | |
| | | | | 8 | 15659 | 15728 | 70 | 70 | 207 | 230 | 100 | 97 | |
| | | | | 9 | 16940 | 17031 | 92 | 92 | 242 | 219 | 100 | 97 | |
| | | | | 10 | 17853 | 17978 | 126 | 126 | 169 | 136 | 100 | 95 | |
| | | | | 11 | 18536 | 18620 | 85 | 85 | 215 | 208 | 100 | 97 | |
| | | | | 12 | 20385 | 20490 | 106 | 106 | 112 | 138 | 98 | 97 | |
| | | | | 13 | 21628 | 21705 | 78 | 78 | 123 | 118 | 99 | 95 | |
| | | | | 14 | 21963 | 22097 | 135 | 135 | 95 | 113 | 99 | 94 | |
| | | | | 15 | 22678 | 22786 | 109 | 109 | 163 | 168 | 100 | 97 | |
| | | | | 16 | 24044 | 24128 | 85 | 85 | 110 | 119 | 99 | 95 | |
| | | | | 17 | 26948 | 27740 | 793 | 104 | 40 | 38 | 98 | 94 | |
| CTH | NG_008041 | NM_001902 | CCDS650.1 | 1 | 4947 | 5312 | 366 | 168 | 303 | 317 | 100 | 99 | |
| | | | | 2 | 9685 | 9766 | 82 | 82 | 692 | 690 | 100 | 100 | |
| | | | | 3 | 11666 | 11761 | 96 | 96 | 680 | 727 | 100 | 100 | |
| | | | | 4 | 15295 | 15404 | 110 | 110 | 611 | 609 | 100 | 100 | |
| | | | | 5 | 18014 | 18145 | 132 | 132 | 790 | 825 | 100 | 100 | |
| | | | | 6 | 23523 | 23580 | 58 | 58 | 517 | 568 | 100 | 100 | |
| | | | | 7 | 24046 | 24123 | 78 | 78 | 213 | 258 | 100 | 100 | |
| | | | | 8 | 25812 | 25964 | 153 | 153 | 507 | 540 | 100 | 100 | |
| | | | | 9 | 27557 | 27678 | 122 | 122 | 566 | 658 | 100 | 100 | |
| | | | | 10 | 28854 | 28906 | 53 | 53 | 473 | 507 | 100 | 100 | |
| | | | | 11 | 32417 | 32555 | 139 | 139 | 515 | 533 | 100 | 100 | |
| | | | | 12 | 32830 | 33580 | 751 | 27 | 190 | 203 | 100 | 100 | |
| CTNNBIP1 | NG_029165 | NM_020248.2 | CCDS106.1 | 4 | 43171 | 43290 | 120 | 96 | 535 | 625 | 100 | 100 | |
| | | | | 5 | 43982 | 44072 | 91 | 91 | 193 | 192 | 100 | 97 | |
| | | | | 6 | 64483 | 66983 | 2501 | 59 | 50 | 50 | 97 | 95 | |
| DHFR | NG_023304 | NM_000791.3 | CCDS47240.1 | 1 | 5001 | 5578 | 578 | 86 | 212 | 194 | 100 | 97 | |
| | | | | 2 | 5925 | 5974 | 50 | 50 | 1006 | 970 | 100 | 100 | |
| | | | | 3 | 10488 | 10593 | 106 | 106 | 957 | 1028 | 100 | 100 | |
| | | | | 4 | 21973 | 22099 | 127 | 127 | 539 | 531 | 100 | 100 | |
| | | | | 5 | 25990 | 26105 | 116 | 116 | 604 | 567 | 100 | 100 | |
| | | | | 6 | 30817 | 33756 | 2940 | 79 | 116 | 118 | 99 | 98 | |
| DMGDH | NG_012164 | NM_013391.3 | CCDS4044.1 | 1 | 4953 | 5107 | 155 | 101 | 0 | 0 | 0 | 0 | exon fail/discard |
| | | | | 2 | 10840 | 11014 | 175 | 175 | 743 | 698 | 100 | 100 | |
| | | | | 3 | 18719 | 18817 | 99 | 99 | 1490 | 1514 | 100 | 100 | |

| | | | | | | | | | | | | | |
|---|---|---|---|---|---|---|---|---|---|---|---|---|---|
| | | | | 4 | 20279 | 20443 | 165 | 165 | 1161 | 1142 | 100 | 100 | |
| | | | | 5 | 23136 | 23340 | 205 | 205 | 1257 | 1087 | 100 | 100 | |
| | | | | 6 | 30075 | 30323 | 249 | 249 | 719 | 662 | 100 | 100 | |
| | | | | 7 | 32146 | 32344 | 199 | 199 | 1104 | 1022 | 100 | 100 | |
| | | | | 8 | 41219 | 41388 | 170 | 170 | 686 | 656 | 100 | 100 | |
| | | | | 9 | 41787 | 41940 | 154 | 154 | 725 | 669 | 100 | 100 | |
| | | | | 10 | 43629 | 43794 | 166 | 166 | 633 | 580 | 100 | 100 | |
| | | | | 11 | 44593 | 44723 | 131 | 131 | 572 | 518 | 100 | 100 | |
| | | | | 12 | 45977 | 46194 | 218 | 218 | 602 | 547 | 100 | 100 | |
| | | | | 13 | 48046 | 48203 | 158 | 158 | 819 | 768 | 100 | 100 | |
| | | | | 14 | 50297 | 50356 | 60 | 60 | 711 | 687 | 100 | 100 | |
| | | | | 15 | 69220 | 69354 | 135 | 135 | 1231 | 1171 | 100 | 100 | |
| | | | | 16 | 76330 | 77063 | 734 | 216 | 553 | 513 | 100 | 100 | |
| FGFR1 | NG_007729 | NM_023110.2 | CCDS6107.2 | 2 | 16301 | 16479 | 179 | 91 | 213 | 224 | 100 | 98 | |
| | | | | 3 | 43887 | 44153 | 267 | 267 | 205 | 236 | 99 | 97 | |
| | | | | 4 | 45400 | 45489 | 90 | 90 | 5 | 4 | 19 | 13 | exon fail/discard |
| | | | | 5 | 45742 | 45914 | 173 | 173 | 518 | 484 | 100 | 100 | |
| | | | | 6 | 47590 | 47713 | 124 | 124 | 593 | 599 | 100 | 100 | |
| | | | | 7 | 49136 | 49326 | 191 | 191 | 651 | 580 | 100 | 100 | |
| | | | | 8 | 51894 | 52038 | 145 | 145 | 502 | 536 | 100 | 100 | |
| | | | | 9 | 54100 | 54302 | 203 | 203 | 471 | 430 | 100 | 100 | |
| | | | | 10 | 55462 | 55607 | 146 | 146 | 191 | 213 | 100 | 98 | |
| | | | | 11 | 55844 | 55965 | 122 | 122 | 499 | 513 | 100 | 100 | |
| | | | | 12 | 56419 | 56529 | 111 | 111 | 385 | 402 | 100 | 100 | |
| | | | | 13 | 57775 | 57965 | 191 | 191 | 71 | 82 | 98 | 94 | |
| | | | | 14 | 58934 | 59056 | 123 | 123 | 480 | 433 | 100 | 99 | |
| | | | | 15 | 59206 | 59276 | 71 | 71 | 449 | 404 | 100 | 99 | |
| | | | | 16 | 59546 | 59683 | 138 | 138 | 468 | 436 | 100 | 99 | |
| | | | | 17 | 59812 | 59917 | 106 | 106 | 327 | 313 | 100 | 98 | |
| | | | | 18 | 60031 | 62697 | 2667 | 177 | 40 | 41 | 92 | 93 | |
| FGF3 | NG_009016 | NM_005247.2 | CCDS8195.1 | 1 | 5001 | 5711 | 711 | 220 | 1 | 1 | 0 | 0 | exon fail/discard |
| | | | | 2 | 8002 | 8105 | 104 | 104 | 155 | 157 | 100 | 97 | |
| | | | | 3 | 13725 | 14457 | 733 | 396 | 15 | 17 | 68 | 72 | poor quality/discard |
| FGF7 | NG_029159 | NM_002009.3 | CCDS10131.1 | 2 | 5855 | 6406 | 552 | 286 | 457 | 464 | 100 | 100 | |
| | | | | 3 | 64974 | 65077 | 104 | 104 | 1252 | 1117 | 100 | 100 | |
| | | | | 4 | 66133 | 69149 | 3017 | 195 | 210 | 184 | 99 | 99 | |
| FGF10 | NG_011446 | NM_004465.1 | CCDS3950.1 | 1 | 5001 | 5325 | 325 | 325 | 689 | 623 | 100 | 100 | |
| | | | | 2 | 83153 | 83256 | 104 | 104 | 668 | 664 | 100 | 100 | |
| | | | | 3 | 88491 | 88688 | 198 | 198 | 756 | 683 | 100 | 100 | |
| FGF18 | NG_029158 | NM_003862.2 | CCDS4378.1 | 1 | 5001 | 5569 | 569 | 32 | 7 | 7 | 27 | 18 | exon fail/discard |
| | | | | 2 | 5745 | 5781 | 37 | 37 | 66 | 68 | 97 | 94 | |
| | | | | 3 | 21431 | 21611 | 181 | 181 | 538 | 462 | 100 | 98 | |
| | | | | 4 | 34485 | 34591 | 107 | 107 | 1053 | 1036 | 100 | 100 | |
| | | | | 5 | 41877 | 42964 | 1088 | 267 | 185 | 181 | 100 | 97 | |
| FPGS | NG_023245 | NM_004957.5 | CCDS35148.1 | 1 | 4984 | 5188 | 205 | 138 | 2 | 2 | 27 | 26 | exon fail/discard |
| | | | | 2 | 6411 | 6539 | 129 | 129 | 208 | 195 | 100 | 97 | |
| | | | | 3 | 6625 | 6678 | 54 | 54 | 156 | 142 | 100 | 97 | |
| | | | | 4 | 6762 | 6826 | 65 | 65 | 156 | 134 | 100 | 97 | |
| | | | | 5 | 9099 | 9213 | 115 | 115 | 123 | 131 | 100 | 97 | |
| | | | | 6 | 9335 | 9412 | 78 | 78 | 204 | 199 | 100 | 95 | |
| | | | | 7 | 9547 | 9608 | 62 | 62 | 140 | 150 | 100 | 97 | |
| | | | | 8 | 9712 | 9814 | 103 | 103 | 114 | 141 | 100 | 95 | |
| | | | | 9 | 10360 | 10437 | 78 | 78 | 174 | 182 | 100 | 95 | |
| | | | | 10 | 10684 | 10831 | 148 | 148 | 52 | 58 | 97 | 95 | |
| | | | | 11 | 10926 | 11015 | 90 | 90 | 101 | 98 | 99 | 95 | |
| | | | | 12 | 11810 | 11960 | 151 | 151 | 13 | 19 | 76 | 77 | poor quality/discard |
| | | | | 13 | 12157 | 12232 | 76 | 76 | 57 | 69 | 98 | 96 | |
| | | | | 14 | 13071 | 13137 | 67 | 67 | 242 | 251 | 100 | 98 | |
| | | | | 15 | 15321 | 16403 | 1083 | 410 | 120 | 119 | 100 | 95 | |
| GART | NG_029160 | NM_001136005.1 | CCDS13627.1 | 2 | 8570 | 8722 | 153 | 145 | 449 | 498 | 100 | 100 | |
| | | | | 3 | 12573 | 12668 | 96 | 96 | 268 | 269 | 100 | 99 | |
| | | | | 4 | 13140 | 13314 | 175 | 175 | 441 | 440 | 100 | 100 | |
| | | | | 5 | 15437 | 15548 | 112 | 112 | 310 | 335 | 100 | 99 | |
| | | | | 6 | 16336 | 16404 | 69 | 69 | 224 | 221 | 100 | 99 | |
| | | | | 7 | 17009 | 17134 | 126 | 126 | 427 | 457 | 100 | 100 | |
| | | | | 8 | 18956 | 19043 | 88 | 88 | 389 | 406 | 100 | 100 | |
| | | | | 9 | 19289 | 19374 | 86 | 86 | 364 | 366 | 100 | 99 | |
| | | | | 10 | 19558 | 19726 | 169 | 169 | 386 | 412 | 100 | 100 | |
| | | | | 11 | 22892 | 23123 | 232 | 232 | 522 | 540 | 100 | 100 | |
| | | | | 12 | 25610 | 25704 | 95 | 95 | 538 | 580 | 100 | 100 | |
| | | | | 13 | 26877 | 26986 | 110 | 110 | 528 | 590 | 100 | 100 | |
| | | | | 14 | 27330 | 27528 | 199 | 199 | 730 | 787 | 100 | 100 | |
| | | | | 15 | 30284 | 30535 | 252 | 252 | 598 | 597 | 100 | 100 | |
| | | | | 16 | 30751 | 30903 | 153 | 153 | 557 | 571 | 100 | 100 | |
| | | | | 17 | 36434 | 36640 | 207 | 207 | 1124 | 1063 | 100 | 100 | |
| | | | | 18 | 37972 | 38109 | 138 | 138 | 755 | 837 | 100 | 100 | |
| | | | | 19 | 41788 | 41918 | 131 | 131 | 668 | 660 | 100 | 100 | |
| | | | | 20 | 42190 | 42331 | 142 | 142 | 711 | 694 | 100 | 100 | |
| | | | | 21 | 43365 | 43480 | 116 | 116 | 565 | 594 | 100 | 100 | |
| | | | | 22 | 43577 | 43961 | 385 | 192 | 633 | 630 | 100 | 100 | |
| GGH | NG_028126 | NM_003878.2 | CCDS6177.1 | 1 | 5001 | 5392 | 392 | 109 | 12 | 13 | 63 | 62 | poor quality/discard |
| | | | | 2 | 8282 | 8396 | 115 | 115 | 525 | 546 | 100 | 100 | |
| | | | | 3 | 13835 | 13885 | 51 | 51 | 1222 | 1146 | 100 | 100 | |
| | | | | 4 | 16787 | 16871 | 85 | 85 | 1179 | 967 | 100 | 100 | |
| | | | | 5 | 17756 | 17894 | 139 | 139 | 882 | 845 | 100 | 100 | |
| | | | | 6 | 19866 | 19972 | 107 | 107 | 1275 | 1254 | 100 | 100 | |
| | | | | 7 | 20055 | 20145 | 91 | 91 | 852 | 912 | 100 | 100 | |
| | | | | 8 | 26422 | 26559 | 138 | 138 | 1743 | 1652 | 100 | 100 | |
| | | | | 9 | 28599 | 28973 | 375 | 122 | 989 | 948 | 100 | 100 | |
| IRF6 | NG_007081 | NM_006147.3 | CCDS1492.1 | 3 | 9719 | 9895 | 177 | 174 | 341 | 368 | 100 | 99 | |
| | | | | 4 | 14583 | 14787 | 205 | 205 | 701 | 679 | 100 | 100 | |
| | | | | 5 | 15717 | 15845 | 129 | 129 | 1202 | 1262 | 100 | 100 | |
| | | | | 6 | 18708 | 18866 | 159 | 159 | 741 | 732 | 100 | 100 | |
| | | | | 7 | 20248 | 20640 | 393 | 393 | 1020 | 955 | 100 | 100 | |
| | | | | 8 | 21350 | 21468 | 119 | 119 | 1081 | 946 | 100 | 100 | |
| | | | | 9 | 22491 | 25512 | 3022 | 225 | 129 | 134 | 99 | 98 | |
| LRP6 | NG_016168 | NM_002336.2 | CCDS8647.1 | 1 | 5001 | 5197 | 197 | 55 | 44 | 45 | 95 | 94 | |

| | | | | | | | | | | | | | |
|---|---|---|---|---|---|---|---|---|---|---|---|---|---|
| | | | | 2 | 27223 | 27616 | 394 | 394 | 1884 | 1692 | 100 | 100 | |
| | | | | 3 | 68478 | 68675 | 198 | 198 | 631 | 595 | 100 | 100 | |
| | | | | 4 | 84759 | 84955 | 197 | 197 | 1750 | 1588 | 100 | 100 | |
| | | | | 5 | 87767 | 87898 | 132 | 132 | 1307 | 1226 | 100 | 100 | |
| | | | | 6 | 90439 | 90835 | 397 | 397 | 1335 | 1155 | 100 | 100 | |
| | | | | 7 | 91897 | 92068 | 172 | 172 | 1098 | 1091 | 100 | 100 | |
| | | | | 8 | 106583 | 106799 | 217 | 217 | 694 | 729 | 100 | 100 | |
| | | | | 9 | 107316 | 107605 | 290 | 290 | 594 | 671 | 100 | 100 | |
| | | | | 10 | 109459 | 109685 | 227 | 227 | 761 | 767 | 100 | 100 | |
| | | | | 11 | 111914 | 112098 | 185 | 185 | 869 | 925 | 100 | 100 | |
| | | | | 12 | 112723 | 113049 | 327 | 327 | 796 | 816 | 100 | 100 | |
| | | | | 13 | 120840 | 121042 | 203 | 203 | 1694 | 1624 | 100 | 100 | |
| | | | | 14 | 122725 | 122936 | 212 | 212 | 1859 | 1706 | 100 | 100 | |
| | | | | 15 | 124322 | 124512 | 191 | 191 | 1691 | 1686 | 100 | 100 | |
| | | | | 16 | 133344 | 133553 | 210 | 210 | 1135 | 1135 | 100 | 100 | |
| | | | | 17 | 136578 | 136703 | 126 | 126 | 1038 | 1105 | 100 | 100 | |
| | | | | 18 | 139821 | 140057 | 237 | 237 | 887 | 925 | 100 | 100 | |
| | | | | 19 | 140985 | 141095 | 111 | 111 | 727 | 717 | 100 | 100 | |
| | | | | 20 | 144957 | 145187 | 231 | 231 | 1577 | 1485 | 100 | 100 | |
| | | | | 21 | 146446 | 146582 | 137 | 137 | 1538 | 1497 | 100 | 100 | |
| | | | | 22 | 147216 | 147313 | 98 | 98 | 444 | 498 | 100 | 100 | |
| | | | | 23 | 150458 | 155853 | 5396 | 295 | 252 | 196 | 100 | 99 | |
| MAT1A | NG_008083 | NM_000429.2 | CCDS7365.1 | 1 | 5001 | 5346 | 346 | 91 | 286 | 312 | 100 | 100 | |
| | | | | 2 | 9090 | 9167 | 78 | 78 | 564 | 574 | 100 | 100 | |
| | | | | 3 | 10641 | 10763 | 123 | 123 | 543 | 545 | 100 | 100 | |
| | | | | 4 | 13887 | 13999 | 113 | 113 | 1014 | 962 | 100 | 100 | |
| | | | | 5 | 14363 | 14506 | 144 | 144 | 662 | 659 | 100 | 100 | |
| | | | | 6 | 18085 | 18303 | 219 | 219 | 373 | 356 | 100 | 99 | |
| | | | | 7 | 19480 | 19662 | 183 | 183 | 236 | 276 | 100 | 97 | |
| | | | | 8 | 20026 | 20159 | 134 | 134 | 332 | 373 | 100 | 98 | |
| | | | | 9 | 20796 | 22859 | 2064 | 103 | 93 | 94 | 100 | 97 | |
| MAT2A | NG_029183 | NM_005911.5 | CCDS1977.1 | 1 | 5001 | 5401 | 401 | 91 | 71 | 59 | 99 | 95 | |
| | | | | 2 | 7106 | 7183 | 78 | 78 | 357 | 342 | 100 | 100 | |
| | | | | 3 | 7278 | 7400 | 123 | 123 | 407 | 387 | 100 | 100 | |
| | | | | 4 | 7656 | 7768 | 113 | 113 | 409 | 405 | 100 | 100 | |
| | | | | 5 | 7852 | 7995 | 144 | 144 | 509 | 501 | 100 | 100 | |
| | | | | 6 | 8178 | 8396 | 219 | 219 | 516 | 497 | 100 | 100 | |
| | | | | 7 | 8588 | 8770 | 183 | 183 | 381 | 381 | 100 | 100 | |
| | | | | 8 | 8924 | 9057 | 134 | 134 | 590 | 586 | 100 | 100 | |
| | | | | 9 | 9693 | 11303 | 1611 | 103 | 278 | 293 | 100 | 100 | |
| MKX | NG_029181 | NM_173576.2 | CCDS7156.1 | 2 | 7353 | 7622 | 270 | 188 | 62 | 54 | 93 | 95 | |
| | | | | 3 | 9346 | 9505 | 160 | 160 | 215 | 190 | 100 | 97 | |
| | | | | 4 | 15476 | 15629 | 154 | 154 | 822 | 717 | 100 | 100 | |
| | | | | 5 | 16059 | 16394 | 336 | 336 | 652 | 576 | 100 | 100 | |
| | | | | 6 | 75296 | 75329 | 34 | 34 | 612 | 649 | 100 | 100 | |
| | | | | 7 | 75435 | 77976 | 2542 | 187 | 185 | 181 | 100 | 99 | |
| MSX1 | NG_008121 | NM_002448.3 | CCDS3378.2 | 1 | 5001 | 5704 | 704 | 469 | 10 | 14 | 66 | 67 | poor quality/discard |
| | | | | 2 | 8037 | 9272 | 1236 | 443 | 81 | 93 | 99 | 95 | |
| MTFMT | NG_029184 | NM_139242.3 | CCDS45280.1 | 1 | 5001 | 5235 | 235 | 209 | 23 | 25 | 84 | 87 | poor quality/discard |
| | | | | 2 | 7600 | 7809 | 210 | 210 | 689 | 731 | 100 | 100 | |
| | | | | 3 | 10846 | 10968 | 123 | 123 | 441 | 461 | 100 | 100 | |
| | | | | 4 | 13024 | 13126 | 103 | 103 | 520 | 528 | 100 | 100 | |
| | | | | 5 | 14368 | 14443 | 76 | 76 | 366 | 346 | 100 | 99 | |
| | | | | 6 | 18113 | 18204 | 92 | 92 | 410 | 403 | 100 | 100 | |
| | | | | 7 | 28449 | 28527 | 79 | 79 | 924 | 821 | 100 | 100 | |
| | | | | 8 | 29704 | 29786 | 83 | 83 | 756 | 750 | 100 | 100 | |
| | | | | 9 | 31384 | 33128 | 1745 | 195 | 155 | 169 | 100 | 99 | |
| MTHFD1 | NG_012450 | NM_005956.4 | CCDS9763.1 | 1 | 5313 | 5428 | 428 | 41 | 104 | 100 | 99 | 95 | |
| | | | | 2 | 17753 | 17837 | 85 | 85 | 872 | 883 | 100 | 100 | |
| | | | | 3 | 28050 | 28109 | 60 | 60 | 515 | 463 | 100 | 100 | |
| | | | | 4 | 29432 | 29485 | 54 | 54 | 631 | 627 | 100 | 100 | |
| | | | | 5 | 32318 | 32454 | 137 | 137 | 724 | 727 | 100 | 100 | |
| | | | | 6 | 32599 | 32699 | 101 | 101 | 607 | 677 | 100 | 100 | |
| | | | | 7 | 34848 | 34984 | 137 | 137 | 483 | 517 | 100 | 100 | |
| | | | | 8 | 36774 | 36885 | 112 | 112 | 705 | 740 | 100 | 100 | |
| | | | | 9 | 41764 | 41891 | 128 | 128 | 590 | 614 | 100 | 100 | |
| | | | | 10 | 42690 | 42787 | 98 | 98 | 746 | 738 | 100 | 100 | |
| | | | | 11 | 42979 | 43152 | 174 | 174 | 807 | 823 | 100 | 100 | |
| | | | | 12 | 44297 | 44433 | 137 | 137 | 1036 | 1116 | 100 | 100 | |
| | | | | 13 | 47144 | 47190 | 47 | 47 | 806 | 758 | 100 | 100 | |
| | | | | 14 | 48492 | 48599 | 108 | 108 | 956 | 962 | 100 | 100 | |
| | | | | 15 | 48747 | 48821 | 75 | 75 | 798 | 757 | 100 | 100 | |
| | | | | 16 | 52529 | 52631 | 103 | 103 | 987 | 957 | 100 | 100 | |
| | | | | 17 | 56056 | 56132 | 77 | 77 | 656 | 657 | 100 | 100 | |
| | | | | 18 | 57086 | 57226 | 141 | 141 | 406 | 437 | 100 | 100 | |
| | | | | 19 | 58345 | 58413 | 69 | 69 | 574 | 670 | 100 | 100 | |
| | | | | 20 | 59014 | 59125 | 112 | 112 | 713 | 785 | 100 | 100 | |
| | | | | 21 | 59223 | 59362 | 140 | 140 | 603 | 647 | 100 | 100 | |
| | | | | 22 | 61653 | 61694 | 42 | 42 | 555 | 599 | 100 | 100 | |
| | | | | 23 | 65177 | 65277 | 101 | 101 | 345 | 364 | 100 | 100 | |
| | | | | 24 | 66405 | 66582 | 178 | 178 | 411 | 421 | 100 | 100 | |
| | | | | 25 | 70714 | 70821 | 108 | 108 | 448 | 513 | 100 | 100 | |
| | | | | 26 | 71683 | 71835 | 153 | 153 | 394 | 458 | 100 | 100 | |
| | | | | 27 | 75174 | 75267 | 94 | 90 | 340 | 331 | 100 | 99 | |
| MTHFD1L | NG_029185 | NM_001242767.1 | CCDS75535.1 | 1 | 5001 | 5371 | 371 | 227 | 1 | 1 | 0 | 0 | exon fail/discard |
| | | | | 2 | 15412 | 15496 | 85 | 85 | 497 | 523 | 100 | 100 | |
| | | | | 3 | 16956 | 17006 | 51 | 51 | 576 | 629 | 100 | 100 | |
| | | | | 4 | 17095 | 17148 | 54 | 54 | 581 | 613 | 100 | 100 | |
| | | | | 5 | 22084 | 22208 | 125 | 125 | 300 | 343 | 100 | 100 | |
| | | | | 6 | 24956 | 25056 | 101 | 101 | 348 | 371 | 100 | 100 | |
| | | | | 7 | 27164 | 27303 | 140 | 140 | 361 | 364 | 100 | 100 | |
| | | | | 8 | 44972 | 45083 | 112 | 112 | 406 | 444 | 100 | 100 | |
| | | | | 9 | 57899 | 57990 | 92 | 92 | 476 | 493 | 100 | 100 | |
| | | | | 10 | 61527 | 61624 | 98 | 98 | 679 | 667 | 100 | 100 | |
| | | | | 11 | 65444 | 65617 | 174 | 174 | 826 | 810 | 100 | 100 | |
| | | | | 12 | 76126 | 76262 | 137 | 137 | 718 | 727 | 100 | 100 | |
| | | | | 13 | 78021 | 78067 | 47 | 47 | 421 | 393 | 100 | 100 | |

| | | | | | | | | | | | | | |
|---|---|---|---|---|---|---|---|---|---|---|---|---|---|
| | | | | 14 | 83808 | 83915 | 108 | 108 | 1346 | 1403 | 100 | 100 | |
| | | | | 15 | 84789 | 84863 | 75 | 75 | 1403 | 1377 | 100 | 100 | |
| | | | | 16 | 88353 | 88455 | 103 | 103 | 1692 | 1569 | 100 | 100 | |
| | | | | 17 | 95317 | 95393 | 77 | 77 | 329 | 317 | 100 | 99 | |
| | | | | 18 | 99597 | 99737 | 141 | 141 | 308 | 268 | 100 | 98 | |
| | | | | 19 | 104291 | 104359 | 69 | 69 | 698 | 783 | 100 | 100 | |
| | | | | 20 | 111269 | 111380 | 112 | 112 | 681 | 725 | 100 | 100 | |
| | | | | 21 | 149141 | 149280 | 140 | 140 | 824 | 719 | 100 | 100 | |
| | | | | 22 | 153101 | 153142 | 42 | 42 | 520 | 497 | 100 | 100 | |
| | | | | 23 | 154202 | 154302 | 101 | 101 | 715 | 695 | 100 | 100 | |
| | | | | 24 | 154838 | 155015 | 178 | 178 | 646 | 611 | 100 | 100 | |
| | | | | 25 | 173815 | 173922 | 108 | 108 | 577 | 571 | 100 | 100 | |
| | | | | 26 | 176287 | 176439 | 153 | 153 | 586 | 577 | 100 | 100 | |
| | | | | 27 | 231789 | 231909 | 121 | 90 | 1004 | 976 | 100 | 100 | |
| MTHFD2 | NG_029162 | NM_006636.3 | CCDS1935.2 | 1 | 5001 | 5180 | 180 | 101 | 103 | 112 | 97 | 95 | |
| | | | | 2 | 12143 | 12327 | 185 | 185 | 732 | 728 | 100 | 100 | |
| | | | | 3 | 14142 | 14264 | 123 | 123 | 813 | 802 | 100 | 100 | |
| | | | | 4 | 15007 | 15159 | 153 | 153 | 698 | 751 | 100 | 100 | |
| | | | | 5 | 16380 | 16487 | 108 | 108 | 592 | 574 | 100 | 100 | |
| | | | | 6 | 17637 | 17729 | 93 | 93 | 824 | 926 | 100 | 100 | |
| | | | | 7 | 18179 | 18304 | 126 | 126 | 505 | 548 | 100 | 100 | |
| | | | | 8 | 20517 | 21736 | 1220 | 164 | 261 | 268 | 100 | 99 | |
| MTHFR | NG_013351 | NM_005957.4 | CCDS137.1 | 2 | 7975 | 8223 | 249 | 236 | 293 | 264 | 100 | 99 | |
| | | | | 3 | 9705 | 9943 | 239 | 239 | 198 | 215 | 100 | 97 | |
| | | | | 4 | 10782 | 10892 | 111 | 111 | 264 | 270 | 100 | 99 | |
| | | | | 5 | 14705 | 14898 | 194 | 194 | 137 | 145 | 100 | 95 | |
| | | | | 6 | 15756 | 16006 | 251 | 251 | 136 | 131 | 100 | 95 | |
| | | | | 7 | 16241 | 16375 | 135 | 135 | 136 | 152 | 100 | 95 | |
| | | | | 8 | 16566 | 16746 | 181 | 181 | 196 | 221 | 100 | 99 | |
| | | | | 9 | 17015 | 17197 | 183 | 183 | 120 | 126 | 99 | 94 | |
| | | | | 10 | 18725 | 18826 | 102 | 102 | 171 | 168 | 100 | 98 | |
| | | | | 11 | 19778 | 19897 | 120 | 120 | 271 | 287 | 100 | 98 | |
| | | | | 12 | 20206 | 25374 | 5169 | 219 | 98 | 106 | 100 | 95 | |
| MTHFS | NG_029243 | NM_001199758.1 | NP_001186687 | 2 | 12932 | 13193 | 262 | 208 | 1542 | 1408 | 100 | 100 | |
| | | | | 3 | 56844 | 58739 | 1896 | 233 | 448 | 423 | 100 | 100 | |
| MTR | NG_008959 | NM_000254.2 | CCDS1614.1 | 1 | 5001 | 5457 | 457 | 34 | 48 | 51 | 97 | 94 | |
| | | | | 2 | 13148 | 13362 | 215 | 215 | 995 | 1015 | 100 | 100 | |
| | | | | 3 | 15864 | 15953 | 90 | 90 | 1210 | 1132 | 100 | 100 | |
| | | | | 4 | 18424 | 18493 | 70 | 70 | 665 | 606 | 100 | 100 | |
| | | | | 5 | 20223 | 20315 | 93 | 93 | 767 | 829 | 100 | 100 | |
| | | | | 6 | 22458 | 22564 | 107 | 107 | 908 | 919 | 100 | 100 | |
| | | | | 7 | 25324 | 25383 | 60 | 60 | 687 | 610 | 100 | 100 | |
| | | | | 8 | 26169 | 26263 | 95 | 95 | 880 | 934 | 100 | 100 | |
| | | | | 9 | 33839 | 33939 | 101 | 101 | 1044 | 1006 | 100 | 100 | |
| | | | | 10 | 35058 | 35119 | 62 | 62 | 930 | 949 | 100 | 100 | |
| | | | | 11 | 36549 | 36616 | 68 | 68 | 854 | 829 | 100 | 100 | |
| | | | | 12 | 38909 | 38988 | 80 | 80 | 669 | 670 | 100 | 100 | |
| | | | | 13 | 41686 | 41798 | 113 | 113 | 945 | 862 | 100 | 100 | |
| | | | | 14 | 45267 | 45407 | 141 | 141 | 976 | 934 | 100 | 100 | |
| | | | | 15 | 48134 | 48319 | 186 | 186 | 759 | 766 | 100 | 100 | |
| | | | | 16 | 60064 | 60243 | 180 | 180 | 1387 | 1391 | 100 | 100 | |
| | | | | 17 | 62241 | 62357 | 117 | 117 | 1144 | 1217 | 100 | 100 | |
| | | | | 18 | 62668 | 62808 | 141 | 141 | 1490 | 1422 | 100 | 100 | |
| | | | | 19 | 69553 | 69642 | 90 | 90 | 619 | 582 | 100 | 100 | |
| | | | | 20 | 70845 | 70997 | 153 | 153 | 677 | 651 | 100 | 100 | |
| | | | | 21 | 71956 | 72063 | 108 | 108 | 979 | 994 | 100 | 100 | |
| | | | | 22 | 73174 | 73274 | 101 | 101 | 1050 | 1029 | 100 | 100 | |
| | | | | 23 | 83493 | 83560 | 68 | 68 | 629 | 650 | 100 | 100 | |
| | | | | 24 | 84446 | 84566 | 121 | 121 | 430 | 460 | 100 | 100 | |
| | | | | 25 | 90475 | 90556 | 82 | 82 | 508 | 523 | 100 | 100 | |
| | | | | 26 | 94841 | 94939 | 99 | 99 | 521 | 554 | 100 | 100 | |
| | | | | 27 | 96012 | 96087 | 76 | 76 | 647 | 712 | 100 | 100 | |
| | | | | 28 | 98901 | 99056 | 156 | 156 | 665 | 634 | 100 | 100 | |
| | | | | 29 | 100853 | 101049 | 197 | 197 | 1116 | 1018 | 100 | 100 | |
| | | | | 30 | 104077 | 104277 | 201 | 201 | 1114 | 1175 | 100 | 100 | |
| | | | | 31 | 105078 | 105270 | 193 | 193 | 806 | 874 | 100 | 100 | |
| | | | | 32 | 106726 | 106838 | 113 | 113 | 914 | 958 | 100 | 100 | |
| | | | | 33 | 107278 | 113701 | 6424 | 87 | 89 | 66 | 99 | 95 | |
| MTRR | NG_008856 | NM_002454.2 | CCDS47190.1 | 2 | 6667 | 6820 | 154 | 129 | 1288 | 1253 | 100 | 100 | |
| | | | | 3 | 9270 | 9423 | 154 | 154 | 1187 | 1178 | 100 | 100 | |
| | | | | 4 | 11155 | 11272 | 118 | 118 | 875 | 909 | 100 | 100 | |
| | | | | 5 | 13841 | 14219 | 379 | 379 | 695 | 738 | 100 | 100 | |
| | | | | 6 | 19052 | 19174 | 123 | 123 | 1075 | 1018 | 100 | 100 | |
| | | | | 7 | 21598 | 21751 | 154 | 154 | 929 | 1026 | 100 | 100 | |
| | | | | 8 | 22512 | 22600 | 89 | 89 | 1198 | 1282 | 100 | 100 | |
| | | | | 9 | 24992 | 25172 | 181 | 181 | 1491 | 1217 | 100 | 100 | |
| | | | | 10 | 27269 | 27311 | 43 | 43 | 615 | 613 | 100 | 100 | |
| | | | | 11 | 28624 | 28810 | 187 | 187 | 1169 | 1121 | 100 | 100 | |
| | | | | 12 | 31631 | 31749 | 119 | 119 | 955 | 1024 | 100 | 100 | |
| | | | | 13 | 32761 | 32853 | 93 | 93 | 846 | 920 | 100 | 100 | |
| | | | | 14 | 32962 | 33144 | 183 | 183 | 846 | 844 | 100 | 100 | |
| | | | | 15 | 35811 | 37021 | 1211 | 145 | 371 | 332 | 100 | 100 | |
| NECTIN1 | NG_013083 | NM_002855.4 | CCDS8426.1 | 1 | 1 | 251 | 251 | 79 | 5 | 3 | 38 | 40 | poor quality/discard |
| | | | | 2 | 49961 | 50311 | 351 | 351 | 197 | 201 | 100 | 97 | |
| | | | | 3 | 50869 | 51171 | 303 | 303 | 323 | 361 | 100 | 98 | |
| | | | | 4 | 51507 | 51624 | 118 | 118 | 237 | 239 | 100 | 98 | |
| | | | | 5 | 53416 | 53567 | 152 | 152 | 160 | 176 | 100 | 97 | |
| | | | | 6 | 63429 | 67734 | 4306 | 551 | 115 | 116 | 99 | 97 | |
| PTCH1 | NG_007664 | NM_000264.4 | CCDS6714.1 | 1 | 13417 | 13805 | 389 | 201 | 13 | 12 | 66 | 62 | poor quality/discard |
| | | | | 2 | 15367 | 15559 | 193 | 193 | 121 | 85 | 99 | 95 | |
| | | | | 3 | 36092 | 36281 | 190 | 190 | 590 | 614 | 100 | 100 | |
| | | | | 4 | 39763 | 39832 | 70 | 70 | 1094 | 1145 | 100 | 100 | |
| | | | | 5 | 39926 | 40017 | 92 | 92 | 1184 | 1197 | 100 | 100 | |
| | | | | 6 | 41378 | 41576 | 199 | 199 | 735 | 802 | 100 | 100 | |
| | | | | 7 | 41876 | 41997 | 122 | 122 | 1178 | 1172 | 100 | 100 | |
| | | | | 8 | 42819 | 42966 | 148 | 148 | 1596 | 1707 | 100 | 100 | |
| | | | | 9 | 43780 | 43911 | 132 | 132 | 1064 | 1086 | 100 | 100 | |

| | | | | | | | | | | | | | |
|---|---|---|---|---|---|---|---|---|---|---|---|---|---|
| | | | | 10 | 44264 | 44419 | 156 | 156 | 1111 | 1169 | 100 | 100 | |
| | | | | 11 | 45109 | 45207 | 99 | 99 | 2136 | 2235 | 100 | 100 | |
| | | | | 12 | 45807 | 45932 | 126 | 126 | 781 | 833 | 100 | 100 | |
| | | | | 13 | 52035 | 52153 | 119 | 119 | 420 | 398 | 100 | 100 | |
| | | | | 14 | 52813 | 53215 | 403 | 403 | 250 | 255 | 100 | 97 | |
| | | | | 15 | 54541 | 54850 | 310 | 310 | 386 | 379 | 100 | 100 | |
| | | | | 16 | 59968 | 60110 | 143 | 143 | 288 | 317 | 100 | 99 | |
| | | | | 17 | 62183 | 62366 | 184 | 184 | 210 | 214 | 100 | 98 | |
| | | | | 18 | 63673 | 63953 | 281 | 281 | 203 | 189 | 100 | 98 | |
| | | | | 19 | 65553 | 65690 | 138 | 138 | 192 | 195 | 100 | 98 | |
| | | | | 20 | 68346 | 68488 | 143 | 143 | 301 | 316 | 100 | 98 | |
| | | | | 21 | 72026 | 72125 | 100 | 100 | 348 | 350 | 100 | 100 | |
| | | | | 22 | 72643 | 72897 | 255 | 255 | 188 | 209 | 99 | 97 | |
| | | | | 23 | 74515 | 75055 | 541 | 540 | 193 | 213 | 100 | 97 | |
| SARDH | NG_008987 | NM_007101.3 | CCDS6978.1 | 2 | 10753 | 11113 | 361 | 331 | 247 | 293 | 99 | 98 | |
| | | | | 3 | 12355 | 12533 | 179 | 179 | 225 | 282 | 100 | 98 | |
| | | | | 4 | 13472 | 13651 | 180 | 180 | 621 | 593 | 100 | 99 | |
| | | | | 5 | 14769 | 14892 | 124 | 124 | 562 | 556 | 100 | 100 | |
| | | | | 6 | 15091 | 15191 | 101 | 101 | 523 | 509 | 100 | 99 | |
| | | | | 7 | 25914 | 26018 | 105 | 105 | 215 | 263 | 100 | 98 | |
| | | | | 8 | 27501 | 27630 | 130 | 130 | 269 | 283 | 100 | 97 | |
| | | | | 9 | 31832 | 31918 | 87 | 87 | 95 | 110 | 99 | 94 | |
| | | | | 10 | 32247 | 32337 | 91 | 91 | 137 | 139 | 100 | 97 | |
| | | | | 11 | 36528 | 36669 | 142 | 142 | 85 | 95 | 99 | 95 | |
| | | | | 12 | 39925 | 40008 | 84 | 84 | 149 | 150 | 100 | 97 | |
| | | | | 13 | 41927 | 42040 | 114 | 114 | 143 | 157 | 99 | 97 | |
| | | | | 14 | 48595 | 48733 | 139 | 139 | 313 | 290 | 100 | 98 | |
| | | | | 15 | 50585 | 50698 | 114 | 114 | 46 | 49 | 94 | 94 | |
| | | | | 16 | 54429 | 54576 | 148 | 148 | 109 | 109 | 100 | 95 | |
| | | | | 17 | 59670 | 59763 | 94 | 94 | 202 | 200 | 100 | 97 | |
| | | | | 18 | 73259 | 73421 | 163 | 163 | 20 | 22 | 83 | 84 | poor quality/discard |
| | | | | 19 | 74204 | 74372 | 169 | 169 | 8 | 12 | 69 | 70 | poor quality/discard |
| | | | | 20 | 78086 | 78221 | 136 | 136 | 35 | 34 | 92 | 91 | |
| | | | | 21 | 80942 | 81396 | 455 | 126 | 77 | 85 | 98 | 95 | |
| SHMT1 | NG_017111 | NM_004169.4 | CCDS11196.1 | 2 | 12543 | 12657 | 115 | 96 | 212 | 214 | 100 | 99 | |
| | | | | 3 | 14726 | 14871 | 146 | 146 | 392 | 404 | 100 | 99 | |
| | | | | 4 | 20103 | 20218 | 116 | 116 | 262 | 289 | 100 | 98 | |
| | | | | 5 | 20887 | 21047 | 161 | 161 | 258 | 308 | 100 | 99 | |
| | | | | 6 | 27730 | 27811 | 82 | 82 | 207 | 194 | 100 | 98 | |
| | | | | 7 | 28288 | 28500 | 213 | 213 | 164 | 165 | 100 | 97 | |
| | | | | 8 | 32868 | 32984 | 117 | 117 | 278 | 309 | 100 | 99 | |
| | | | | 9 | 35255 | 35377 | 123 | 123 | 291 | 309 | 100 | 99 | |
| | | | | 10 | 37872 | 37988 | 117 | 117 | 258 | 263 | 100 | 99 | |
| | | | | 11 | 39155 | 39265 | 111 | 111 | 223 | 250 | 100 | 99 | |
| | | | | 12 | 39624 | 40683 | 1047 | 170 | 114 | 115 | 100 | 97 | |
| SHMT2 | NG_029163 | NM_005412.5 | CCDS8934.1 | 1 | 5001 | 5239 | 239 | 33 | 68 | 76 | 100 | 94 | |
| | | | | 2 | 6231 | 6428 | 198 | 198 | 157 | 187 | 100 | 95 | |
| | | | | 3 | 6909 | 6988 | 80 | 80 | 134 | 150 | 100 | 95 | |
| | | | | 4 | 7141 | 7341 | 201 | 201 | 219 | 265 | 100 | 97 | |
| | | | | 5 | 7639 | 7720 | 82 | 82 | 266 | 281 | 100 | 98 | |
| | | | | 6 | 7881 | 8003 | 123 | 123 | 223 | 237 | 100 | 98 | |
| | | | | 7 | 8132 | 8271 | 140 | 140 | 257 | 268 | 100 | 98 | |
| | | | | 8 | 8608 | 8773 | 166 | 166 | 296 | 300 | 100 | 99 | |
| | | | | 9 | 8991 | 9090 | 100 | 100 | 279 | 314 | 100 | 98 | |
| | | | | 10 | 9177 | 9332 | 156 | 156 | 262 | 278 | 100 | 98 | |
| | | | | 11 | 9431 | 9538 | 108 | 108 | 314 | 313 | 100 | 99 | |
| | | | | 12 | 9662 | 10363 | 702 | 128 | 150 | 173 | 100 | 98 | |
| SOX4 | NG_029166 | NM_003107.2 | CCDS4547.1 | 1 | 5001 | 9879 | 4879 | 1425 | 46 | 43 | 95 | 94 | |
| SP8 | NG_029244 | NM_198956.3 | CCDS5372.1 | 3 | 6095 | 9615 | 3521 | 1473 | 45 | 42 | 92 | 90 | |
| TFAP2A | NG_016151 | NM_001042425 | CCDS43422.1 | 1 | 5001 | 5147 | 147 | 33 | 0 | 0 | 0 | 0 | exon fail/discard |
| | | | | 2 | 14230 | 14664 | 435 | 435 | 74 | 100 | 99 | 95 | |
| | | | | 3 | 17721 | 17772 | 52 | 52 | 507 | 507 | 100 | 100 | |
| | | | | 4 | 19826 | 20057 | 232 | 232 | 153 | 147 | 100 | 97 | |
| | | | | 5 | 21955 | 22073 | 119 | 119 | 576 | 678 | 100 | 100 | |
| | | | | 6 | 23976 | 24117 | 142 | 142 | 612 | 645 | 100 | 100 | |
| | | | | 7 | 25860 | 27882 | 2023 | 289 | 141 | 147 | 99 | 97 | |
| TYMS | NG_028255 | NM_001071.2 | CCDS11821.1 | 1 | 5001 | 5344 | 344 | 205 | 12 | 17 | 65 | 66 | poor quality/discard |
| | | | | 2 | 7038 | 7111 | 74 | 74 | 306 | 325 | 100 | 100 | |
| | | | | 3 | 9543 | 9717 | 175 | 175 | 490 | 583 | 100 | 100 | |
| | | | | 4 | 16469 | 16570 | 102 | 102 | 1215 | 1202 | 100 | 100 | |
| | | | | 5 | 18089 | 18264 | 176 | 176 | 1343 | 1310 | 100 | 100 | |
| | | | | 6 | 18777 | 18848 | 72 | 72 | 1412 | 1487 | 100 | 100 | |
| | | | | 7 | 20257 | 20896 | 640 | 138 | 458 | 479 | 100 | 100 | |
| VANGL2 | NG_023420 | NM_020335.2 | CCDS30915.1 | 2 | 20075 | 20335 | 261 | 71 | 190 | 169 | 99 | 98 | |
| | | | | 3 | 20489 | 20609 | 121 | 121 | 403 | 377 | 100 | 98 | |
| | | | | 4 | 23429 | 24036 | 608 | 608 | 817 | 874 | 100 | 100 | |
| | | | | 5 | 24838 | 24974 | 137 | 137 | 2077 | 1971 | 100 | 100 | |
| | | | | 6 | 25479 | 25614 | 136 | 136 | 813 | 777 | 100 | 100 | |
| | | | | 7 | 28479 | 28710 | 232 | 232 | 1317 | 1275 | 100 | 100 | |
| | | | | 8 | 29545 | 33105 | 3561 | 261 | 227 | 190 | 100 | 99 | |
| WNT9B | NG_029164 | NM_003396.2 | CCDS11506.1 | 1 | 4985 | 5114 | 114 | 77 | 1 | 1 | 0 | 0 | exon fail/discard |
| | | | | 2 | 25916 | 26172 | 257 | 257 | 266 | 293 | 99 | 98 | |
| | | | | 3 | 28500 | 28765 | 266 | 266 | 310 | 292 | 100 | 98 | |
| | | | | 4 | 29644 | 33959 | 4316 | 474 | 173 | 177 | 99 | 97 | |
