## Supplementary material for "Accumulation of Rare Coding Variants in Genes Implicated in Risk of Human Cleft Lip with or without Cleft Palate": Table S2

**Table S2. Concordance in genotype calling by TaqMan allelic discrimination VS Illumina sequencing**

1) 476 samples were TaqMan genotyped for 91 variants in 24 of the canddiate genes.
2) Concordance of variant calls is shown at all positions where a variant was called by either method. Coordinate is from the NG RefSeq (Table S1)
3) Genotypes are only shown for discordancies; gray-shading are positions with multiple discordant events
4) For 8,537 variant calls, we observed 99.7% concordance between the two methods.

| Gene | Coordinate (based on NG refseq) | Concordant Genotypes | Discordant Genotypes | Illumina Genotype | TaqMan Gentoype | Class | Variant ID | Amino Acid Affected |
|---|---|---|---|---|---|---|---|---|
| AHCY | 21301 | 16 | 0 | | | Non-synonymous | rs13043752 | R10W |
| AHCY | 24367 | 3 | 0 | | | Non-synonymous | rs41301825 | G95R |
| ALDH1L1 | 32102 | 82 | 0 | | | Non-synonymous | rs3796191 | L254P |
| ALDH1L1 | 32193 | 48 | 0 | | | Synonymous | rs16837178 | D284D |
| ALDH1L1 | 48760 | 2 | 0 | | | Non-synonymous | rs116683776 | V409M |
| ALDH1L1 | 48821 | 14 | 0 | | | Non-synonymous | rs9282691 | E429A |
| ALDH1L1 | 50077 | 116 | 0 | | | Non-synonymous | rs2276724 | S481G |
| ALDH1L1 | 78427 | 138 | 1 | A/G | A/A | Non-synonymous | rs1127717 | D793G |
| ALDH1L1 | 78483 | 43 | 0 | | | Non-synonymous | rs4646750 | I812V |
| ALDH1L2 | 49819 | 266 | 1 | A/T | T/T | Synonymous | rs4964317 | G671G |
| AMT | 8354 | 9 | 0 | | | Non-synonymous | rs116192290 | E211K |
| AMT | 9782 | 204 | 3 | G/A | A/A | Synonymous | rs11715915 | R318R |
| ATIC | 18342 | 278 | 1 | C/C | C/G | Non-synonymous | rs2372536 | T116S |
| ATIC | 39894 | 5 | 0 | | | Non-synonymous | rs56117859 | P471S |
| ATIC | 39977 | 4 | 0 | | | Synonymous | rs116255553 | T498T |
| BHMT | 14555 | 2 | 0 | | | Non-synonymous | rs59866108 | G199S |
| BHMT | 19356 | 275 | 0 | | | Non-synonymous | rs3733890 | R239Q |
| BHMT | 21018 | 49 | 0 | | | Synonymous | rs60340837 | Y284Y |
| BHMT2 | 12885 | 393 | 0 | | | Synonymous | rs682985 | D54D |
| BHMT2 | 14676 | 8 | 0 | | | Non-synonymous | rs60158007 | A66V |
| CBS | 12321 | 2 | 0 | | | Non-synonymous | rs192232907 | K72I |
| CBS | 12410 | 5 | 0 | | | Non-synonymous | rs34040148 | K102Q |
| CBS | 20404 | 3 | 0 | | | Synonymous | rs61735859 | T353T |
| CBS | 20425 | 188 | 5 | C/T | T/T | Synonymous | rs1801181 | A360A |
| CTH | 18053 | 2 | 0 | | | Non-synonymous | rs149505686 | V166M |
| CTH | 32846 | 275 | 0 | | | Non-synonymous | rs1021737 | S403I |
| CTH | 9716 | 5 | 0 | | | Non-synonymous | rs28941785 | T67I |
| DMGDH | 30164 | 297 | 0 | | | Non-synonymous | rs532964 | S279P |
| DMGDH | 30227 | 2 | 1 | C/T | T/T | Non-synonymous | s145258663 | L300F |
| DMGDH | 32248 | 8 | 0 | | | Non-synonymous | rs77116243 | N366S |
| DMGDH | 46098 | 185 | 0 | | | Non-synonymous | rs1805074 | S646P |
| DMGDH | 69230 | 2 | 0 | | | Non-synonymous | rs75051122 | F754S |
| DMGDH | 69278 | 2 | 0 | | | Non-synonymous | rs41272262 | R770Q |
| DMGDH | 43700 | 185 | 1 | G/G | G/C | Non-synonymous | rs1805073 | A530G |
| FPGS | 15362 | 3 | 0 | | | Non-synonymous | rs35789560 | R466C |
| GART | 23086 | 109 | 1 | G/A | A/A | Non-synonymous | rs60421747 | V421I |
| GART | 27355 | 2 | 0 | | | Non-synonymous | rs35927582 | D510G |
| GART | 30476 | 6 | 0 | | | Non-synonymous | rs59920090 | A632V |
| GART | 36581 | 176 | 1 | A/G | A/A | Non-synonymous | rs8971 | D752G |
| GART | 43661 | 3 | 1 | G/G | G/A | Non-synonymous | rs9636610 | V976I |
| GGH | 17756 | 2 | 0 | | | Non-synonymous | rs746403595 | S121G |
| GGH | 17847 | 56 | 0 | | | Non-synonymous | rs11545078 | T151I |
| MAT1A | 14383 | 163 | 0 | | | Synonymous | rs1143694 | A142A |
| MAT1A | 19581 | 176 | 0 | | | Synonymous | rs60582388 | V290V |
| MAT1A | 20841 | 365 | 0 | | | Synonymous | rs57257983 | Y377Y |
| MAT1A | 19593 | 177 | 0 | | | Synonymous | rs59923268 | A294A |
| MAT2A | 8611 | 310 | 0 | | | Synonymous | rs1078004 | R264R |
| MTFMT | 13082 | 4 | 0 | | | Non-synonymous | rs111388106 | A201T |
| MTFMT | 29717 | 9 | 0 | | | Synonymous | rs34636936 | T302T |
| MTHFD1 | 42712 | 2 | 0 | | | Non-synonymous | rs34181110 | R293H |
| MTHFD1 | 17764 | 3 | 0 | | | Non-synonymous | rs151019303 | A18V |
| MTHFD1 | 32622 | 120 | 1 | A/G | G/G | Non-synonymous | rs1950902 | K134R |
| MTHFD1 | 59087 | 355 | 1 | G/A | A/A | Non-synonymous | rs2236225 | R653Q |
| MTHFD1 | 66430 | 9 | 0 | | | Non-synonymous | rs17857382 | L769F |
| MTHFR | 14783 | 298 | 0 | | | Non-synonymous | rs59514310 | A222V |
| MTHFR | 16265 | 89 | 0 | | | Synonymous | rs2066462 | S352S |
| MTHFR | 16685 | 184 | 1 | A/C | C/C | Non-synonymous | rs1801131 | E429A |
| MTHFR | 16704 | 25 | 0 | | | Synonymous | rs57431061 | F435F |
| MTHFR | 20411 | 11 | 0 | | | Non-synonymous | rs35737219 | T653M |
| MTHFR | 20234 | 62 | 0 | | | Non-synonymous | rs58316272 | R594Q |
| MTHFR | 8104 | 84 | 1 | C/T | T/T | Synonymous | rs2066470 | P39P |
| MTHFS | 57068 | 38 | 0 | | | Non-synonymous | rs8923 | T145A |
| MTR | 13268 | 5 | 0 | | | Non-synonymous | rs12749581 | R52Q |
| MTR | 36561 | 22 | 0 | | | Non-synonymous | rs2229274 | D314N |
| MTR | 94867 | 3 | 0 | | | Synonymous | rs780615625 | L901L |
| MTR | 94920 | 166 | 0 | | | Non-synonymous | rs1805087 | D919G |

| | | | | | | | | |
|---|---|---|---|---|---|---|---|---|
| MTR | 96051 | 2 | 0 | | | Non-synonymous | rs113042166 | G939R |
| MTR | 105248 | 300 | 1 | C/T | T/T | Synonymous | rs1131449 | L1192L |
| MTRR | 6757 | 250 | 0 | | | Non-synonymous | rs1801394 | I22M |
| MTRR | 13902 | 4 | 0 | | | Synonymous | rs556611332 | R155R |
| MTRR | 13963 | 200 | 0 | | | Non-synonymous | rs1532268 | S175L |
| MTRR | 14208 | 60 | 0 | | | Non-synonymous | rs2303080 | S257T |
| MTRR | 21691 | 12 | 0 | | | Non-synonymous | rs10064631 | L333V |
| MTRR | 21743 | 233 | 1 | A/G | G/G | Non-synonymous | rs162036 | K350R |
| MTRR | 25088 | 98 | 0 | | | Non-synonymous | rs2287780 | R415C |
| MTRR | 27290 | 92 | 0 | | | Non-synonymous | rs16879334 | P450R |
| MTRR | 28721 | 2 | 0 | | | Non-synonymous | rs41283145 | T490A |
| MTRR | 32975 | 224 | 1 | C/T | T/T | Non-synonymous | rs10380 | H595Y |
| MTRR | 33103 | 200 | 0 | | | Synonymous | rs1802059 | A637A |
| SARDH | 10846 | 3 | 0 | | | Non-synonymous | rs35559818 | G22C |
| SARDH | 10932 | 214 | 1 | G/A | G/G | Synonymous | rs573904 | Q50Q |
| SARDH | 13527 | 2 | 0 | | | Non-synonymous | rs149810392 | T189I |
| SARDH | 27596 | 10 | 0 | | | Non-synonymous | rs35218200 | E372D |
| SARDH | 54449 | 329 | 1 | A/G | A/A | Non-synonymous | rs886016 | M648V |
| SHMT1 | 28333 | 7 | 0 | | | Non-synonymous | rs78909145 | K216R |
| SHMT1 | 28439 | 3 | 1 | G/A | G/G | Synonymous | rs141575508 | V251V |
| SHMT2 | 6346 | 13 | 0 | | | Non-synonymous | rs73338162 | S50L |
| SHMT2 | 7663 | 15 | 0 | | | Synonymous | rs11557166 | D179D |
| SHMT2 | 8227 | 25 | 1 | G/A | G/G | Synonymous | rs2229716 | A271A |
| SHMT2 | 8719 | 57 | 0 | | | Synonymous | rs2229717 | L323L |
| SHMT2 | 9716 | 2 | 0 | | | Non-synonymous | rs536394351 | R481H |
| **TOTAL** | **TOTAL** | **8510** | **27** | | | | | |
