## Supplementary material for "Accumulation of Rare Coding Variants in Genes Implicated in Risk of Human Cleft Lip with or without Cleft Palate": Table S3

**Verification of rare variants by resequencing.**

1) All putative loss-of-function (LOF) alleles plus all rare protein-altering variants from genes with strong CLP case representation were verified by Sanger sequencing from the original bloodspot gDNA preparation (prior to Whole Genome Amplification) using exon-specific primers as described in Methods.

2) Outside of one frameshift in a low-complexity region (see below), all occurrences of all such variants were verified.

[1] according to NG refseq (Table S1)

[2] # called in the final QC'ed dataset, prior to verification resequencing

[3] # confirmed by Sanger sequencing using exon-specific primers as described in Methods

| | Gene | Coordinate[1] | Type | Variation | Amino acid change | # Occurrences called[2] | # Confirmed[3] | Comment |
|---|---|---|---|---|---|---|---|---|
| Nonsynonymous Variants | BHMT | 13597 | SNV | G/A | D105N | 1 | 1 | |
| | BHMT | 13648 | SNV | G/A | V122M | 1 | 1 | |
| | BHMT | 13690 | SNV | G/A | E136K | 1 | 1 | |
| | BHMT | 14549 | SNV | C/T | P197S | 3 | 3 | |
| | BHMT | 14555 | SNV | G/A | G199S | 2 | 2 | |
| | BHMT2 | 14676 | SNV | C/T | A66V | 12 | 12 | |
| | BHMT2 | 18600 | SNV | G/A | A244T | 1 | 1 | |
| | BHMT2 | 18908 | SNV | T/C | L262P | 1 | 1 | |
| | BHMT2 | 23804 | SNV | G/A | A349T | 1 | 1 | |
| | BHMT2 | 23814 | SNV | G/A | R352K | 1 | 1 | |
| | BHMT2 | 23817 | SNV | C/A | P353H | 1 | 1 | |
| | BMP4 | 9782 | SNV | C/A | D56E | 1 | 1 | |
| | BMP4 | 10994 | SNV | G/A | R139H | 1 | 1 | |
| | BMP4 | 11063 | SNV | G/A | R162Q | 1 | 1 | |
| | BMP4 | 11246 | SNV | G/A | R223H | 1 | 1 | |
| | BMP4 | 11255 | SNV | G/A | R226Q | 1 | 1 | |
| | BMP4 | 11384 | SNV | G/A | R269Q | 1 | 1 | |
| | BMPR1B | 351515 | SNV | C/T | P23S | 1 | 1 | |
| | BMPR1B | 361800 | SNV | T/A | V67D | 1 | 1 | |
| | BMPR1B | 370902 | SNV | G/A | V140I | 2 | 2 | |
| | BMPR1B | 370929 | SNV | C/T | R149W | 1 | 1 | |
| | BMPR1B | 372018 | SNV | A/G | Q153R | 1 | 1 | |
| | BMPR1B | 378361 | SNV | A/G | M301V | 1 | 1 | |
| | BMPR1B | 395807 | SNV | G/A | R371Q | 1 | 1 | |
| | BMPR1B | 395934 | SNV | A/T | R413S | 1 | 1 | |
| | DMGDH | 20322 | SNV | C/G | T140S | 1 | 1 | |
| | DMGDH | 20332 | SNV | G/C | R143S | 1 | 1 | |
| | DMGDH | 30203 | SNV | C/T | R292C | 1 | 1 | |
| | DMGDH | 30227 | SNV | C/T | L300F | 3 | 3 | |
| | DMGDH | 32248 | SNV | A/G | N366S | 11 | 11 | |
| | DMGDH | 32257 | SNV | A/G | N369S | 1 | 1 | |
| | DMGDH | 45996 | SNV | G/A | V612I | 1 | 1 | |
| | DMGDH | 46051 | SNV | T/C | L630P | 1 | 1 | |
| | DMGDH | 46089 | SNV | A/G | K643E | 1 | 1 | |
| | DMGDH | 69230 | SNV | T/C | F754S | 2 | 2 | |
| | DMGDH | 69278 | SNV | G/A | R770Q | 3 | 3 | |
| | WNT9B | 25963 | SNV | C/T | A42V | 1 | 1 | |
| | WNT9B | 25978 | SNV | A/G | Q47R | 2 | 2 | |
| | WNT9B | 26119 | SNV | G/A | R94Q | 3 | 3 | |
| | WNT9B | 26134 | SNV | A/G | N99S | 1 | 1 | |
| | WNT9B | 28541 | SNV | G/A | A126T | 3 | 3 | |
| | WNT9B | 28717 | SNV | G/C | K184N | 1 | 1 | |
| | WNT9B | 29708 | SNV | G/A | R222H | 1 | 1 | |
| | WNT9B | 29774 | SNV | C/T | S244L | 2 | 2 | |
| Putative LOF Variants | GGH | 19935 | Insertion/Frameshift | -/T | L190fs | 1 | 1 | |
| | IRF6 | 14742 | SNV/Stop-gain | C/T | Q112* | 1 | 1 | |
| | MTRR | 25001 | SNV/Stop-gain | C/T | R386* | 1 | 1 | |
| | SARDH | 36568 | SNV/Stop-gain | C/T | R457* | 1 | 1 | |
| | SARDH | 39994 | SNV/Stop-gain | C/T | R514* | 1 | 1 | |
| | SHMT1 | 20155 | SNV/Stop-gain | C/T | R99* | 1 | 1 | |
| | SHMT1 | 28305 | SNV/Stop-gain | C/T | R207* | 1 | 1 | |
| | SHMT1 | 39712 | SNV/Stop-gain | C/G | Y457* | 1 | 1 | |
| | SP8 | 6566 | Deletion/Frameshift | G/- | G147fs | 1 | 0 | Low complexity region |
| | SP8 | 6909 | SNV/Stop-gain | C/A | S261* | 1 | 1 | |
| | DMGDH | 41908 | SNV/Stop-gain | G/A | W495* | 1 | 1 | |
| | BHMT | 24315 | Deletion/Frameshift | A/- | K400fs | 4 | 4 | |
