## Supplementary material for "Accumulation of Rare Coding Variants in Genes Implicated in Risk of Human Cleft Lip with or without Cleft Palate": Table S4

**Table S4.** **Annotations and genotype distributions for all variants scored**

[1] NG refseq coordinate

[2] Exon number based on NG refseq

[3] The mRNA coordinate for all exon variants, or the nearest exon boundary for intronic changes

[4] The distance (upstream (+) or downstream (-)) from the nearest exon boundary

[5] Number of individuals: major allele homozygotes/heterozygotes/minor allele homozygotes

| Gene | Coordinate[1] | Type | Variation | Exon[2] | Exon Coordinate[3] | Distance from Exon[4] | Class | Amino Acid Change | Cases (N) | Case Genotype Counts[5] | Case MAF | Controls (N) | Control Genotype Counts[5] | Control MAF |
|---|---|---|---|---|---|---|---|---|---|---|---|---|---|---|
| AHCY | 21301 | SNV | C/T | 2 | 415 | 0 | Nonsynonymous | R10W | 316 | 309/7/0 | 1.11 | 210 | 201/9/0 | 2.14 |
| AHCY | 21415 | SNV | C/G | 2 | 522 | 7 | intron | | 317 | 317/0/0 | 0.00 | 210 | 209/1/0 | 0.24 |
| AHCY | 22691 | SNV | G/A | 3 | 567 | 0 | Synonymous | A60A | 320 | 319/1/0 | 0.16 | 210 | 210/0/0 | 0.00 |
| AHCY | 22701 | SNV | A/G | 3 | 577 | 0 | Nonsynonymous | I64V | 320 | 319/1/0 | 0.16 | 210 | 210/0/0 | 0.00 |
| AHCY | 24312 | SNV | C/T | 4 | 615 | 0 | Synonymous | G76G | 320 | 319/1/0 | 0.16 | 210 | 210/0/0 | 0.00 |
| AHCY | 24367 | SNV | G/A | 4 | 670 | 0 | Nonsynonymous | G95R | 320 | 316/4/0 | 0.63 | 210 | 210/0/0 | 0.00 |
| AHCY | 25375 | SNV | C/T | 5 | 852 | 0 | Synonymous | S155S | 322 | 322/0/0 | 0.00 | 211 | 210/1/0 | 0.24 |
| AHCY | 25969 | SNV | T/C | 6 | 966 | 0 | Synonymous | Y193Y | 321 | 321/0/0 | 0.00 | 210 | 209/1/0 | 0.24 |
| AHCY | 26050 | SNV | C/T | 6 | 1047 | 0 | Synonymous | N220N | 321 | 320/1/0 | 0.16 | 210 | 207/3/0 | 0.71 |
| AHCY | 31162 | SNV | C/T | 9 | 1276 | -7 | intron | | 320 | 305/13/2 | 2.66 | 210 | 199/10/1 | 2.86 |
| AHCY | 31175 | SNV | C/T | 9 | 1282 | 0 | Nonsynonymous | R299W | 320 | 320/0/0 | 0.00 | 210 | 209/1/0 | 0.24 |
| AHCY | 31329 | SNV | A/G | 9 | 1436 | 0 | Nonsynonymous | K350R | 320 | 319/1/0 | 0.16 | 210 | 210/0/0 | 0.00 |
| AHCYL1 | 29290 | SNV | A/C | 2 | 429 | 0 | Nonsynonymous | D47A | 323 | 322/1/0 | 0.15 | 211 | 211/0/0 | 0.00 |
| AHCYL1 | 32667 | SNV | G/A | 4 | 734 | 0 | Nonsynonymous | V149M | 323 | 323/0/0 | 0.00 | 211 | 210/1/0 | 0.24 |
| AHCYL1 | 35658 | SNV | A/G | 7 | 965 | -4 | intron | | 323 | 323/0/0 | 0.00 | 211 | 210/1/0 | 0.24 |
| AHCYL1 | 37828 | SNV | C/T | 10 | 1341 | 8 | intron | | 323 | 322/1/0 | 0.15 | 211 | 211/0/0 | 0.00 |
| AHCYL1 | 38803 | SNV | T/G | 13 | 1522 | 0 | Synonymous | T411T | 323 | 322/1/0 | 0.15 | 211 | 211/0/0 | 0.00 |
| AHCYL1 | 38806 | SNV | G/A | 13 | 1525 | 0 | Synonymous | P412P | 323 | 323/0/0 | 0.00 | 211 | 210/1/0 | 0.24 |
| AHCYL1 | 38873 | SNV | G/A | 13 | 1592 | 0 | Nonsynonymous | V435I | 323 | 323/0/0 | 0.00 | 211 | 210/1/0 | 0.24 |
| AHCYL1 | 41907 | SNV | A/T | 17 | 1876 | -4 | intron | | 323 | 320/3/0 | 0.46 | 211 | 209/2/0 | 0.47 |
| AHCYL2 | 180299 | SNV | G/A | 6 | 909 | 0 | Synonymous | K282K | 323 | 322/1/0 | 0.15 | 211 | 209/2/0 | 0.47 |
| AHCYL2 | 183380 | SNV | A/C | 7 | 996 | 0 | Synonymous | G311G | 323 | 323/0/0 | 0.00 | 211 | 210/1/0 | 0.24 |
| AHCYL2 | 183444 | SNV | G/A | 7 | 1060 | 0 | Nonsynonymous | V333I | 323 | 323/0/0 | 0.00 | 211 | 210/1/0 | 0.24 |
| AHCYL2 | 185204 | SNV | A/G | 8 | 1205 | 4 | intron | | 323 | 323/0/0 | 0.00 | 211 | 209/2/0 | 0.47 |
| AHCYL2 | 189497 | SNV | G/A | 11 | 1393 | 0 | Nonsynonymous | V444I | 323 | 323/0/0 | 0.00 | 211 | 210/1/0 | 0.24 |
| ALDH1L1 | 24731 | SNV | G/A | 2 | 418 | 0 | Nonsynonymous | G23D | 323 | 322/1/0 | 0.15 | 211 | 211/0/0 | 0.00 |
| ALDH1L1 | 24786 | SNV | C/T | 2 | 473 | 0 | Synonymous | P41P | 323 | 322/1/0 | 0.15 | 211 | 211/0/0 | 0.00 |
| ALDH1L1 | 26995 | SNV | G/A | 3 | 478 | -9 | intron | | 323 | 323/0/0 | 0.00 | 211 | 210/1/0 | 0.24 |
| ALDH1L1 | 27110 | SNV | C/T | 3 | 584 | 0 | Synonymous | A78A | 323 | 322/1/0 | 0.15 | 211 | 211/0/0 | 0.00 |
| ALDH1L1 | 27196 | SNV | C/T | 3 | 670 | 0 | Nonsynonymous | P107L | 323 | 322/1/0 | 0.15 | 211 | 209/2/0 | 0.47 |
| ALDH1L1 | 28138 | SNV | C/A | 4 | 716 | 0 | Synonymous | T122T | 321 | 320/1/0 | 0.16 | 210 | 210/0/0 | 0.00 |
| ALDH1L1 | 28201 | SNV | C/A | 4 | 779 | 0 | Synonymous | T143T | 321 | 318/3/0 | 0.47 | 210 | 208/2/0 | 0.48 |
| ALDH1L1 | 28247 | SNV | A/T | 4 | 825 | 0 | Nonsynonymous | T159S | 321 | 321/0/0 | 0.00 | 210 | 209/1/0 | 0.24 |
| ALDH1L1 | 31023 | SNV | G/A | 6 | 1004 | 0 | Synonymous | E218E | 320 | 319/1/0 | 0.16 | 210 | 210/0/0 | 0.00 |
| ALDH1L1 | 31044 | SNV | C/T | 6 | 1025 | 0 | Synonymous | R225R | 320 | 319/1/0 | 0.16 | 210 | 210/0/0 | 0.00 |
| ALDH1L1 | 32070 | SNV | A/G | 7 | 1079 | 0 | Synonymous | T243T | 323 | 323/0/0 | 0.00 | 211 | 210/1/0 | 0.24 |
| ALDH1L1 | 32102 | SNV | T/C | 7 | 1111 | 0 | Nonsynonymous | L254P | 323 | 260/58/5 | 10.53 | 211 | 183/26/2 | 7.11 |
| ALDH1L1 | 32193 | SNV | C/T | 7 | 1202 | 0 | Synonymous | D284D | 323 | 287/35/1 | 5.73 | 211 | 192/19/0 | 4.50 |
| ALDH1L1 | 32209 | SNV | C/T | 7 | 1208 | 10 | intron | | 323 | 321/2/0 | 0.31 | 211 | 211/0/0 | 0.00 |
| ALDH1L1 | 38720 | SNV | G/T | 9 | 1338 | 0 | Nonsynonymous | V330F | 323 | 227/89/7 | 15.94 | 211 | 152/51/8 | 15.88 |
| ALDH1L1 | 38730 | SNV | G/A | 9 | 1348 | 0 | Nonsynonymous | R333Q | 323 | 322/1/0 | 0.15 | 211 | 211/0/0 | 0.00 |
| ALDH1L1 | 38801 | SNV | G/A | 9 | 1419 | 0 | Nonsynonymous | V357I | 323 | 323/0/0 | 0.00 | 211 | 209/2/0 | 0.47 |
| ALDH1L1 | 47696 | SNV | G/T | 10 | 1440 | 0 | Nonsynonymous | V364L | 323 | 322/1/0 | 0.15 | 211 | 211/0/0 | 0.00 |
| ALDH1L1 | 47758 | SNV | T/C | 10 | 1502 | 0 | Synonymous | F384F | 323 | 322/1/0 | 0.15 | 211 | 209/2/0 | 0.47 |
| ALDH1L1 | 47791 | SNV | G/T | 10 | 1535 | 0 | Synonymous | L395L | 323 | 226/90/7 | 16.10 | 211 | 152/51/8 | 15.88 |
| ALDH1L1 | 48760 | SNV | G/A | 11 | 1575 | 0 | Nonsynonymous | V409M | 322 | 321/1/0 | 0.16 | 211 | 209/2/0 | 0.47 |
| ALDH1L1 | 48815 | SNV | G/C | 11 | 1630 | 0 | Nonsynonymous | G427A | 322 | 322/0/0 | 0.00 | 211 | 210/1/0 | 0.24 |
| ALDH1L1 | 48821 | SNV | A/C | 11 | 1636 | 0 | Nonsynonymous | E429A | 322 | 315/6/1 | 1.24 | 211 | 203/8/0 | 1.90 |
| ALDH1L1 | 48852 | SNV | T/C | 11 | 1667 | 0 | Synonymous | S439S | 322 | 320/2/0 | 0.31 | 211 | 210/1/0 | 0.24 |
| ALDH1L1 | 48887 | SNV | C/A | 11 | 1694 | 8 | intron | | 322 | 232/84/6 | 14.91 | 211 | 159/46/6 | 13.74 |
| ALDH1L1 | 50077 | SNV | A/G | 12 | 1791 | 0 | Nonsynonymous | S481G | 320 | 235/80/5 | 14.06 | 211 | 162/43/6 | 13.03 |
| ALDH1L1 | 50082 | SNV | G/A | 12 | 1796 | 0 | Synonymous | A482A | 320 | 319/1/0 | 0.16 | 210 | 209/1/0 | 0.24 |
| ALDH1L1 | 54105 | SNV | C/T | 13 | 1823 | -4 | intron | | 321 | 104/159/58 | 42.83 | 211 | 71/96/44 | 43.60 |
| ALDH1L1 | 55397 | SNV | C/G | 14 | 2021 | 0 | Synonymous | T557T | 320 | 320/0/0 | 0.00 | 210 | 209/1/0 | 0.24 |
| ALDH1L1 | 55421 | SNV | G/A | 14 | 2044 | 1 | intron | | 321 | 321/0/0 | 0.00 | 210 | 209/1/0 | 0.24 |
| ALDH1L1 | 72853 | SNV | C/G | 19 | 2523 | 0 | Nonsynonymous | R725G | 320 | 319/1/0 | 0.16 | 210 | 210/0/0 | 0.00 |
| ALDH1L1 | 75596 | SNV | G/A | 20 | 2594 | 0 | Synonymous | P748P | 320 | 319/1/0 | 0.16 | 210 | 210/0/0 | 0.00 |
| ALDH1L1 | 75674 | SNV | C/T | 20 | 2672 | 0 | Synonymous | C774C | 320 | 319/1/0 | 0.16 | 210 | 210/0/0 | 0.00 |
| ALDH1L1 | 75686 | SNV | G/A | 20 | 2684 | 0 | Synonymous | Q778Q | 320 | 317/3/0 | 0.47 | 210 | 209/1/0 | 0.24 |
| ALDH1L1 | 75689 | SNV | C/T | 20 | 2687 | 0 | Synonymous | V779V | 320 | 320/0/0 | 0.00 | 210 | 209/1/0 | 0.24 |
| ALDH1L1 | 75704 | SNV | G/A | 20 | 2697 | 5 | intron | | 320 | 318/2/0 | 0.31 | 210 | 210/0/0 | 0.00 |
| ALDH1L1 | 78427 | SNV | A/G | 21 | 2728 | 0 | Nonsynonymous | D793G | 323 | 237/78/8 | 14.55 | 211 | 139/61/11 | 19.67 |
| ALDH1L1 | 78483 | SNV | A/G | 21 | 2784 | 0 | Nonsynonymous | I812V | 323 | 284/38/1 | 6.19 | 211 | 200/10/1 | 2.84 |
| ALDH1L1 | 78508 | SNV | G/A | 21 | 2803 | 6 | intron | | 323 | 319/4/0 | 0.62 | 211 | 211/0/0 | 0.00 |
| ALDH1L1 | 79753 | SNV | C/T | 22 | 2839 | 0 | Nonsynonymous | T830M | 322 | 321/1/0 | 0.16 | 211 | 211/0/0 | 0.00 |
| ALDH1L1 | 79767 | SNV | G/T | 22 | 2853 | 0 | Nonsynonymous | A835S | 322 | 322/0/0 | 0.00 | 211 | 210/1/0 | 0.24 |
| ALDH1L1 | 79880 | SNV | C/T | 22 | 2966 | 0 | Synonymous | F872F | 322 | 321/1/0 | 0.16 | 211 | 211/0/0 | 0.00 |
| ALDH1L2 | 15713 | SNV | T/A | 2 | 320 | 10 | intron | | 323 | 322/1/0 | 0.15 | 211 | 211/0/0 | 0.00 |
| ALDH1L2 | 18844 | SNV | A/G | 3 | 405 | 0 | Nonsynonymous | E93G | 323 | 322/1/0 | 0.15 | 211 | 211/0/0 | 0.00 |

| | | | | | | | | | | | | | | |
|---|---|---|---|---|---|---|---|---|---|---|---|---|---|---|
| ALDH1L2 | 18857 | SNV | C/T | 3 | 418 | 0 | Synonymous | S97S | 323 | 317/6/0 | 0.93 | 211 | 207/4/0 | 0.95 |
| ALDH1L2 | 18932 | SNV | C/T | 3 | 493 | 0 | Synonymous | H122H | 323 | 322/1/0 | 0.15 | 211 | 211/0/0 | 0.00 |
| ALDH1L2 | 26683 | SNV | G/A | 7 | 1048 | 7 | intron | | 323 | 323/0/0 | 0.00 | 211 | 210/1/0 | 0.24 |
| ALDH1L2 | 27876 | SNV | C/T | 8 | 1113 | 0 | Nonsynonymous | T329M | 323 | 322/1/0 | 0.15 | 211 | 211/0/0 | 0.00 |
| ALDH1L2 | 36623 | SNV | C/A | 11 | 1415 | -10 | intron | | 323 | 115/160/48 | 39.63 | 211 | 62/123/26 | 41.47 |
| ALDH1L2 | 36745 | SNV | A/T | 11 | 1527 | 0 | Nonsynonymous | D467V | 323 | 323/0/0 | 0.00 | 211 | 210/1/0 | 0.24 |
| ALDH1L2 | 37361 | SNV | A/G | 12 | 1549 | 0 | Synonymous | V474V | 322 | 321/1/0 | 0.16 | 211 | 211/0/0 | 0.00 |
| ALDH1L2 | 37424 | SNV | C/T | 12 | 1612 | 0 | Synonymous | N495N | 322 | 321/0/1 | 0.31 | 211 | 211/0/0 | 0.00 |
| ALDH1L2 | 42623 | SNV | G/A | 14 | 1842 | 0 | Nonsynonymous | R572H | 323 | 318/4/1 | 0.93 | 211 | 209/2/0 | 0.47 |
| ALDH1L2 | 42637 | SNV | C/G | 14 | 1856 | 0 | Nonsynonymous | L577V | 323 | 322/1/0 | 0.15 | 211 | 211/0/0 | 0.00 |
| ALDH1L2 | 48860 | SNV | T/C | 16 | 1991 | -10 | intron | | 323 | 322/1/0 | 0.15 | 211 | 211/0/0 | 0.00 |
| ALDH1L2 | 48872 | SNV | C/G | 16 | 1993 | 0 | Synonymous | V622V | 323 | 323/0/0 | 0.00 | 211 | 210/1/0 | 0.24 |
| ALDH1L2 | 48875 | SNV | G/A | 16 | 1996 | 0 | Synonymous | T623T | 323 | 146/137/40 | 33.59 | 211 | 88/95/28 | 35.78 |
| ALDH1L2 | 49762 | SNV | C/T | 17 | 2083 | 0 | Synonymous | G652G | 323 | 323/0/0 | 0.00 | 211 | 210/1/0 | 0.24 |
| ALDH1L2 | 49800 | SNV | G/A | 17 | 2121 | 0 | Nonsynonymous | R665H | 323 | 322/1/0 | 0.15 | 211 | 211/0/0 | 0.00 |
| ALDH1L2 | 49819 | SNV | A/T | 17 | 2140 | 0 | Synonymous | G671G | 323 | 146/137/40 | 33.59 | 211 | 87/96/28 | 36.02 |
| ALDH1L2 | 51435 | SNV | T/C | 18 | 2246 | 0 | Nonsynonymous | C707R | 323 | 323/0/0 | 0.00 | 211 | 210/1/0 | 0.24 |
| ALDH1L2 | 51441 | SNV | C/T | 18 | 2252 | 0 | Nonsynonymous | L709F | 323 | 322/1/0 | 0.15 | 211 | 211/0/0 | 0.00 |
| ALDH1L2 | 55217 | SNV | C/T | 19 | 2324 | 0 | Nonsynonymous | R733W | 323 | 322/1/0 | 0.15 | 211 | 211/0/0 | 0.00 |
| ALDH1L2 | 57685 | SNV | A/G | 20 | 2427 | 0 | Nonsynonymous | H767R | 323 | 322/1/0 | 0.15 | 211 | 211/0/0 | 0.00 |
| ALDH1L2 | 57765 | SNV | G/T | 20 | 2507 | 0 | Nonsynonymous | V794L | 323 | 322/1/0 | 0.15 | 211 | 211/0/0 | 0.00 |
| ALDH1L2 | 62985 | SNV | C/T | 22 | 2809 | 0 | Synonymous | G894G | 323 | 323/0/0 | 0.00 | 211 | 210/1/0 | 0.24 |
| ALDH1L2 | 65143 | SNV | C/A | 23 | 2854 | 0 | 3-UTR | | 323 | 322/1/0 | 0.15 | 211 | 211/0/0 | 0.00 |
| AMT | 5257 | SNV | G/A | 1 | 257 | 0 | Nonsynonymous | R10H | 316 | 315/1/0 | 0.16 | 210 | 210/0/0 | 0.00 |
| AMT | 5418 | SNV | G/A | 2 | 329 | 0 | Nonsynonymous | R34H | 316 | 314/2/0 | 0.32 | 210 | 209/1/0 | 0.24 |
| AMT | 5548 | SNV | G/A | 2 | 459 | 0 | Synonymous | S77S | 317 | 316/1/0 | 0.16 | 210 | 210/0/0 | 0.00 |
| AMT | 7351 | SNV | G/A | 4 | 582 | 0 | Synonymous | L118L | 320 | 319/1/0 | 0.16 | 210 | 210/0/0 | 0.00 |
| AMT | 8354 | SNV | G/A | 6 | 859 | 0 | Nonsynonymous | E211K | 320 | 316/4/0 | 0.63 | 210 | 204/6/0 | 1.43 |
| AMT | 8525 | SNV | C/T | 7 | 925 | -3 | intron | | 320 | 320/0/0 | 0.00 | 210 | 209/1/0 | 0.24 |
| AMT | 8583 | SNV | C/G | 7 | 980 | 0 | Nonsynonymous | P251R | 320 | 319/1/0 | 0.16 | 210 | 210/0/0 | 0.00 |
| AMT | 9701 | SNV | T/G | 8 | 1106 | -5 | intron | | 320 | 319/1/0 | 0.16 | 210 | 210/0/0 | 0.00 |
| AMT | 9714 | SNV | C/T | 8 | 1114 | 0 | Nonsynonymous | R296C | 320 | 319/1/0 | 0.16 | 210 | 210/0/0 | 0.00 |
| AMT | 9726 | SNV | A/G | 8 | 1126 | 0 | Nonsynonymous | M300V | 320 | 320/0/0 | 0.00 | 210 | 209/1/0 | 0.24 |
| AMT | 9772 | SNV | G/A | 8 | 1172 | 0 | Nonsynonymous | R315K | 320 | 319/1/0 | 0.16 | 210 | 210/0/0 | 0.00 |
| AMT | 9782 | SNV | G/A | 8 | 1182 | 0 | Synonymous | R318R | 320 | 211/93/16 | 19.53 | 210 | 142/58/10 | 18.57 |
| AMT | 10072 | SNV | G/A | 9 | 1373 | 0 | Nonsynonymous | R382Q | 320 | 319/1/0 | 0.16 | 210 | 210/0/0 | 0.00 |
| AMT | 10114 | SNV | C/T | 9 | 1415 | 0 | Nonsynonymous | P396L | 320 | 319/1/0 | 0.16 | 210 | 210/0/0 | 0.00 |
| ATIC | 5567 | SNV | C/G | 2 | 231 | 0 | Nonsynonymous | T15S | 322 | 321/1/0 | 0.16 | 211 | 210/1/0 | 0.24 |
| ATIC | 18342 | SNV | C/G | 5 | 534 | 0 | Nonsynonymous | T116S | 323 | 134/145/44 | 36.07 | 211 | 86/105/20 | 34.36 |
| ATIC | 19054 | SNV | T/C | 6 | 589 | 0 | Synonymous | A134A | 323 | 323/0/0 | 0.00 | 211 | 210/1/0 | 0.24 |
| ATIC | 19113 | SNV | T/C | 6 | 648 | 0 | Nonsynonymous | V154A | 323 | 322/1/0 | 0.15 | 211 | 211/0/0 | 0.00 |
| ATIC | 19154 | SNV | T/C | 6 | 689 | 0 | Synonymous | L168L | 323 | 323/0/0 | 0.00 | 211 | 210/1/0 | 0.24 |
| ATIC | 19862 | SNV | T/G | 7 | 719 | -5 | intron | | 323 | 323/0/0 | 0.00 | 211 | 210/1/0 | 0.24 |
| ATIC | 19908 | SNV | T/G | 7 | 760 | 0 | Nonsynonymous | D191E | 323 | 322/1/0 | 0.15 | 211 | 211/0/0 | 0.00 |
| ATIC | 19930 | SNV | A/G | 7 | 782 | 0 | Nonsynonymous | K199E | 323 | 323/0/0 | 0.00 | 211 | 210/1/0 | 0.24 |
| ATIC | 19935 | SNV | C/T | 7 | 787 | 0 | Synonymous | G200G | 323 | 322/1/0 | 0.15 | 211 | 211/0/0 | 0.00 |
| ATIC | 25455 | SNV | C/G | 8 | 904 | 0 | Nonsynonymous | N239K | 323 | 323/0/0 | 0.00 | 211 | 210/1/0 | 0.24 |
| ATIC | 31912 | SNV | A/G | 12 | 1374 | 0 | Nonsynonymous | D396G | 323 | 322/1/0 | 0.15 | 211 | 211/0/0 | 0.00 |
| ATIC | 37830 | SNV | G/A | 13 | 1421 | 0 | Nonsynonymous | E412K | 323 | 322/1/0 | 0.15 | 211 | 211/0/0 | 0.00 |
| ATIC | 39849 | SNV | C/T | 14 | 1553 | 0 | Nonsynonymous | R456C | 323 | 323/0/0 | 0.00 | 211 | 210/1/0 | 0.24 |
| ATIC | 39894 | SNV | C/T | 14 | 1598 | 0 | Nonsynonymous | P471S | 323 | 317/6/0 | 0.93 | 211 | 210/1/0 | 0.24 |
| ATIC | 39926 | SNV | A/G | 14 | 1630 | 0 | Synonymous | G481G | 323 | 317/6/0 | 0.93 | 211 | 203/8/0 | 1.90 |
| ATIC | 39977 | SNV | C/G | 14 | 1681 | 0 | Synonymous | T498T | 323 | 319/4/0 | 0.62 | 211 | 209/2/0 | 0.47 |
| ATIC | 39982 | SNV | G/A | 14 | 1686 | 0 | Nonsynonymous | G500D | 323 | 322/1/0 | 0.15 | 211 | 211/0/0 | 0.00 |
| ATIC | 42276 | SNV | C/T | 15 | 1828 | 0 | Synonymous | N547N | 323 | 322/1/0 | 0.15 | 211 | 211/0/0 | 0.00 |
| ATIC | 42636 | SNV | T/C | 16 | 1902 | 0 | Nonsynonymous | I572T | 323 | 323/0/0 | 0.00 | 211 | 210/1/0 | 0.24 |
| ATIC | 42693 | SNV | A/G | 16 | 1959 | 0 | Nonsynonymous | H591R | 323 | 323/0/0 | 0.00 | 211 | 210/1/0 | 0.24 |
| BHMT | 12534 | SNV | C/T | 3 | 327 | 0 | Synonymous | F74F | 323 | 320/3/0 | 0.46 | 211 | 211/0/0 | 0.00 |
| BHMT | 13597 | SNV | G/A | 4 | 418 | 0 | Nonsynonymous | D105N | 323 | 322/1/0 | 0.15 | 211 | 211/0/0 | 0.00 |
| BHMT | 13648 | SNV | G/A | 4 | 469 | 0 | Nonsynonymous | V122M | 323 | 323/0/0 | 0.00 | 211 | 210/1/0 | 0.24 |
| BHMT | 13690 | SNV | G/A | 4 | 511 | 0 | Nonsynonymous | E136K | 323 | 322/1/0 | 0.15 | 211 | 211/0/0 | 0.00 |
| BHMT | 14549 | SNV | C/T | 5 | 694 | 0 | Nonsynonymous | P197S | 323 | 320/3/0 | 0.46 | 211 | 211/0/0 | 0.00 |
| BHMT | 14555 | SNV | G/A | 5 | 700 | 0 | Nonsynonymous | G199S | 323 | 321/2/0 | 0.31 | 211 | 211/0/0 | 0.00 |
| BHMT | 14569 | SNV | G/T | 5 | 714 | 0 | Synonymous | V203V | 323 | 323/0/0 | 0.00 | 211 | 210/1/0 | 0.24 |
| BHMT | 19356 | SNV | G/A | 6 | 821 | 0 | Nonsynonymous | R239Q | 323 | 138/149/36 | 34.21 | 211 | 88/96/27 | 35.55 |
| BHMT | 19432 | SNV | C/T | 6 | 897 | 0 | Synonymous | L264L | 323 | 271/49/3 | 8.51 | 211 | 163/44/4 | 12.32 |
| BHMT | 20976 | SNV | A/C | 7 | 915 | 0 | Synonymous | G270G | 323 | 322/1/0 | 0.15 | 211 | 211/0/0 | 0.00 |
| BHMT | 21018 | SNV | C/T | 7 | 957 | 0 | Synonymous | Y284Y | 323 | 287/34/2 | 5.88 | 211 | 193/18/0 | 4.27 |
| BHMT | 21132 | SNV | A/G | 7 | 1071 | 0 | Synonymous | P322P | 323 | 317/6/0 | 0.93 | 211 | 208/3/0 | 0.71 |
| BHMT | 24315 | Deletion | A/- | 8 | 1305 | 0 | Frameshift | K400fs | 323 | 319/4/0 | 0.62 | 211 | 211/0/0 | 0.00 |
| BHMT2 | 12885 | SNV | T/C | 2 | 221 | 0 | Synonymous | D54D | 323 | 126/151/46 | 37.62 | 211 | 70/99/42 | 43.36 |
| BHMT2 | 14676 | SNV | C/T | 3 | 256 | 0 | Nonsynonymous | A66V | 323 | 315/8/0 | 1.24 | 211 | 207/4/0 | 0.95 |
| BHMT2 | 14695 | SNV | G/A | 3 | 275 | 0 | Synonymous | Q72Q | 323 | 322/1/0 | 0.15 | 211 | 211/0/0 | 0.00 |
| BHMT2 | 16101 | SNV | T/C | 4 | 455 | 0 | Synonymous | F132F | 323 | 322/1/0 | 0.15 | 211 | 211/0/0 | 0.00 |
| BHMT2 | 18226 | SNV | G/A | 5 | 650 | 0 | Synonymous | V197V | 323 | 322/1/0 | 0.15 | 211 | 210/1/0 | 0.24 |
| BHMT2 | 18234 | SNV | G/A | 5 | 657 | 1 | intron | | 323 | 323/0/0 | 0.00 | 211 | 210/1/0 | 0.24 |
| BHMT2 | 18518 | SNV | G/A | 6 | 707 | 0 | Synonymous | L216L | 323 | 321/2/0 | 0.31 | 211 | 206/5/0 | 1.18 |
| BHMT2 | 18569 | SNV | G/T | 6 | 758 | 0 | Synonymous | A233A | 323 | 319/4/0 | 0.62 | 211 | 210/1/0 | 0.24 |
| BHMT2 | 18600 | SNV | G/A | 6 | 789 | 0 | Nonsynonymous | A244T | 323 | 322/1/0 | 0.15 | 211 | 211/0/0 | 0.00 |
| BHMT2 | 18903 | SNV | A/G | 7 | 841 | -2 | intron | | 323 | 322/1/0 | 0.15 | 211 | 211/0/0 | 0.00 |
| BHMT2 | 18908 | SNV | T/C | 7 | 844 | 0 | Nonsynonymous | L262P | 323 | 323/0/0 | 0.00 | 211 | 210/1/0 | 0.24 |
| BHMT2 | 23804 | SNV | G/A | 8 | 1104 | 0 | Nonsynonymous | A349T | 323 | 322/1/0 | 0.15 | 211 | 211/0/0 | 0.00 |

| | | | | | | | | | | | | | | |
|---|---|---|---|---|---|---|---|---|---|---|---|---|---|---|
| BHMT2 | 23814 | SNV | G/A | 8 | 1114 | 0 | Nonsynonymous | R352K | 323 | 322/1/0 | 0.15 | 211 | 211/0/0 | 0.00 |
| BHMT2 | 23817 | SNV | C/A | 8 | 1117 | 0 | Nonsynonymous | P353H | 323 | 323/0/0 | 0.00 | 211 | 210/1/0 | 0.24 |
| BMP4 | 9677 | SNV | G/C | 3 | 537 | 0 | Synonymous | A21A | 320 | 319/1/0 | 0.16 | 210 | 210/0/0 | 0.00 |
| BMP4 | 9782 | SNV | C/A | 3 | 642 | 0 | Nonsynonymous | D56E | 320 | 319/1/0 | 0.16 | 210 | 210/0/0 | 0.00 |
| BMP4 | 9959 | SNV | C/T | 3 | 819 | 0 | Synonymous | N115N | 320 | 320/0/0 | 0.00 | 210 | 209/1/0 | 0.24 |
| BMP4 | 10994 | SNV | G/A | 4 | 890 | 0 | Nonsynonymous | R139H | 320 | 319/1/0 | 0.16 | 210 | 210/0/0 | 0.00 |
| BMP4 | 11033 | SNV | T/C | 4 | 929 | 0 | Nonsynonymous | V152A | 320 | 138/137/45 | 35.47 | 210 | 102/79/29 | 32.62 |
| BMP4 | 11063 | SNV | G/A | 4 | 959 | 0 | Nonsynonymous | R162Q | 320 | 319/1/0 | 0.16 | 210 | 210/0/0 | 0.00 |
| BMP4 | 11246 | SNV | G/A | 4 | 1142 | 0 | Nonsynonymous | R223H | 320 | 319/1/0 | 0.16 | 210 | 210/0/0 | 0.00 |
| BMP4 | 11255 | SNV | G/A | 4 | 1151 | 0 | Nonsynonymous | R226Q | 320 | 319/1/0 | 0.16 | 210 | 210/0/0 | 0.00 |
| BMP4 | 11384 | SNV | G/A | 4 | 1280 | 0 | Nonsynonymous | R269Q | 320 | 319/1/0 | 0.16 | 210 | 210/0/0 | 0.00 |
| BMPR1B | 351515 | SNV | C/T | 4 | 341 | 0 | Nonsynonymous | P23S | 323 | 322/1/0 | 0.15 | 211 | 211/0/0 | 0.00 |
| BMPR1B | 361800 | SNV | T/A | 5 | 474 | 0 | Nonsynonymous | V67D | 323 | 322/1/0 | 0.15 | 211 | 211/0/0 | 0.00 |
| BMPR1B | 362705 | SNV | A/G | 6 | 521 | -4 | intron | | 323 | 322/1/0 | 0.15 | 211 | 211/0/0 | 0.00 |
| BMPR1B | 370902 | SNV | G/A | 7 | 692 | 0 | Nonsynonymous | V140I | 323 | 321/2/0 | 0.31 | 211 | 211/0/0 | 0.00 |
| BMPR1B | 370929 | SNV | C/T | 7 | 719 | 0 | Nonsynonymous | R149W | 323 | 322/1/0 | 0.15 | 211 | 211/0/0 | 0.00 |
| BMPR1B | 372018 | SNV | A/G | 8 | 732 | 0 | Nonsynonymous | Q153R | 323 | 322/1/0 | 0.15 | 211 | 211/0/0 | 0.00 |
| BMPR1B | 377005 | SNV | C/T | 9 | 979 | 0 | Synonymous | T235T | 323 | 321/2/0 | 0.31 | 211 | 211/0/0 | 0.00 |
| BMPR1B | 378361 | SNV | A/G | 10 | 1175 | 0 | Nonsynonymous | M301V | 323 | 322/1/0 | 0.15 | 211 | 211/0/0 | 0.00 |
| BMPR1B | 378525 | SNV | T/A | 10 | 1339 | 0 | Synonymous | V355V | 323 | 322/1/0 | 0.15 | 211 | 211/0/0 | 0.00 |
| BMPR1B | 395807 | SNV | G/A | 11 | 1386 | 0 | Nonsynonymous | R371Q | 323 | 322/1/0 | 0.15 | 211 | 211/0/0 | 0.00 |
| BMPR1B | 395934 | SNV | A/T | 11 | 1513 | 0 | Nonsynonymous | R413S | 323 | 322/1/0 | 0.15 | 211 | 211/0/0 | 0.00 |
| BMPR1B | 401564 | SNV | T/C | 13 | 1658 | -8 | intron | | 323 | 323/0/0 | 0.00 | 211 | 210/1/0 | 0.24 |
| BMPR1B | 401622 | SNV | T/A | 13 | 1708 | 0 | Synonymous | P478P | 323 | 322/1/0 | 0.15 | 211 | 211/0/0 | 0.00 |
| BMPR1B | 401706 | SNV | G/C | 13 | 2066 | 0 | 3-UTR | | 323 | 322/1/0 | 0.15 | 211 | 208/3/0 | 0.71 |
| CBS | 12321 | SNV | A/T | 4 | 460 | 0 | Nonsynonymous | K72I | 316 | 315/1/0 | 0.16 | 207 | 206/1/0 | 0.24 |
| CBS | 12324 | SNV | C/G | 4 | 463 | 0 | Nonsynonymous | S73C | 316 | 316/0/0 | 0.00 | 207 | 206/1/0 | 0.24 |
| CBS | 12327 | SNV | C/T | 4 | 466 | 0 | Nonsynonymous | P74L | 316 | 316/0/0 | 0.00 | 207 | 206/1/0 | 0.24 |
| CBS | 12410 | SNV | A/C | 4 | 549 | 0 | Nonsynonymous | K102Q | 316 | 313/3/0 | 0.47 | 207 | 204/3/0 | 0.72 |
| CBS | 14621 | SNV | G/A | 5 | 629 | 0 | Synonymous | E128E | 316 | 316/0/0 | 0.00 | 210 | 209/1/0 | 0.24 |
| CBS | 14660 | SNV | G/C | 5 | 668 | 0 | Synonymous | T141T | 316 | 315/1/0 | 0.16 | 210 | 209/1/0 | 0.24 |
| CBS | 14666 | SNV | C/T | 5 | 674 | 0 | Synonymous | I143I | 316 | 316/0/0 | 0.00 | 210 | 209/1/0 | 0.24 |
| CBS | 15451 | SNV | G/A | 7 | 818 | 0 | Synonymous | T191T | 316 | 316/0/0 | 0.00 | 210 | 209/1/0 | 0.24 |
| CBS | 15503 | SNV | C/A | 7 | 870 | 0 | Synonymous | R209R | 316 | 315/1/0 | 0.16 | 210 | 210/0/0 | 0.00 |
| CBS | 15514 | SNV | C/T | 7 | 881 | 0 | Synonymous | N212N | 316 | 314/2/0 | 0.32 | 210 | 210/0/0 | 0.00 |
| CBS | 15691 | SNV | C/T | 8 | 944 | 0 | Synonymous | Y233Y | 320 | 199/108/13 | 20.94 | 210 | 122/75/13 | 24.05 |
| CBS | 17857 | SNV | T/C | 10 | 1078 | 0 | Nonsynonymous | I278T | 316 | 315/1/0 | 0.16 | 210 | 210/0/0 | 0.00 |
| CBS | 17963 | SNV | G/A | 10 | 1184 | 0 | Synonymous | T313T | 316 | 316/0/0 | 0.00 | 210 | 209/1/0 | 0.24 |
| CBS | 17977 | SNV | C/T | 10 | 1198 | 0 | Nonsynonymous | T318M | 316 | 315/1/0 | 0.16 | 210 | 210/0/0 | 0.00 |
| CBS | 18623 | SNV | G/A | 11 | 1284 | 3 | intron | | 316 | 316/0/0 | 0.00 | 210 | 209/1/0 | 0.24 |
| CBS | 20404 | SNV | G/A | 12 | 1304 | 0 | Synonymous | T353T | 316 | 315/1/0 | 0.16 | 210 | 208/2/0 | 0.48 |
| CBS | 20425 | SNV | C/T | 12 | 1325 | 0 | Synonymous | A360A | 317 | 185/99/33 | 26.03 | 211 | 124/67/20 | 25.36 |
| CBS | 20497 | SNV | C/T | 12 | 1390 | 7 | intron | | 316 | 315/1/0 | 0.16 | 210 | 210/0/0 | 0.00 |
| CBS | 21957 | SNV | C/T | 14 | 1469 | -6 | intron | | 316 | 315/1/0 | 0.16 | 210 | 210/0/0 | 0.00 |
| CBS | 22011 | SNV | C/T | 14 | 1517 | 0 | Synonymous | T424T | 316 | 315/1/0 | 0.16 | 210 | 208/2/0 | 0.48 |
| CBS | 22012 | SNV | G/A | 14 | 1518 | 0 | Nonsynonymous | V425M | 316 | 315/1/0 | 0.16 | 210 | 210/0/0 | 0.00 |
| CBS | 22019 | SNV | C/T | 14 | 1525 | 0 | Nonsynonymous | P427L | 316 | 315/1/0 | 0.16 | 210 | 210/0/0 | 0.00 |
| CBS | 22668 | SNV | C/T | 15 | 1604 | -10 | intron | | 316 | 315/1/0 | 0.16 | 210 | 210/0/0 | 0.00 |
| CBS | 22730 | SNV | G/A | 15 | 1656 | 0 | Nonsynonymous | G471R | 316 | 315/1/0 | 0.16 | 210 | 210/0/0 | 0.00 |
| CTH | 5159 | SNV | C/A | 1 | 213 | 0 | Nonsynonymous | D5E | 320 | 319/1/0 | 0.16 | 210 | 210/0/0 | 0.00 |
| CTH | 5273 | SNV | G/A | 1 | 327 | 0 | Synonymous | L43L | 320 | 319/1/0 | 0.16 | 210 | 210/0/0 | 0.00 |
| CTH | 9716 | SNV | C/T | 2 | 398 | 0 | Nonsynonymous | T67I | 323 | 320/3/0 | 0.46 | 211 | 208/3/0 | 0.71 |
| CTH | 9723 | SNV | T/C | 2 | 405 | 0 | Synonymous | N69N | 323 | 323/0/0 | 0.00 | 211 | 210/1/0 | 0.24 |
| CTH | 11740 | SNV | T/G | 3 | 523 | 0 | Nonsynonymous | C109G | 323 | 323/0/0 | 0.00 | 211 | 210/1/0 | 0.24 |
| CTH | 11768 | SNV | G/A | 3 | 544 | 7 | intron | | 323 | 321/2/0 | 0.31 | 211 | 211/0/0 | 0.00 |
| CTH | 18053 | SNV | G/A | 5 | 694 | 0 | Nonsynonymous | V166M | 323 | 321/2/0 | 0.31 | 211 | 211/0/0 | 0.00 |
| CTH | 24038 | SNV | G/C | 7 | 845 | -8 | intron | | 323 | 323/0/0 | 0.00 | 211 | 210/1/0 | 0.24 |
| CTH | 25951 | SNV | G/A | 8 | 1062 | 0 | Synonymous | K288K | 323 | 322/1/0 | 0.15 | 211 | 211/0/0 | 0.00 |
| CTH | 27580 | SNV | G/A | 9 | 1099 | 0 | Nonsynonymous | E301K | 323 | 323/0/0 | 0.00 | 211 | 210/1/0 | 0.24 |
| CTH | 32846 | SNV | G/T | 12 | 1406 | 0 | Nonsynonymous | S403I | 323 | 134/146/43 | 35.91 | 211 | 91/99/21 | 33.41 |
| CTNNBIP1 | 43216 | SNV | G/A | 4 | 329 | 0 | Nonsynonymous | G8R | 320 | 319/1/0 | 0.16 | 210 | 210/0/0 | 0.00 |
| CTNNBIP1 | 43227 | SNV | G/A | 4 | 340 | 0 | Synonymous | P11P | 320 | 319/1/0 | 0.16 | 210 | 210/0/0 | 0.00 |
| CTNNBIP1 | 43237 | SNV | T/G | 4 | 350 | 0 | Nonsynonymous | Y15D | 320 | 320/0/0 | 0.00 | 210 | 209/1/0 | 0.24 |
| CTNNBIP1 | 44005 | SNV | C/T | 5 | 427 | 0 | Synonymous | F40F | 316 | 314/2/0 | 0.32 | 210 | 210/0/0 | 0.00 |
| CTNNBIP1 | 64544 | SNV | G/T | 6 | 863 | 0 | 3-UTR | | 320 | 319/1/0 | 0.16 | 210 | 210/0/0 | 0.00 |
| DMGDH | 18712 | SNV | T/G | 3 | 331 | -7 | intron | | 323 | 322/1/0 | 0.15 | 211 | 211/0/0 | 0.00 |
| DMGDH | 18757 | SNV | G/A | 3 | 369 | 0 | Synonymous | L105L | 323 | 322/1/0 | 0.15 | 211 | 211/0/0 | 0.00 |
| DMGDH | 18814 | SNV | G/T | 3 | 426 | 0 | Synonymous | G124G | 323 | 148/145/30 | 31.73 | 211 | 87/92/32 | 36.97 |
| DMGDH | 20322 | SNV | C/G | 4 | 473 | 0 | Nonsynonymous | T140S | 323 | 323/0/0 | 0.00 | 211 | 210/1/0 | 0.24 |
| DMGDH | 20332 | SNV | G/C | 4 | 483 | 0 | Nonsynonymous | R143S | 323 | 322/1/0 | 0.15 | 211 | 211/0/0 | 0.00 |
| DMGDH | 30164 | SNV | C/T | 6 | 889 | 0 | Nonsynonymous | S279P | 323 | 125/159/39 | 36.69 | 211 | 72/97/42 | 42.89 |
| DMGDH | 30193 | SNV | C/G | 6 | 918 | 0 | Synonymous | L288L | 323 | 125/159/39 | 36.69 | 211 | 72/97/42 | 42.89 |
| DMGDH | 30203 | SNV | C/T | 6 | 928 | 0 | Nonsynonymous | R292C | 323 | 322/1/0 | 0.15 | 211 | 211/0/0 | 0.00 |
| DMGDH | 30222 | SNV | A/C | 6 | 947 | 0 | Nonsynonymous | Y298S | 323 | 323/0/0 | 0.00 | 211 | 210/1/0 | 0.24 |
| DMGDH | 30227 | SNV | C/T | 6 | 952 | 0 | Nonsynonymous | L300F | 323 | 322/1/0 | 0.15 | 211 | 209/2/0 | 0.47 |
| DMGDH | 32248 | SNV | A/G | 7 | 1151 | 0 | Nonsynonymous | N366S | 323 | 314/9/0 | 1.39 | 211 | 209/2/0 | 0.47 |
| DMGDH | 32257 | SNV | A/G | 7 | 1160 | 0 | Nonsynonymous | N369S | 323 | 322/1/0 | 0.15 | 211 | 211/0/0 | 0.00 |
| DMGDH | 41372 | SNV | T/C | 8 | 1401 | 0 | Synonymous | Y449Y | 323 | 322/1/0 | 0.15 | 211 | 211/0/0 | 0.00 |
| DMGDH | 41852 | SNV | A/C | 9 | 1483 | 0 | Synonymous | R477R | 323 | 322/1/0 | 0.15 | 211 | 210/1/0 | 0.24 |
| DMGDH | 41890 | SNV | C/T | 9 | 1521 | 0 | Synonymous | G489G | 323 | 237/77/9 | 14.71 | 211 | 156/49/6 | 14.45 |
| DMGDH | 41908 | SNV | G/A | 9 | 1539 | 0 | Truncation | W495* | 323 | 322/1/0 | 0.15 | 211 | 211/0/0 | 0.00 |
| DMGDH | 43625 | SNV | T/A | 10 | 1572 | -4 | intron | | 323 | 321/2/0 | 0.31 | 211 | 211/0/0 | 0.00 |

| | | | | | | | | | | | | | | |
|---|---|---|---|---|---|---|---|---|---|---|---|---|---|---|
| DMGDH | 43700 | SNV | C/G | 10 | 1643 | 0 | Nonsynonymous | A530G | 323 | 208/97/18 | 20.59 | 211 | 125/73/13 | 23.46 |
| DMGDH | 44688 | SNV | A/G | 11 | 1833 | 0 | Synonymous | L593L | 323 | 323/0/0 | 0.00 | 211 | 210/1/0 | 0.24 |
| DMGDH | 45968 | SNV | A/G | 12 | 1869 | -9 | intron | | 323 | 209/96/18 | 20.43 | 211 | 125/74/12 | 23.22 |
| DMGDH | 45996 | SNV | G/A | 12 | 1888 | 0 | Nonsynonymous | V612I | 323 | 322/1/0 | 0.15 | 211 | 211/0/0 | 0.00 |
| DMGDH | 46051 | SNV | T/C | 12 | 1943 | 0 | Nonsynonymous | L630P | 323 | 322/1/0 | 0.15 | 211 | 211/0/0 | 0.00 |
| DMGDH | 46089 | SNV | A/G | 12 | 1981 | 0 | Nonsynonymous | K643E | 323 | 323/0/0 | 0.00 | 211 | 210/1/0 | 0.24 |
| DMGDH | 46098 | SNV | T/C | 12 | 1990 | 0 | Nonsynonymous | S646P | 323 | 209/96/18 | 20.43 | 211 | 125/74/12 | 23.22 |
| DMGDH | 46202 | SNV | T/A | 12 | 2086 | 8 | intron | | 323 | 322/1/0 | 0.15 | 211 | 211/0/0 | 0.00 |
| DMGDH | 50365 | SNV | A/G | 14 | 2304 | 9 | intron | | 323 | 298/24/1 | 4.02 | 211 | 195/15/1 | 4.03 |
| DMGDH | 69230 | SNV | T/C | 15 | 2315 | 0 | Nonsynonymous | F754S | 323 | 321/2/0 | 0.31 | 211 | 211/0/0 | 0.00 |
| DMGDH | 69278 | SNV | G/A | 15 | 2363 | 0 | Nonsynonymous | R770Q | 323 | 322/1/0 | 0.15 | 211 | 209/2/0 | 0.47 |
| DMGDH | 76367 | SNV | G/T | 16 | 2477 | 0 | Nonsynonymous | S808I | 323 | 323/0/0 | 0.00 | 211 | 210/1/0 | 0.24 |
| DMGDH | 76408 | SNV | C/T | 16 | 2518 | 0 | Synonymous | Q821Q | 323 | 322/1/0 | 0.15 | 211 | 211/0/0 | 0.00 |
| DMGDH | 76479 | SNV | A/G | 16 | 2589 | 0 | Synonymous | E845E | 323 | 320/3/0 | 0.46 | 211 | 211/0/0 | 0.00 |
| DMGDH | 76489 | SNV | T/C | 16 | 2599 | 0 | Synonymous | L849L | 323 | 323/0/0 | 0.00 | 211 | 210/1/0 | 0.24 |
| DMGDH | 76535 | SNV | A/C | 16 | 2645 | 0 | Nonsynonymous | D864A | 323 | 323/0/0 | 0.00 | 211 | 210/1/0 | 0.24 |
| DMGDH | 76547 | SNV | A/G | 16 | 2711 | 0 | 3-UTR | | 323 | 322/1/0 | 0.15 | 211 | 211/0/0 | 0.00 |
| FGF10 | 5058 | SNV | T/A | 1 | 58 | 0 | Nonsynonymous | C20S | 323 | 322/1/0 | 0.15 | 211 | 211/0/0 | 0.00 |
| FGF10 | 5097 | SNV | G/A | 1 | 97 | 0 | Nonsynonymous | V33I | 323 | 323/0/0 | 0.00 | 211 | 210/1/0 | 0.24 |
| FGF10 | 5127 | SNV | G/A | 1 | 127 | 0 | Nonsynonymous | D43N | 323 | 323/0/0 | 0.00 | 211 | 210/1/0 | 0.24 |
| FGF10 | 88652 | SNV | C/T | 3 | 591 | 0 | Synonymous | T197T | 323 | 316/7/0 | 1.08 | 211 | 209/2/0 | 0.47 |
| FGF18 | 5778 | SNV | A/G | 2 | 603 | 0 | Synonymous | V22V | 316 | 315/1/0 | 0.16 | 207 | 207/0/0 | 0.00 |
| FGF18 | 41960 | SNV | C/T | 5 | 978 | 0 | Synonymous | S147S | 320 | 320/0/0 | 0.00 | 210 | 209/1/0 | 0.24 |
| FGF18 | 42002 | SNV | G/A | 5 | 1020 | 0 | Synonymous | K161K | 320 | 319/1/0 | 0.16 | 210 | 210/0/0 | 0.00 |
| FGF18 | 42067 | SNV | C/T | 5 | 1085 | 0 | Nonsynonymous | P183L | 320 | 319/1/0 | 0.16 | 210 | 210/0/0 | 0.00 |
| FGF18 | 42068 | SNV | G/A | 5 | 1086 | 0 | Synonymous | P183P | 320 | 236/78/6 | 14.06 | 210 | 166/43/1 | 10.71 |
| FGF3 | 8084 | SNV | G/C | 2 | 794 | 0 | Nonsynonymous | K101N | 316 | 315/1/0 | 0.16 | 210 | 210/0/0 | 0.00 |
| FGFR1 | 16406 | SNV | C/T | 2 | 960 | 0 | Synonymous | C6C | 320 | 319/1/0 | 0.16 | 210 | 210/0/0 | 0.00 |
| FGFR1 | 16463 | SNV | G/A | 2 | 1017 | 0 | Synonymous | P25P | 320 | 320/0/0 | 0.00 | 210 | 209/1/0 | 0.24 |
| FGFR1 | 44068 | SNV | C/T | 3 | 1215 | 0 | Synonymous | S91S | 316 | 315/1/0 | 0.16 | 210 | 210/0/0 | 0.00 |
| FGFR1 | 44099 | SNV | G/A | 3 | 1246 | 0 | Nonsynonymous | V102I | 316 | 315/1/0 | 0.16 | 210 | 209/1/0 | 0.24 |
| FGFR1 | 44115 | SNV | C/T | 3 | 1262 | 0 | Nonsynonymous | S107L | 316 | 315/1/0 | 0.16 | 210 | 210/0/0 | 0.00 |
| FGFR1 | 44140 | SNV | C/T | 3 | 1287 | 0 | Synonymous | S115S | 316 | 315/1/0 | 0.16 | 210 | 210/0/0 | 0.00 |
| FGFR1 | 45733 | SNV | C/G | 2 | 1391 | -9 | intron | | 320 | 319/1/0 | 0.16 | 210 | 210/0/0 | 0.00 |
| FGFR1 | 45893 | SNV | C/T | 5 | 1542 | 0 | Synonymous | D200D | 322 | 312/10/0 | 1.55 | 211 | 207/4/0 | 0.95 |
| FGFR1 | 47720 | SNV | G/A | 2 | 1687 | 7 | intron | | 320 | 319/1/0 | 0.16 | 210 | 210/0/0 | 0.00 |
| FGFR1 | 47722 | SNV | G/A | 2 | 1687 | 9 | intron | | 320 | 319/1/0 | 0.16 | 210 | 210/0/0 | 0.00 |
| FGFR1 | 54116 | SNV | G/A | 9 | 2040 | 0 | Synonymous | P366P | 320 | 319/1/0 | 0.16 | 210 | 210/0/0 | 0.00 |
| FGFR1 | 54239 | SNV | C/T | 9 | 2163 | 0 | Synonymous | D407D | 320 | 320/0/0 | 0.00 | 210 | 209/1/0 | 0.24 |
| FGFR1 | 55545 | SNV | G/T | 10 | 2310 | 0 | Nonsynonymous | M456I | 320 | 320/0/0 | 0.00 | 210 | 209/1/0 | 0.24 |
| FGFR1 | 55575 | SNV | C/T | 10 | 2340 | 0 | Synonymous | P466P | 320 | 320/0/0 | 0.00 | 210 | 209/1/0 | 0.24 |
| FGFR1 | 58967 | SNV | C/T | 14 | 2830 | 0 | Synonymous | L630L | 320 | 316/4/0 | 0.63 | 210 | 210/0/0 | 0.00 |
| FGFR1 | 59692 | SNV | G/A | 2 | 3128 | 9 | intron | | 320 | 320/0/0 | 0.00 | 210 | 209/1/0 | 0.24 |
| FGFR1 | 59806 | SNV | C/T | 2 | 3129 | -6 | intron | | 320 | 314/6/0 | 0.94 | 210 | 209/1/0 | 0.24 |
| FGFR1 | 59887 | SNV | G/A | 17 | 3204 | 0 | Synonymous | L754L | 320 | 314/5/1 | 1.09 | 210 | 206/4/0 | 0.95 |
| FGFR1 | 59903 | SNV | T/C | 17 | 3220 | 0 | Synonymous | L760L | 320 | 319/1/0 | 0.16 | 210 | 210/0/0 | 0.00 |
| FPGS | 6420 | SNV | C/T | 2 | 215 | 0 | Nonsynonymous | R50C | 320 | 319/1/0 | 0.16 | 210 | 210/0/0 | 0.00 |
| FPGS | 6515 | SNV | G/A | 2 | 310 | 0 | Synonymous | L81L | 320 | 318/2/0 | 0.31 | 210 | 208/2/0 | 0.48 |
| FPGS | 6525 | SNV | C/T | 2 | 320 | 0 | Nonsynonymous | R85W | 320 | 320/0/0 | 0.00 | 210 | 209/1/0 | 0.24 |
| FPGS | 6807 | SNV | C/T | 4 | 434 | 0 | Synonymous | L123L | 320 | 319/1/0 | 0.16 | 210 | 210/0/0 | 0.00 |
| FPGS | 9218 | SNV | C/T | 5 | 568 | 5 | intron | | 316 | 316/0/0 | 0.00 | 210 | 209/1/0 | 0.24 |
| FPGS | 13061 | SNV | G/T | 14 | 1355 | -10 | intron | | 317 | 317/0/0 | 0.00 | 210 | 209/1/0 | 0.24 |
| FPGS | 15362 | SNV | C/T | 15 | 1463 | 0 | Nonsynonymous | R466C | 316 | 314/2/0 | 0.32 | 210 | 209/1/0 | 0.24 |
| FPGS | 15432 | SNV | C/T | 15 | 1533 | 0 | Nonsynonymous | A489V | 317 | 317/0/0 | 0.00 | 210 | 209/1/0 | 0.24 |
| FPGS | 15457 | SNV | C/T | 15 | 1558 | 0 | Synonymous | S497S | 316 | 315/1/0 | 0.16 | 210 | 210/0/0 | 0.00 |
| FPGS | 15462 | SNV | C/T | 15 | 1563 | 0 | Nonsynonymous | S499F | 316 | 313/3/0 | 0.47 | 210 | 208/2/0 | 0.48 |
| FPGS | 15464 | SNV | C/T | 15 | 1565 | 0 | Synonymous | L500L | 317 | 317/0/0 | 0.00 | 210 | 209/1/0 | 0.24 |
| FPGS | 15682 | SNV | C/T | 15 | 1783 | 0 | Synonymous | H572H | 316 | 315/1/0 | 0.16 | 210 | 210/0/0 | 0.00 |
| GART | 8624 | SNV | C/T | 2 | 358 | 0 | Nonsynonymous | T16M | 322 | 322/0/0 | 0.00 | 211 | 209/2/0 | 0.47 |
| GART | 8699 | SNV | G/C | 2 | 433 | 0 | Nonsynonymous | C41S | 322 | 322/0/0 | 0.00 | 211 | 210/1/0 | 0.24 |
| GART | 13175 | SNV | T/A | 4 | 588 | 0 | Nonsynonymous | C93I | 321 | 321/0/0 | 0.00 | 209 | 208/1/0 | 0.24 |
| GART | 13176 | SNV | G/T | 4 | 589 | 0 | Nonsynonymous | C93I | 322 | 321/1/0 | 0.16 | 209 | 208/1/0 | 0.24 |
| GART | 13197 | SNV | C/T | 4 | 610 | 0 | Nonsynonymous | A100V | 322 | 321/1/0 | 0.16 | 211 | 211/0/0 | 0.00 |
| GART | 13198 | SNV | G/A | 4 | 611 | 0 | Synonymous | A100A | 322 | 321/1/0 | 0.16 | 211 | 211/0/0 | 0.00 |
| GART | 15557 | SNV | G/A | 5 | 839 | 9 | intron | | 322 | 322/0/0 | 0.00 | 211 | 210/1/0 | 0.24 |
| GART | 16338 | SNV | G/T | 6 | 842 | 0 | Nonsynonymous | E177D | 322 | 321/1/0 | 0.16 | 211 | 211/0/0 | 0.00 |
| GART | 16408 | SNV | T/C | 6 | 908 | 4 | intron | | 322 | 322/0/0 | 0.00 | 211 | 210/1/0 | 0.24 |
| GART | 22918 | SNV | C/G | 11 | 1404 | 0 | Nonsynonymous | L365V | 323 | 322/1/0 | 0.15 | 211 | 211/0/0 | 0.00 |
| GART | 22979 | SNV | G/A | 11 | 1465 | 0 | Nonsynonymous | R385K | 323 | 323/0/0 | 0.00 | 211 | 210/1/0 | 0.24 |
| GART | 23014 | SNV | A/G | 11 | 1500 | 0 | Nonsynonymous | I397V | 323 | 323/0/0 | 0.00 | 211 | 210/1/0 | 0.24 |
| GART | 23036 | SNV | A/G | 11 | 1522 | 0 | Nonsynonymous | K404R | 323 | 322/1/0 | 0.15 | 211 | 211/0/0 | 0.00 |
| GART | 23044 | SNV | C/T | 11 | 1530 | 0 | Synonymous | L407L | 323 | 322/1/0 | 0.15 | 211 | 211/0/0 | 0.00 |
| GART | 23086 | SNV | A/G | 11 | 1572 | 0 | Nonsynonymous | V421I | 323 | 246/74/3 | 12.38 | 211 | 154/49/8 | 15.40 |
| GART | 27355 | SNV | A/G | 14 | 1840 | 0 | Nonsynonymous | D510G | 323 | 321/2/0 | 0.31 | 211 | 211/0/0 | 0.00 |
| GART | 30311 | SNV | T/C | 15 | 2041 | 0 | Nonsynonymous | M577T | 323 | 322/1/0 | 0.15 | 211 | 211/0/0 | 0.00 |
| GART | 30476 | SNV | C/T | 15 | 2206 | 0 | Nonsynonymous | A632V | 323 | 318/5/0 | 0.77 | 211 | 209/2/0 | 0.47 |
| GART | 30502 | SNV | C/G | 15 | 2232 | 0 | Nonsynonymous | P641A | 323 | 322/1/0 | 0.15 | 211 | 211/0/0 | 0.00 |
| GART | 30909 | SNV | A/G | 16 | 2418 | 6 | intron | | 323 | 323/0/0 | 0.00 | 211 | 210/1/0 | 0.24 |
| GART | 36581 | SNV | A/G | 17 | 2566 | 0 | Nonsynonymous | D752G | 323 | 205/114/4 | 18.89 | 211 | 131/72/8 | 20.85 |
| GART | 38068 | SNV | A/C | 18 | 2722 | 0 | Nonsynonymous | E804A | 323 | 321/2/0 | 0.31 | 211 | 210/1/0 | 0.24 |
| GART | 41823 | SNV | G/A | 19 | 2799 | 0 | Nonsynonymous | E830K | 323 | 323/0/0 | 0.00 | 211 | 210/1/0 | 0.24 |
| GART | 42210 | SNV | T/C | 20 | 2915 | 0 | Synonymous | Y868Y | 323 | 322/1/0 | 0.15 | 211 | 211/0/0 | 0.00 |

| | | | | | | | | | | | | | | |
|---|---|---|---|---|---|---|---|---|---|---|---|---|---|---|
| GART | 43370 | SNV | A/T | 21 | 3042 | 0 | Nonsynonymous | M911L | 323 | 322/1/0 | 0.15 | 211 | 211/0/0 | 0.00 |
| GART | 43661 | SNV | G/A | 22 | 3237 | 0 | Nonsynonymous | V976I | 323 | 323/0/0 | 0.00 | 211 | 210/1/0 | 0.24 |
| GART | 43696 | SNV | C/A | 22 | 3272 | 0 | Synonymous | A987A | 323 | 322/1/0 | 0.15 | 211 | 211/0/0 | 0.00 |
| GGH | 8284 | SNV | A/G | 2 | 395 | 0 | Nonsynonymous | I38V | 323 | 322/1/0 | 0.15 | 211 | 211/0/0 | 0.00 |
| GGH | 8346 | SNV | G/A | 2 | 457 | 0 | Synonymous | A58A | 323 | 321/2/0 | 0.31 | 211 | 209/2/0 | 0.47 |
| GGH | 13878 | SNV | A/G | 3 | 551 | 0 | Nonsynonymous | I90V | 323 | 323/0/0 | 0.00 | 211 | 210/1/0 | 0.24 |
| GGH | 13894 | SNV | G/A | 3 | 558 | 9 | intron | | 323 | 159/127/37 | 31.11 | 211 | 84/96/31 | 37.44 |
| GGH | 17756 | SNV | A/G | 5 | 644 | 0 | Nonsynonymous | S121G | 323 | 321/2/0 | 0.31 | 211 | 211/0/0 | 0.00 |
| GGH | 17847 | SNV | C/T | 5 | 735 | 0 | Nonsynonymous | T151I | 323 | 284/38/1 | 6.19 | 211 | 190/21/0 | 4.98 |
| GGH | 17878 | SNV | G/A | 5 | 766 | 0 | Synonymous | P161P | 323 | 322/1/0 | 0.15 | 211 | 211/0/0 | 0.00 |
| GGH | 19935 | Insertion | -/T | 6 | 852 | 0 | Frameshift | L190fs | 323 | 322/1/0 | 0.15 | 211 | 211/0/0 | 0.00 |
| IRF6 | 9822 | SNV | A/C | 3 | 405 | 0 | Nonsynonymous | K34T | 320 | 319/1/0 | 0.16 | 210 | 210/0/0 | 0.00 |
| IRF6 | 14578 | SNV | C/G | 4 | 479 | -5 | intron | | 322 | 173/118/31 | 27.95 | 211 | 118/76/17 | 26.07 |
| IRF6 | 14742 | SNV | C/T | 4 | 638 | 0 | Truncation | Q112* | 323 | 322/1/0 | 0.15 | 211 | 211/0/0 | 0.00 |
| IRF6 | 14759 | SNV | T/G | 4 | 655 | 0 | Synonymous | P117P | 323 | 322/1/0 | 0.15 | 211 | 211/0/0 | 0.00 |
| IRF6 | 15727 | SNV | G/T | 5 | 694 | 0 | Synonymous | G130G | 323 | 322/1/0 | 0.15 | 211 | 209/2/0 | 0.47 |
| IRF6 | 15796 | SNV | G/T | 5 | 763 | 0 | Synonymous | S153S | 323 | 116/141/66 | 42.26 | 211 | 62/87/62 | 50.00 |
| IRF6 | 18720 | SNV | C/T | 6 | 825 | 0 | Nonsynonymous | A174V | 323 | 322/1/0 | 0.15 | 211 | 211/0/0 | 0.00 |
| IRF6 | 18813 | SNV | C/T | 6 | 918 | 0 | Nonsynonymous | A205V | 323 | 323/0/0 | 0.00 | 211 | 210/1/0 | 0.24 |
| IRF6 | 18831 | SNV | A/G | 6 | 936 | 0 | Nonsynonymous | Y211C | 323 | 323/0/0 | 0.00 | 211 | 210/1/0 | 0.24 |
| IRF6 | 20240 | SNV | C/G | 7 | 972 | -8 | intron | | 323 | 322/1/0 | 0.15 | 211 | 211/0/0 | 0.00 |
| IRF6 | 20400 | SNV | G/A | 7 | 1124 | 0 | Nonsynonymous | V274I | 323 | 244/65/14 | 14.40 | 211 | 126/69/16 | 23.93 |
| IRF6 | 20412 | SNV | G/A | 7 | 1136 | 0 | Nonsynonymous | G278S | 323 | 322/1/0 | 0.15 | 211 | 211/0/0 | 0.00 |
| IRF6 | 21442 | SNV | T/C | 8 | 1457 | 0 | Synonymous | L385L | 323 | 317/6/0 | 0.93 | 211 | 207/4/0 | 0.95 |
| LRP6 | 27219 | SNV | C/G | 2 | 198 | -4 | intron | | 323 | 322/1/0 | 0.15 | 211 | 211/0/0 | 0.00 |
| LRP6 | 27230 | SNV | T/A | 2 | 205 | 0 | Synonymous | P21P | 323 | 323/0/0 | 0.00 | 211 | 210/1/0 | 0.24 |
| LRP6 | 27413 | SNV | A/T | 2 | 388 | 0 | Nonsynonymous | K82N | 323 | 322/1/0 | 0.15 | 211 | 211/0/0 | 0.00 |
| LRP6 | 27546 | SNV | T/A | 2 | 521 | 0 | Nonsynonymous | S127T | 323 | 322/1/0 | 0.15 | 211 | 209/2/0 | 0.47 |
| LRP6 | 68607 | SNV | A/G | 3 | 721 | 0 | Synonymous | E193E | 323 | 322/1/0 | 0.15 | 211 | 211/0/0 | 0.00 |
| LRP6 | 87825 | SNV | A/T | 5 | 1045 | 0 | Synonymous | P301P | 323 | 322/1/0 | 0.15 | 211 | 211/0/0 | 0.00 |
| LRP6 | 87846 | SNV | T/C | 5 | 1066 | 0 | Synonymous | C308C | 323 | 246/69/8 | 13.16 | 211 | 161/45/5 | 13.03 |
| LRP6 | 90618 | SNV | C/T | 6 | 1298 | 0 | Nonsynonymous | R386C | 323 | 322/1/0 | 0.15 | 211 | 210/1/0 | 0.24 |
| LRP6 | 90806 | SNV | C/G | 6 | 1486 | 0 | Synonymous | P448P | 323 | 253/63/7 | 11.92 | 211 | 165/42/4 | 11.85 |
| LRP6 | 91970 | SNV | G/A | 7 | 1589 | 0 | Nonsynonymous | V483I | 323 | 317/4/2 | 1.24 | 211 | 203/8/0 | 1.90 |
| LRP6 | 92059 | SNV | C/T | 7 | 1678 | 0 | Synonymous | D512D | 323 | 323/0/0 | 0.00 | 211 | 210/1/0 | 0.24 |
| LRP6 | 106662 | SNV | G/A | 8 | 1767 | 0 | Nonsynonymous | G542D | 323 | 322/1/0 | 0.15 | 211 | 211/0/0 | 0.00 |
| LRP6 | 106684 | SNV | C/T | 8 | 1789 | 0 | Synonymous | D549D | 323 | 323/0/0 | 0.00 | 211 | 210/1/0 | 0.24 |
| LRP6 | 106787 | SNV | C/T | 8 | 1892 | 0 | Nonsynonymous | H584Y | 323 | 323/0/0 | 0.00 | 211 | 210/1/0 | 0.24 |
| LRP6 | 112016 | SNV | C/T | 11 | 2524 | 0 | Synonymous | N794N | 323 | 323/0/0 | 0.00 | 211 | 210/1/0 | 0.24 |
| LRP6 | 112084 | SNV | C/G | 11 | 2592 | 0 | Nonsynonymous | S817C | 323 | 323/0/0 | 0.00 | 211 | 210/1/0 | 0.24 |
| LRP6 | 112964 | SNV | C/T | 12 | 2848 | 0 | Synonymous | H902H | 323 | 321/2/0 | 0.31 | 211 | 210/1/0 | 0.24 |
| LRP6 | 120878 | SNV | A/G | 13 | 2972 | 0 | Nonsynonymous | I944V | 323 | 323/0/0 | 0.00 | 211 | 210/1/0 | 0.24 |
| LRP6 | 120913 | SNV | C/T | 13 | 3007 | 0 | Synonymous | P955P | 323 | 319/4/0 | 0.62 | 211 | 210/1/0 | 0.24 |
| LRP6 | 122860 | SNV | G/A | 14 | 3272 | 0 | Nonsynonymous | D1044N | 323 | 323/0/0 | 0.00 | 211 | 210/1/0 | 0.24 |
| LRP6 | 122914 | SNV | A/G | 14 | 3326 | 0 | Nonsynonymous | V1062I | 323 | 249/71/3 | 11.92 | 211 | 171/34/6 | 10.90 |
| LRP6 | 122922 | SNV | A/G | 14 | 3334 | 0 | Synonymous | V1064V | 323 | 323/0/0 | 0.00 | 211 | 210/1/0 | 0.24 |
| LRP6 | 133531 | SNV | G/A | 16 | 3727 | 0 | Synonymous | K1195K | 323 | 322/1/0 | 0.15 | 211 | 211/0/0 | 0.00 |
| LRP6 | 136586 | SNV | C/G | 17 | 3758 | 0 | Nonsynonymous | P1206A | 323 | 322/1/0 | 0.15 | 211 | 211/0/0 | 0.00 |
| LRP6 | 139897 | SNV | C/T | 18 | 3952 | 0 | Synonymous | C1270C | 323 | 295/27/1 | 4.49 | 211 | 192/18/1 | 4.74 |
| LRP6 | 141064 | SNV | T/C | 19 | 4192 | 0 | Synonymous | D1350D | 323 | 322/1/0 | 0.15 | 211 | 211/0/0 | 0.00 |
| LRP6 | 141076 | SNV | G/C | 19 | 4204 | 0 | Nonsynonymous | K1354N | 323 | 322/1/0 | 0.15 | 211 | 211/0/0 | 0.00 |
| LRP6 | 144995 | SNV | G/A | 20 | 4262 | 0 | Nonsynonymous | V1374I | 323 | 322/1/0 | 0.15 | 211 | 211/0/0 | 0.00 |
| LRP6 | 145019 | SNV | G/T | 20 | 4286 | 0 | Nonsynonymous | V1382F | 323 | 320/3/0 | 0.46 | 211 | 211/0/0 | 0.00 |
| LRP6 | 145024 | SNV | C/G | 20 | 4291 | 0 | Synonymous | T1383T | 323 | 322/1/0 | 0.15 | 211 | 211/0/0 | 0.00 |
| LRP6 | 145077 | SNV | G/A | 20 | 4344 | 0 | Nonsynonymous | R1401H | 323 | 322/1/0 | 0.15 | 211 | 211/0/0 | 0.00 |
| LRP6 | 147208 | Deletion | T/- | 22 | 4592 | -8 | intron | | 323 | 322/1/0 | 0.15 | 210 | 210/0/0 | 0.00 |
| MAT1A | 9081 | SNV | C/G | 2 | 347 | -9 | intron | | 323 | 256/59/8 | 11.61 | 211 | 165/43/3 | 11.61 |
| MAT1A | 10675 | SNV | T/C | 3 | 459 | 0 | Synonymous | C68C | 323 | 322/1/0 | 0.15 | 211 | 211/0/0 | 0.00 |
| MAT1A | 10698 | SNV | T/C | 3 | 482 | 0 | Nonsynonymous | M76T | 323 | 322/1/0 | 0.15 | 211 | 211/0/0 | 0.00 |
| MAT1A | 14356 | SNV | G/A | 5 | 661 | -7 | intron | | 321 | 320/1/0 | 0.16 | 210 | 210/0/0 | 0.00 |
| MAT1A | 14383 | SNV | C/T | 5 | 681 | 0 | Synonymous | A142A | 322 | 213/93/16 | 19.41 | 211 | 137/63/11 | 20.14 |
| MAT1A | 14467 | SNV | C/T | 5 | 765 | 0 | Synonymous | S170S | 321 | 321/0/0 | 0.00 | 210 | 209/1/0 | 0.24 |
| MAT1A | 18124 | SNV | C/G | 6 | 844 | 0 | Nonsynonymous | P197A | 320 | 320/0/0 | 0.00 | 210 | 209/1/0 | 0.24 |
| MAT1A | 19500 | SNV | C/T | 7 | 1044 | 0 | Synonymous | G263G | 320 | 318/2/0 | 0.31 | 210 | 210/0/0 | 0.00 |
| MAT1A | 19579 | SNV | G/A | 7 | 1123 | 0 | Nonsynonymous | V290M | 320 | 320/0/0 | 0.00 | 210 | 209/1/0 | 0.24 |
| MAT1A | 19581 | SNV | G/A | 7 | 1125 | 0 | Synonymous | V290V | 320 | 203/99/18 | 21.09 | 210 | 130/65/15 | 22.62 |
| MAT1A | 19593 | SNV | C/T | 7 | 1137 | 0 | Synonymous | A294A | 321 | 204/99/18 | 21.03 | 211 | 131/65/15 | 22.51 |
| MAT1A | 20079 | SNV | C/T | 8 | 1260 | 0 | Synonymous | Y335Y | 320 | 319/1/0 | 0.16 | 210 | 210/0/0 | 0.00 |
| MAT1A | 20841 | SNV | T/C | 9 | 1386 | 0 | Synonymous | Y377Y | 322 | 87/162/73 | 47.83 | 211 | 56/100/55 | 49.76 |
| MAT2A | 5376 | SNV | A/C | 1 | 376 | 0 | Synonymous | S22S | 316 | 315/1/0 | 0.16 | 210 | 210/0/0 | 0.00 |
| MAT2A | 7122 | SNV | A/G | 2 | 418 | 0 | Synonymous | Q36Q | 323 | 322/1/0 | 0.15 | 211 | 211/0/0 | 0.00 |
| MAT2A | 7385 | SNV | G/A | 3 | 587 | 0 | Nonsynonymous | D93N | 323 | 322/1/0 | 0.15 | 211 | 211/0/0 | 0.00 |
| MAT2A | 7649 | SNV | C/G | 4 | 603 | -7 | intron | | 323 | 323/0/0 | 0.00 | 211 | 210/1/0 | 0.24 |
| MAT2A | 8326 | SNV | C/T | 6 | 1008 | 0 | Nonsynonymous | A233V | 323 | 322/1/0 | 0.15 | 211 | 211/0/0 | 0.00 |
| MAT2A | 8357 | SNV | C/T | 6 | 1039 | 0 | Synonymous | H243H | 323 | 322/1/0 | 0.15 | 211 | 211/0/0 | 0.00 |
| MAT2A | 8611 | SNV | C/G | 7 | 1102 | 0 | Synonymous | R264R | 323 | 100/170/53 | 42.72 | 211 | 83/95/33 | 38.15 |
| MAT2A | 8632 | SNV | T/C | 7 | 1123 | 0 | Synonymous | Y271Y | 323 | 322/1/0 | 0.15 | 211 | 210/1/0 | 0.24 |
| MKX | 15602 | SNV | G/A | 4 | 700 | 0 | Nonsynonymous | V159I | 323 | 320/3/0 | 0.46 | 211 | 207/4/0 | 0.95 |
| MKX | 16091 | SNV | G/A | 5 | 760 | 0 | Nonsynonymous | E179K | 323 | 321/2/0 | 0.31 | 211 | 205/5/0 | 1.42 |
| MKX | 16109 | SNV | C/G | 5 | 778 | 0 | Nonsynonymous | P185A | 323 | 322/1/0 | 0.15 | 211 | 211/0/0 | 0.00 |
| MKX | 16147 | SNV | C/A | 5 | 816 | 0 | Synonymous | V197V | 323 | 322/1/0 | 0.15 | 211 | 211/0/0 | 0.00 |
| MKX | 16309 | SNV | G/T | 5 | 978 | 0 | Synonymous | S251S | 323 | 323/0/0 | 0.00 | 211 | 210/1/0 | 0.24 |

| | | | | | | | | | | | | | | |
|---|---|---|---|---|---|---|---|---|---|---|---|---|---|---|
| MKX | 75309 | SNV | C/T | 6 | 1077 | 0 | Synonymous | N284N | 323 | 165/125/33 | 29.57 | 211 | 97/94/20 | 31.75 |
| MTFMT | 7594 | SNV | A/C | 2 | 236 | -6 | intron | | 323 | 320/3/0 | 0.46 | 211 | 210/1/0 | 0.24 |
| MTFMT | 7625 | SNV | G/A | 2 | 261 | 0 | Nonsynonymous | D79N | 323 | 322/1/0 | 0.15 | 211 | 211/0/0 | 0.00 |
| MTFMT | 7668 | SNV | G/C | 2 | 304 | 0 | Nonsynonymous | G93A | 323 | 323/0/0 | 0.00 | 211 | 210/1/0 | 0.24 |
| MTFMT | 7808 | SNV | T/C | 2 | 444 | 0 | Nonsynonymous | Y140H | 323 | 322/1/0 | 0.15 | 211 | 211/0/0 | 0.00 |
| MTFMT | 7812 | SNV | A/G | 2 | 445 | 3 | intron | | 323 | 323/0/0 | 0.00 | 211 | 210/1/0 | 0.24 |
| MTFMT | 13082 | SNV | G/A | 4 | 627 | 0 | Nonsynonymous | A201T | 323 | 321/2/0 | 0.31 | 211 | 209/2/0 | 0.47 |
| MTFMT | 14363 | Deletion | T/- | 5 | 672 | -5 | intron | | 322 | 307/15/0 | 2.33 | 211 | 202/9/0 | 2.13 |
| MTFMT | 14375 | SNV | C/T | 5 | 679 | 0 | Nonsynonymous | S218L | 322 | 322/0/0 | 0.00 | 211 | 210/1/0 | 0.24 |
| MTFMT | 14391 | SNV | G/T | 5 | 695 | 0 | Nonsynonymous | L223F | 322 | 322/0/0 | 0.00 | 211 | 210/1/0 | 0.24 |
| MTFMT | 18187 | SNV | C/T | 6 | 822 | 0 | Nonsynonymous | R266C | 323 | 322/1/0 | 0.15 | 211 | 211/0/0 | 0.00 |
| MTFMT | 28520 | SNV | C/T | 7 | 911 | 0 | Synonymous | V295V | 323 | 322/1/0 | 0.15 | 211 | 211/0/0 | 0.00 |
| MTFMT | 29717 | SNV | G/A | 8 | 932 | 0 | Synonymous | T302T | 323 | 316/7/0 | 1.08 | 211 | 208/3/0 | 0.71 |
| MTFMT | 31418 | SNV | A/G | 9 | 1036 | 0 | Nonsynonymous | K337R | 323 | 322/1/0 | 0.15 | 211 | 211/0/0 | 0.00 |
| MTFMT | 31473 | SNV | G/A | 9 | 1091 | 0 | Synonymous | Q355Q | 323 | 313/10/0 | 1.55 | 211 | 203/8/0 | 1.90 |
| MTFMT | 31500 | SNV | A/G | 9 | 1118 | 0 | Synonymous | Q364Q | 323 | 322/1/0 | 0.15 | 211 | 211/0/0 | 0.00 |
| MTFMT | 31502 | SNV | G/A | 9 | 1120 | 0 | Nonsynonymous | C365Y | 323 | 320/3/0 | 0.46 | 211 | 211/0/0 | 0.00 |
| MTFMT | 31537 | SNV | A/C | 9 | 1155 | 0 | Nonsynonymous | K377Q | 323 | 322/1/0 | 0.15 | 211 | 211/0/0 | 0.00 |
| MTHFD1 | 17764 | SNV | C/T | 2 | 440 | 0 | Nonsynonymous | A18V | 323 | 320/3/0 | 0.46 | 211 | 211/0/0 | 0.00 |
| MTHFD1 | 29491 | SNV | C/A | 4 | 627 | 6 | intron | | 323 | 321/2/0 | 0.31 | 211 | 211/0/0 | 0.00 |
| MTHFD1 | 32622 | SNV | G/A | 6 | 788 | 0 | Nonsynonymous | K134R | 323 | 252/66/5 | 11.76 | 211 | 148/57/6 | 16.35 |
| MTHFD1 | 32665 | SNV | G/A | 6 | 831 | 0 | Synonymous | T148T | 323 | 323/0/0 | 0.00 | 211 | 210/1/0 | 0.24 |
| MTHFD1 | 42712 | SNV | G/A | 10 | 1265 | 0 | Nonsynonymous | R293H | 323 | 321/2/0 | 0.31 | 211 | 211/0/0 | 0.00 |
| MTHFD1 | 42749 | SNV | T/C | 10 | 1302 | 0 | Synonymous | I305I | 323 | 322/1/0 | 0.15 | 211 | 211/0/0 | 0.00 |
| MTHFD1 | 47198 | SNV | A/G | 13 | 1698 | 8 | intron | | 323 | 318/5/0 | 0.77 | 211 | 208/3/0 | 0.71 |
| MTHFD1 | 52595 | SNV | T/C | 16 | 1948 | 0 | Synonymous | L521L | 323 | 321/2/0 | 0.31 | 211 | 210/1/0 | 0.24 |
| MTHFD1 | 52616 | SNV | A/G | 16 | 1969 | 0 | Nonsynonymous | I528V | 323 | 322/1/0 | 0.15 | 211 | 211/0/0 | 0.00 |
| MTHFD1 | 56109 | SNV | A/G | 17 | 2038 | 0 | Nonsynonymous | T551A | 323 | 323/0/0 | 0.00 | 211 | 210/1/0 | 0.24 |
| MTHFD1 | 59087 | SNV | A/G | 20 | 2345 | 0 | Nonsynonymous | R653Q | 323 | 84/163/76 | 48.76 | 211 | 53/101/57 | 50.95 |
| MTHFD1 | 66407 | SNV | C/T | 24 | 2669 | 0 | Nonsynonymous | T761M | 323 | 322/1/0 | 0.15 | 211 | 209/2/0 | 0.47 |
| MTHFD1 | 66430 | SNV | C/T | 24 | 2692 | 0 | Nonsynonymous | L769F | 323 | 313/10/0 | 1.55 | 211 | 208/3/0 | 0.71 |
| MTHFD1 | 66440 | SNV | G/A | 24 | 2702 | 0 | Nonsynonymous | R772H | 323 | 321/2/0 | 0.31 | 211 | 211/0/0 | 0.00 |
| MTHFD1 | 70814 | SNV | C/T | 25 | 2945 | 0 | Nonsynonymous | T853M | 323 | 322/1/0 | 0.15 | 211 | 211/0/0 | 0.00 |
| MTHFD1L | 15493 | SNV | C/T | 2 | 453 | 0 | Synonymous | I103I | 323 | 322/1/0 | 0.15 | 211 | 211/0/0 | 0.00 |
| MTHFD1L | 16964 | SNV | C/T | 3 | 465 | 0 | Synonymous | D107D | 323 | 323/0/0 | 0.00 | 211 | 210/1/0 | 0.24 |
| MTHFD1L | 16965 | SNV | G/A | 3 | 466 | 0 | Nonsynonymous | D108N | 323 | 322/1/0 | 0.15 | 211 | 211/0/0 | 0.00 |
| MTHFD1L | 17118 | SNV | T/C | 4 | 531 | 0 | Synonymous | I129I | 323 | 322/1/0 | 0.15 | 211 | 211/0/0 | 0.00 |
| MTHFD1L | 22096 | SNV | A/G | 5 | 574 | 0 | Nonsynonymous | I144V | 323 | 323/0/0 | 0.00 | 211 | 210/1/0 | 0.24 |
| MTHFD1L | 57899 | SNV | G/A | 9 | 1040 | 0 | Nonsynonymous | G299E | 323 | 321/2/0 | 0.31 | 211 | 211/0/0 | 0.00 |
| MTHFD1L | 61634 | SNV | G/A | 10 | 1229 | 10 | intron | | 323 | 319/4/0 | 0.62 | 211 | 207/4/0 | 0.95 |
| MTHFD1L | 65485 | SNV | A/G | 11 | 1271 | 0 | Nonsynonymous | D376G | 323 | 321/2/0 | 0.31 | 211 | 209/2/0 | 0.47 |
| MTHFD1L | 65497 | SNV | A/G | 11 | 1283 | 0 | Nonsynonymous | K380R | 323 | 322/1/0 | 0.15 | 211 | 211/0/0 | 0.00 |
| MTHFD1L | 65557 | SNV | G/A | 11 | 1343 | 0 | Nonsynonymous | R400H | 323 | 322/1/0 | 0.15 | 211 | 210/1/0 | 0.24 |
| MTHFD1L | 65624 | SNV | C/T | 11 | 1403 | 7 | intron | | 323 | 322/1/0 | 0.15 | 211 | 211/0/0 | 0.00 |
| MTHFD1L | 65625 | SNV | G/A | 11 | 1403 | 8 | intron | | 322 | 248/67/7 | 12.58 | 211 | 156/52/3 | 13.74 |
| MTHFD1L | 78014 | SNV | T/C | 13 | 1541 | -7 | intron | | 323 | 322/1/0 | 0.15 | 211 | 211/0/0 | 0.00 |
| MTHFD1L | 83870 | SNV | C/T | 14 | 1650 | 0 | Synonymous | A502A | 323 | 322/1/0 | 0.15 | 211 | 210/1/0 | 0.24 |
| MTHFD1L | 83902 | SNV | C/T | 14 | 1682 | 0 | Nonsynonymous | T513M | 323 | 322/1/0 | 0.15 | 211 | 211/0/0 | 0.00 |
| MTHFD1L | 88367 | SNV | T/A | 16 | 1785 | 0 | Nonsynonymous | N547K | 323 | 322/1/0 | 0.15 | 211 | 211/0/0 | 0.00 |
| MTHFD1L | 88417 | SNV | G/A | 16 | 1835 | 0 | Nonsynonymous | R564H | 323 | 319/4/0 | 0.62 | 211 | 208/3/0 | 0.71 |
| MTHFD1L | 95355 | SNV | A/G | 17 | 1912 | 0 | Nonsynonymous | I590V | 322 | 321/1/0 | 0.16 | 211 | 211/0/0 | 0.00 |
| MTHFD1L | 95357 | SNV | C/T | 17 | 1914 | 0 | Synonymous | I590I | 322 | 321/1/0 | 0.16 | 211 | 211/0/0 | 0.00 |
| MTHFD1L | 99643 | SNV | C/A | 18 | 1997 | 0 | Nonsynonymous | A618D | 320 | 319/1/0 | 0.16 | 210 | 210/0/0 | 0.00 |
| MTHFD1L | 111276 | SNV | C/A | 20 | 2168 | 0 | Nonsynonymous | P675H | 323 | 322/1/0 | 0.15 | 211 | 211/0/0 | 0.00 |
| MTHFD1L | 111329 | SNV | G/A | 20 | 2221 | 0 | Nonsynonymous | V693M | 323 | 322/1/0 | 0.15 | 211 | 211/0/0 | 0.00 |
| MTHFD1L | 154193 | SNV | T/C | 23 | 2455 | -9 | intron | | 323 | 322/1/0 | 0.15 | 211 | 211/0/0 | 0.00 |
| MTHFD1L | 154832 | SNV | C/A | 24 | 2556 | -6 | intron | | 323 | 322/1/0 | 0.15 | 211 | 211/0/0 | 0.00 |
| MTHFD1L | 154849 | SNV | G/A | 24 | 2567 | 0 | Nonsynonymous | R808H | 323 | 322/1/0 | 0.15 | 211 | 210/1/0 | 0.24 |
| MTHFD1L | 154925 | SNV | C/G | 24 | 2643 | 0 | Synonymous | S833S | 323 | 99/146/78 | 46.75 | 211 | 62/114/35 | 43.60 |
| MTHFD1L | 154956 | SNV | C/T | 24 | 2674 | 0 | Nonsynonymous | R844W | 323 | 322/1/0 | 0.15 | 211 | 211/0/0 | 0.00 |
| MTHFD1L | 173882 | SNV | C/T | 25 | 2801 | 0 | Nonsynonymous | S886F | 323 | 322/1/0 | 0.15 | 211 | 211/0/0 | 0.00 |
| MTHFD1L | 176382 | SNV | T/A | 26 | 2937 | 0 | Synonymous | P931P | 323 | 321/2/0 | 0.31 | 211 | 210/1/0 | 0.24 |
| MTHFD1L | 176409 | SNV | C/T | 26 | 2964 | 0 | Synonymous | G940G | 323 | 320/3/0 | 0.46 | 211 | 211/0/0 | 0.00 |
| MTHFD2 | 15083 | SNV | A/G | 4 | 565 | 0 | Synonymous | V162V | 323 | 322/1/0 | 0.15 | 211 | 211/0/0 | 0.00 |
| MTHFD2 | 17627 | SNV | T/C | 6 | 750 | -10 | intron | | 323 | 323/0/0 | 0.00 | 211 | 207/4/0 | 0.95 |
| MTHFD2 | 20554 | SNV | T/A | 8 | 1006 | 0 | Synonymous | P309P | 323 | 322/1/0 | 0.15 | 211 | 211/0/0 | 0.00 |
| MTHFR | 7997 | SNV | G/A | 2 | 233 | 0 | Nonsynonymous | E4K | 320 | 319/1/0 | 0.16 | 210 | 210/0/0 | 0.00 |
| MTHFR | 8104 | SNV | C/T | 2 | 340 | 0 | Synonymous | P39P | 320 | 266/51/3 | 8.91 | 210 | 172/35/3 | 9.76 |
| MTHFR | 8226 | SNV | A/G | 2 | 459 | 3 | intron | | 320 | 319/1/0 | 0.16 | 210 | 210/0/0 | 0.00 |
| MTHFR | 9744 | SNV | C/T | 3 | 499 | 0 | Synonymous | D92D | 320 | 319/1/0 | 0.16 | 210 | 210/0/0 | 0.00 |
| MTHFR | 9814 | SNV | G/A | 3 | 569 | 0 | Nonsynonymous | A116T | 320 | 320/0/0 | 0.00 | 210 | 209/1/0 | 0.24 |
| MTHFR | 9816 | SNV | C/T | 3 | 571 | 0 | Synonymous | A116A | 320 | 319/1/0 | 0.16 | 210 | 210/0/0 | 0.00 |
| MTHFR | 9885 | SNV | G/A | 3 | 640 | 0 | Synonymous | T139T | 320 | 310/10/0 | 1.56 | 210 | 205/5/0 | 1.19 |
| MTHFR | 10776 | SNV | C/A | 4 | 699 | -6 | intron | | 320 | 319/1/0 | 0.16 | 210 | 210/0/0 | 0.00 |
| MTHFR | 14783 | SNV | C/T | 5 | 888 | 0 | Nonsynonymous | A222V | 319 | 105/161/53 | 41.85 | 210 | 85/92/33 | 37.62 |
| MTHFR | 14830 | SNV | C/T | 5 | 935 | 0 | Nonsynonymous | R238C | 316 | 315/1/0 | 0.16 | 210 | 210/0/0 | 0.00 |
| MTHFR | 14836 | SNV | G/A | 5 | 941 | 0 | Nonsynonymous | V240M | 316 | 316/0/0 | 0.00 | 210 | 209/1/0 | 0.24 |
| MTHFR | 15978 | SNV | C/T | 6 | 1226 | 0 | Nonsynonymous | R335C | 316 | 316/0/0 | 0.00 | 210 | 209/1/0 | 0.24 |
| MTHFR | 15979 | SNV | G/A | 6 | 1227 | 0 | Nonsynonymous | R335H | 316 | 315/1/0 | 0.16 | 210 | 210/0/0 | 0.00 |
| MTHFR | 16235 | SNV | A/G | 7 | 1255 | -6 | intron | | 320 | 320/0/0 | 0.00 | 210 | 209/1/0 | 0.24 |
| MTHFR | 16265 | SNV | C/T | 7 | 1279 | 0 | Synonymous | S352S | 320 | 262/56/2 | 9.38 | 210 | 172/35/3 | 9.76 |
| MTHFR | 16663 | SNV | G/A | 8 | 1487 | 0 | Nonsynonymous | G422R | 320 | 319/1/0 | 0.16 | 210 | 210/0/0 | 0.00 |

| | | | | | | | | | | | | | | |
|---|---|---|---|---|---|---|---|---|---|---|---|---|---|---|
| MTHFR | 16680 | SNV | T/C | 8 | 1504 | 0 | Synonymous | S427S | 320 | 319/1/0 | 0.16 | 210 | 210/0/0 | 0.00 |
| MTHFR | 16685 | SNV | A/C | 8 | 1509 | 0 | Nonsynonymous | E429A | 320 | 190/115/15 | 22.66 | 210 | 124/76/10 | 22.86 |
| MTHFR | 16704 | SNV | T/C | 8 | 1528 | 0 | Synonymous | F435F | 321 | 303/18/0 | 2.80 | 211 | 202/9/0 | 2.13 |
| MTHFR | 17059 | SNV | C/T | 9 | 1615 | 0 | Synonymous | T464T | 316 | 315/1/0 | 0.16 | 210 | 210/0/0 | 0.00 |
| MTHFR | 17075 | SNV | G/C | 9 | 1631 | 0 | Nonsynonymous | E470L | 316 | 315/1/0 | 0.16 | 210 | 209/1/0 | 0.24 |
| MTHFR | 17143 | SNV | G/A | 9 | 1699 | 0 | Synonymous | P492P | 317 | 317/0/0 | 0.00 | 210 | 209/1/0 | 0.24 |
| MTHFR | 18720 | SNV | C/T | 10 | 1754 | -5 | intron | | 320 | 319/1/0 | 0.16 | 210 | 210/0/0 | 0.00 |
| MTHFR | 18749 | SNV | C/T | 10 | 1778 | 0 | Nonsynonymous | R519C | 320 | 318/2/0 | 0.31 | 210 | 210/0/0 | 0.00 |
| MTHFR | 18750 | SNV | G/T | 10 | 1779 | 0 | Nonsynonymous | R519L | 320 | 319/1/0 | 0.16 | 210 | 210/0/0 | 0.00 |
| MTHFR | 20214 | SNV | C/T | 12 | 1984 | 0 | Synonymous | A587A | 320 | 318/2/0 | 0.31 | 210 | 210/0/0 | 0.00 |
| MTHFR | 20234 | SNV | G/A | 12 | 2004 | 0 | Nonsynonymous | R594Q | 320 | 284/35/1 | 5.78 | 210 | 177/33/0 | 7.86 |
| MTHFR | 20398 | SNV | A/G | 12 | 2168 | 0 | Nonsynonymous | N649D | 320 | 319/1/0 | 0.16 | 210 | 210/0/0 | 0.00 |
| MTHFR | 20411 | SNV | C/T | 12 | 2181 | 0 | Nonsynonymous | T653M | 320 | 311/9/0 | 1.41 | 210 | 207/3/0 | 0.71 |
| MTHFR | 20425 | SNV | C/T | 12 | 2424 | 0 | 3-UTR | | 320 | 319/1/0 | 0.16 | 210 | 210/0/0 | 0.00 |
| MTHFS | 13134 | SNV | C/T | 2 | 328 | 0 | Nonsynonymous | T50I | 323 | 322/1/0 | 0.15 | 211 | 210/1/0 | 0.24 |
| MTHFS | 13162 | SNV | G/C | 2 | 356 | 0 | Nonsynonymous | E59D | 323 | 323/0/0 | 0.00 | 211 | 210/1/0 | 0.24 |
| MTHFS | 13169 | SNV | G/C | 2 | 363 | 0 | Nonsynonymous | V62L | 323 | 323/0/0 | 0.00 | 211 | 210/1/0 | 0.24 |
| MTHFS | 57068 | SNV | A/G | 3 | 612 | 0 | Nonsynonymous | T145A | 323 | 295/28/0 | 4.33 | 211 | 195/16/0 | 3.79 |
| MTR | 5418 | SNV | G/C | 1 | 841 | 0 | 5-UTR | | 317 | 317/0/0 | 0.00 | 210 | 209/1/0 | 0.24 |
| MTR | 13268 | SNV | G/A | 2 | 578 | 0 | Nonsynonymous | R52Q | 323 | 320/3/0 | 0.46 | 211 | 208/3/0 | 0.71 |
| MTR | 15857 | SNV | G/A | 3 | 673 | -7 | intron | | 323 | 305/18/0 | 2.79 | 211 | 196/15/0 | 3.55 |
| MTR | 18418 | SNV | C/T | 4 | 763 | -6 | intron | | 323 | 304/19/0 | 2.94 | 211 | 199/12/0 | 2.84 |
| MTR | 25373 | SNV | G/A | 7 | 1082 | 0 | Nonsynonymous | R220Q | 323 | 322/1/0 | 0.15 | 211 | 211/0/0 | 0.00 |
| MTR | 26210 | SNV | C/T | 8 | 1134 | 0 | Synonymous | S237S | 323 | 323/0/0 | 0.00 | 211 | 210/1/0 | 0.24 |
| MTR | 26241 | SNV | G/A | 8 | 1165 | 0 | Nonsynonymous | V248M | 323 | 322/1/0 | 0.15 | 211 | 210/1/0 | 0.24 |
| MTR | 33932 | SNV | C/T | 9 | 1281 | 0 | Synonymous | P286P | 323 | 321/2/0 | 0.31 | 211 | 210/1/0 | 0.24 |
| MTR | 36561 | SNV | G/A | 11 | 1363 | 0 | Nonsynonymous | D314N | 323 | 306/17/0 | 2.63 | 211 | 203/8/0 | 1.90 |
| MTR | 38952 | SNV | C/T | 12 | 1462 | 0 | Nonsynonymous | P347S | 323 | 323/0/0 | 0.00 | 211 | 210/1/0 | 0.24 |
| MTR | 41678 | SNV | C/T | 13 | 1499 | -8 | intron | | 323 | 323/0/0 | 0.00 | 211 | 209/2/0 | 0.47 |
| MTR | 45413 | SNV | A/G | 14 | 1752 | 6 | intron | | 323 | 322/1/0 | 0.15 | 211 | 211/0/0 | 0.00 |
| MTR | 60080 | SNV | C/A | 16 | 1955 | 0 | Nonsynonymous | T511K | 323 | 322/1/0 | 0.15 | 211 | 211/0/0 | 0.00 |
| MTR | 69576 | SNV | A/G | 19 | 2400 | 0 | Synonymous | K659K | 323 | 323/0/0 | 0.00 | 211 | 210/1/0 | 0.24 |
| MTR | 73218 | SNV | C/T | 22 | 2772 | 0 | Synonymous | D783D | 323 | 322/1/0 | 0.15 | 211 | 211/0/0 | 0.00 |
| MTR | 83568 | SNV | G/T | 23 | 2896 | 8 | intron | | 323 | 322/1/0 | 0.15 | 211 | 211/0/0 | 0.00 |
| MTR | 94867 | SNV | A/G | 26 | 3126 | 0 | Synonymous | L901L | 323 | 322/1/0 | 0.15 | 211 | 209/2/0 | 0.47 |
| MTR | 94908 | SNV | A/T | 26 | 3167 | 0 | Nonsynonymous | D915V | 323 | 322/1/0 | 0.15 | 211 | 211/0/0 | 0.00 |
| MTR | 94920 | SNV | A/G | 26 | 3179 | 0 | Nonsynonymous | D919G | 323 | 215/97/11 | 18.42 | 211 | 135/68/8 | 19.91 |
| MTR | 96049 | SNV | G/T | 27 | 3236 | 0 | Nonsynonymous | S938I | 323 | 322/1/0 | 0.15 | 211 | 210/1/0 | 0.24 |
| MTR | 96051 | SNV | G/C | 27 | 3238 | 0 | Nonsynonymous | G939R | 323 | 321/2/0 | 0.31 | 211 | 211/0/0 | 0.00 |
| MTR | 98951 | SNV | A/G | 28 | 3325 | 0 | Nonsynonymous | K968E | 323 | 322/1/0 | 0.15 | 211 | 211/0/0 | 0.00 |
| MTR | 100880 | SNV | A/T | 29 | 3458 | 0 | Nonsynonymous | D1012V | 323 | 322/1/0 | 0.15 | 211 | 210/1/0 | 0.24 |
| MTR | 100924 | SNV | C/T | 29 | 3502 | 0 | Nonsynonymous | R1027W | 323 | 322/1/0 | 0.15 | 211 | 210/1/0 | 0.24 |
| MTR | 100986 | SNV | C/T | 29 | 3564 | 0 | Synonymous | Y1047Y | 323 | 323/0/0 | 0.00 | 211 | 210/1/0 | 0.24 |
| MTR | 100989 | SNV | A/G | 29 | 3567 | 0 | Synonymous | A1048A | 323 | 101/153/69 | 45.05 | 211 | 57/109/45 | 47.16 |
| MTR | 104178 | SNV | T/A | 30 | 3729 | 0 | Synonymous | V1102V | 323 | 323/0/0 | 0.00 | 211 | 210/1/0 | 0.24 |
| MTR | 104255 | SNV | C/T | 30 | 3806 | 0 | Nonsynonymous | A1128V | 323 | 322/1/0 | 0.15 | 211 | 211/0/0 | 0.00 |
| MTR | 105146 | SNV | G/A | 31 | 3897 | 0 | Synonymous | L1158L | 321 | 320/1/0 | 0.16 | 210 | 209/1/0 | 0.24 |
| MTR | 105149 | SNV | C/T | 31 | 3900 | 0 | Synonymous | D1159D | 321 | 320/1/0 | 0.16 | 211 | 209/2/0 | 0.47 |
| MTR | 105163 | SNV | G/A | 31 | 3914 | 0 | Nonsynonymous | R1164H | 321 | 319/2/0 | 0.31 | 210 | 209/1/0 | 0.24 |
| MTR | 105164 | SNV | C/A | 31 | 3915 | 0 | Synonymous | R1164R | 322 | 101/154/67 | 44.72 | 211 | 57/109/45 | 47.16 |
| MTR | 105168 | SNV | C/T | 31 | 3919 | 0 | Synonymous | L1166L | 321 | 307/13/1 | 2.34 | 210 | 202/8/0 | 1.90 |
| MTR | 105248 | SNV | T/C | 31 | 3999 | 0 | Synonymous | L1192L | 323 | 126/142/55 | 39.01 | 211 | 87/101/23 | 34.83 |
| MTR | 106716 | SNV | C/A | 32 | 4022 | -10 | intron | | 323 | 322/1/0 | 0.15 | 211 | 211/0/0 | 0.00 |
| MTR | 106722 | SNV | C/G | 32 | 4022 | -4 | intron | | 323 | 322/1/0 | 0.15 | 211 | 211/0/0 | 0.00 |
| MTR | 106792 | SNV | A/G | 32 | 4088 | 0 | Nonsynonymous | N1222S | 323 | 321/2/0 | 0.31 | 211 | 211/0/0 | 0.00 |
| MTR | 107270 | SNV | T/C | 33 | 4135 | -8 | intron | | 323 | 280/39/4 | 7.28 | 211 | 183/27/1 | 6.87 |
| MTR | 107271 | SNV | T/G | 33 | 4135 | -7 | intron | | 323 | 321/2/0 | 0.31 | 211 | 211/0/0 | 0.00 |
| MTRR | 6718 | SNV | T/C | 2 | 164 | 0 | Synonymous | A9A | 323 | 322/1/0 | 0.15 | 211 | 211/0/0 | 0.00 |
| MTRR | 6745 | SNV | C/T | 2 | 191 | 0 | Synonymous | I18I | 323 | 322/1/0 | 0.15 | 211 | 211/0/0 | 0.00 |
| MTRR | 6757 | SNV | A/G | 2 | 203 | 0 | Nonsynonymous | I22M | 323 | 154/139/30 | 30.80 | 211 | 98/99/14 | 30.09 |
| MTRR | 6797 | SNV | C/T | 2 | 243 | 0 | Nonsynonymous | H36Y | 323 | 322/1/0 | 0.15 | 211 | 211/0/0 | 0.00 |
| MTRR | 11159 | SNV | C/T | 4 | 425 | 0 | Synonymous | L96L | 323 | 322/1/0 | 0.15 | 211 | 211/0/0 | 0.00 |
| MTRR | 13902 | SNV | A/C | 5 | 600 | 0 | Synonymous | R155R | 323 | 319/4/0 | 0.62 | 211 | 211/0/0 | 0.00 |
| MTRR | 13963 | SNV | C/T | 5 | 661 | 0 | Nonsynonymous | S175L | 323 | 184/121/18 | 24.30 | 211 | 125/68/18 | 24.64 |
| MTRR | 13976 | SNV | T/C | 5 | 674 | 0 | Synonymous | L179L | 323 | 168/134/21 | 27.24 | 211 | 106/80/25 | 30.81 |
| MTRR | 14093 | SNV | T/G | 5 | 791 | 0 | Nonsynonymous | N218K | 323 | 322/1/0 | 0.15 | 211 | 211/0/0 | 0.00 |
| MTRR | 14117 | SNV | C/T | 5 | 815 | 0 | Synonymous | S226S | 323 | 323/0/0 | 0.00 | 211 | 210/1/0 | 0.24 |
| MTRR | 14208 | SNV | T/A | 5 | 906 | 0 | Nonsynonymous | S257T | 323 | 278/44/1 | 7.12 | 211 | 185/26/0 | 6.16 |
| MTRR | 19065 | SNV | T/G | 6 | 931 | 0 | Nonsynonymous | V265G | 323 | 322/1/0 | 0.15 | 211 | 211/0/0 | 0.00 |
| MTRR | 19128 | SNV | C/T | 6 | 994 | 0 | Nonsynonymous | T286M | 323 | 320/3/0 | 0.46 | 211 | 211/0/0 | 0.00 |
| MTRR | 19140 | SNV | T/C | 6 | 1006 | 0 | Nonsynonymous | I290T | 323 | 322/1/0 | 0.15 | 211 | 211/0/0 | 0.00 |
| MTRR | 19147 | SNV | C/T | 6 | 1013 | 0 | Synonymous | T292T | 323 | 323/0/0 | 0.00 | 211 | 210/1/0 | 0.24 |
| MTRR | 21691 | SNV | C/G | 7 | 1134 | 0 | Nonsynonymous | L333V | 323 | 314/9/0 | 1.39 | 211 | 204/7/0 | 1.66 |
| MTRR | 21714 | SNV | C/T | 7 | 1157 | 0 | Synonymous | C340C | 323 | 322/1/0 | 0.15 | 211 | 211/0/0 | 0.00 |
| MTRR | 21743 | SNV | A/G | 7 | 1186 | 0 | Nonsynonymous | K350R | 323 | 168/134/21 | 27.24 | 211 | 106/81/24 | 30.57 |
| MTRR | 21750 | SNV | A/C | 7 | 1193 | 0 | Nonsynonymous | K352N | 323 | 322/1/0 | 0.15 | 211 | 211/0/0 | 0.00 |
| MTRR | 25000 | SNV | G/A | 9 | 1292 | 0 | Synonymous | L385L | 323 | 249/67/7 | 12.54 | 211 | 170/38/3 | 10.43 |
| MTRR | 25001 | SNV | C/T | 9 | 1293 | 0 | Truncation | R386* | 323 | 322/1/0 | 0.15 | 211 | 211/0/0 | 0.00 |
| MTRR | 25088 | SNV | C/T | 9 | 1380 | 0 | Nonsynonymous | R415C | 323 | 249/67/7 | 12.54 | 211 | 170/38/3 | 10.43 |
| MTRR | 25098 | SNV | G/A | 9 | 1390 | 0 | Nonsynonymous | R418Q | 323 | 322/1/0 | 0.15 | 211 | 211/0/0 | 0.00 |
| MTRR | 27290 | SNV | C/G | 10 | 1486 | 0 | Nonsynonymous | P450R | 323 | 251/65/7 | 12.23 | 211 | 174/34/3 | 9.48 |
| MTRR | 28717 | SNV | A/G | 11 | 1601 | 0 | Synonymous | V488V | 323 | 318/5/0 | 0.77 | 211 | 206/5/0 | 1.18 |

| | | | | | | | | | | | | | | |
|---|---|---|---|---|---|---|---|---|---|---|---|---|---|---|
| MTRR | 28721 | SNV | A/G | 11 | 1605 | 0 | Nonsynonymous | T490A | 323 | 321/2/0 | 0.31 | 211 | 210/1/0 | 0.24 |
| MTRR | 28789 | SNV | C/T | 11 | 1673 | 0 | Synonymous | S512S | 323 | 318/5/0 | 0.77 | 211 | 206/5/0 | 1.18 |
| MTRR | 31726 | SNV | G/A | 12 | 1790 | 0 | Synonymous | P551P | 323 | 319/4/0 | 0.62 | 211 | 210/1/0 | 0.24 |
| MTRR | 31759 | SNV | T/C | 12 | 1813 | 10 | intron | | 323 | 322/1/0 | 0.15 | 211 | 210/1/0 | 0.24 |
| MTRR | 32752 | Insertion | -/T | 13 | 1814 | -9 | intron | | 323 | 321/2/0 | 0.31 | 211 | 210/1/0 | 0.24 |
| MTRR | 32760 | SNV | G/A | 13 | 1814 | -1 | intron | | 323 | 322/1/0 | 0.15 | 211 | 211/0/0 | 0.00 |
| MTRR | 32791 | SNV | A/C | 13 | 1844 | 0 | Synonymous | G569G | 323 | 323/0/0 | 0.00 | 211 | 210/1/0 | 0.24 |
| MTRR | 32845 | SNV | T/C | 13 | 1898 | 0 | Synonymous | Y587Y | 323 | 318/5/0 | 0.77 | 211 | 206/5/0 | 1.18 |
| MTRR | 32975 | SNV | C/T | 14 | 1920 | 0 | Nonsynonymous | H595Y | 323 | 176/127/20 | 25.85 | 211 | 108/82/21 | 29.38 |
| MTRR | 33011 | SNV | G/A | 14 | 1956 | 0 | Nonsynonymous | V607I | 323 | 323/0/0 | 0.00 | 211 | 210/1/0 | 0.24 |
| MTRR | 33067 | SNV | G/A | 14 | 2012 | 0 | Synonymous | V625V | 323 | 173/130/20 | 26.32 | 211 | 106/82/23 | 30.33 |
| MTRR | 33103 | SNV | G/A | 14 | 2048 | 0 | Synonymous | A637A | 323 | 185/120/18 | 24.15 | 211 | 124/70/17 | 24.64 |
| MTRR | 35879 | SNV | T/C | 15 | 2158 | 0 | Nonsynonymous | V674A | 323 | 322/1/0 | 0.15 | 211 | 211/0/0 | 0.00 |
| PTCH1 | 15377 | SNV | C/T | 2 | 400 | 0 | Nonsynonymous | T71I | 317 | 317/0/0 | 0.00 | 210 | 209/1/0 | 0.24 |
| PTCH1 | 15483 | SNV | C/T | 2 | 506 | 0 | Synonymous | L106L | 316 | 306/10/0 | 1.58 | 210 | 206/4/0 | 0.95 |
| PTCH1 | 15549 | SNV | G/C | 2 | 572 | 0 | Synonymous | L128L | 316 | 315/1/0 | 0.16 | 210 | 210/0/0 | 0.00 |
| PTCH1 | 36264 | SNV | T/C | 3 | 755 | 0 | Synonymous | H189H | 323 | 321/2/0 | 0.31 | 211 | 210/1/0 | 0.24 |
| PTCH1 | 40006 | SNV | A/G | 5 | 923 | 0 | Synonymous | T245T | 323 | 315/8/0 | 1.24 | 211 | 206/5/0 | 1.18 |
| PTCH1 | 41377 | SNV | G/C | 6 | 935 | -1 | intron | | 322 | 321/1/0 | 0.16 | 211 | 211/0/0 | 0.00 |
| PTCH1 | 42825 | SNV | T/C | 8 | 1262 | 0 | Synonymous | H358H | 323 | 322/1/0 | 0.15 | 211 | 211/0/0 | 0.00 |
| PTCH1 | 42849 | SNV | A/G | 8 | 1286 | 0 | Synonymous | L366L | 323 | 322/1/0 | 0.15 | 211 | 211/0/0 | 0.00 |
| PTCH1 | 42870 | SNV | C/T | 8 | 1307 | 0 | Synonymous | Y373Y | 323 | 322/1/0 | 0.15 | 211 | 211/0/0 | 0.00 |
| PTCH1 | 42871 | SNV | G/C | 8 | 1308 | 0 | Nonsynonymous | E374Q | 323 | 323/0/0 | 0.00 | 211 | 210/1/0 | 0.24 |
| PTCH1 | 42888 | SNV | C/T | 8 | 1325 | 0 | Synonymous | Y379Y | 323 | 322/1/0 | 0.15 | 211 | 211/0/0 | 0.00 |
| PTCH1 | 43774 | SNV | C/A | 9 | 1404 | -6 | intron | | 323 | 322/1/0 | 0.15 | 211 | 211/0/0 | 0.00 |
| PTCH1 | 43811 | SNV | C/G | 9 | 1435 | 0 | Nonsynonymous | T416S | 323 | 322/1/0 | 0.15 | 211 | 211/0/0 | 0.00 |
| PTCH1 | 43870 | SNV | G/A | 9 | 1494 | 0 | Nonsynonymous | D436N | 323 | 323/0/0 | 0.00 | 211 | 210/1/0 | 0.24 |
| PTCH1 | 45101 | SNV | T/C | 11 | 1692 | -8 | intron | | 323 | 306/16/1 | 2.79 | 211 | 196/13/2 | 4.03 |
| PTCH1 | 45845 | SNV | C/T | 12 | 1829 | 0 | Synonymous | S547S | 323 | 320/3/0 | 0.46 | 211 | 209/2/0 | 0.47 |
| PTCH1 | 45869 | SNV | T/C | 12 | 1853 | 0 | Synonymous | N555N | 323 | 280/43/0 | 6.66 | 211 | 189/22/0 | 5.21 |
| PTCH1 | 45890 | SNV | C/T | 12 | 1874 | 0 | Synonymous | A562A | 323 | 256/63/4 | 10.99 | 211 | 154/53/4 | 14.45 |
| PTCH1 | 45914 | SNV | G/T | 12 | 1898 | 0 | Synonymous | L570L | 323 | 322/1/0 | 0.15 | 211 | 211/0/0 | 0.00 |
| PTCH1 | 52819 | SNV | C/T | 14 | 2042 | 0 | Synonymous | C618C | 320 | 320/0/0 | 0.00 | 210 | 209/1/0 | 0.24 |
| PTCH1 | 52878 | SNV | G/A | 14 | 2101 | 0 | Nonsynonymous | R638H | 320 | 319/1/0 | 0.16 | 210 | 210/0/0 | 0.00 |
| PTCH1 | 53138 | SNV | C/T | 14 | 2361 | 0 | Nonsynonymous | P725S | 320 | 319/1/0 | 0.16 | 210 | 209/1/0 | 0.24 |
| PTCH1 | 53148 | SNV | C/T | 14 | 2371 | 0 | Nonsynonymous | T728M | 320 | 319/1/0 | 0.16 | 210 | 210/0/0 | 0.00 |
| PTCH1 | 53164 | SNV | A/G | 14 | 2387 | 0 | Synonymous | S733S | 320 | 310/10/0 | 1.56 | 210 | 208/2/0 | 0.48 |
| PTCH1 | 54594 | SNV | C/T | 15 | 2492 | 0 | Synonymous | T768T | 321 | 320/1/0 | 0.16 | 210 | 210/0/0 | 0.00 |
| PTCH1 | 54719 | SNV | C/T | 15 | 2617 | 0 | Nonsynonymous | A810V | 321 | 321/0/0 | 0.00 | 210 | 209/1/0 | 0.24 |
| PTCH1 | 54775 | SNV | G/A | 15 | 2673 | 0 | Nonsynonymous | V829M | 320 | 320/0/0 | 0.00 | 210 | 209/1/0 | 0.24 |
| PTCH1 | 54859 | SNV | G/C | 15 | 2748 | 9 | intron | | 319 | 187/111/21 | 23.98 | 211 | 119/74/18 | 26.07 |
| PTCH1 | 60087 | SNV | G/A | 16 | 2868 | 0 | Nonsynonymous | D894N | 320 | 320/0/0 | 0.00 | 210 | 209/1/0 | 0.24 |
| PTCH1 | 60099 | SNV | G/A | 16 | 2880 | 0 | Nonsynonymous | D898N | 320 | 319/1/0 | 0.16 | 210 | 210/0/0 | 0.00 |
| PTCH1 | 62198 | SNV | C/A | 17 | 2907 | 0 | Nonsynonymous | L907M | 320 | 319/1/0 | 0.16 | 210 | 210/0/0 | 0.00 |
| PTCH1 | 62308 | SNV | A/G | 17 | 3017 | 0 | Synonymous | P943P | 320 | 319/1/0 | 0.16 | 210 | 210/0/0 | 0.00 |
| PTCH1 | 63698 | SNV | T/C | 18 | 3101 | 0 | Synonymous | Y971Y | 320 | 318/2/0 | 0.31 | 210 | 209/1/0 | 0.24 |
| PTCH1 | 63722 | SNV | C/T | 18 | 3125 | 0 | Synonymous | N979N | 320 | 318/2/0 | 0.31 | 210 | 210/0/0 | 0.00 |
| PTCH1 | 63745 | SNV | T/C | 18 | 3148 | 0 | Nonsynonymous | F987S | 320 | 320/0/0 | 0.00 | 210 | 209/1/0 | 0.24 |
| PTCH1 | 63774 | SNV | A/G | 18 | 3177 | 0 | Nonsynonymous | I997V | 320 | 319/1/0 | 0.16 | 210 | 210/0/0 | 0.00 |
| PTCH1 | 63791 | SNV | G/A | 18 | 3194 | 0 | Synonymous | T1002T | 320 | 320/0/0 | 0.00 | 210 | 209/1/0 | 0.24 |
| PTCH1 | 63926 | SNV | T/G | 18 | 3329 | 0 | Synonymous | L1047L | 320 | 311/9/0 | 1.41 | 210 | 202/8/0 | 1.90 |
| PTCH1 | 63940 | SNV | C/T | 18 | 3343 | 0 | Nonsynonymous | T1052M | 320 | 320/0/0 | 0.00 | 210 | 209/1/0 | 0.24 |
| PTCH1 | 65548 | SNV | T/C | 19 | 3357 | -5 | intron | | 320 | 319/1/0 | 0.16 | 210 | 210/0/0 | 0.00 |
| PTCH1 | 68426 | SNV | C/T | 20 | 3575 | 0 | Synonymous | G1129G | 320 | 318/2/0 | 0.31 | 210 | 210/0/0 | 0.00 |
| PTCH1 | 72063 | SNV | G/A | 21 | 3675 | 0 | Nonsynonymous | G1163S | 323 | 322/1/0 | 0.15 | 211 | 211/0/0 | 0.00 |
| PTCH1 | 72117 | SNV | T/C | 21 | 3729 | 0 | Nonsynonymous | Y1181H | 323 | 322/1/0 | 0.15 | 211 | 211/0/0 | 0.00 |
| PTCH1 | 72660 | SNV | C/T | 22 | 3755 | 0 | Synonymous | G1189G | 316 | 314/2/0 | 0.32 | 210 | 209/1/0 | 0.24 |
| PTCH1 | 72676 | SNV | A/T | 22 | 3771 | 0 | Nonsynonymous | T1195S | 316 | 305/11/0 | 1.74 | 210 | 197/11/2 | 3.57 |
| PTCH1 | 72699 | SNV | C/T | 22 | 3794 | 0 | Synonymous | P1202P | 316 | 314/2/0 | 0.32 | 210 | 210/0/0 | 0.00 |
| PTCH1 | 72727 | SNV | G/A | 22 | 3822 | 0 | Nonsynonymous | G1212S | 316 | 315/1/0 | 0.16 | 210 | 210/0/0 | 0.00 |
| PTCH1 | 72741 | SNV | C/T | 22 | 3836 | 0 | Synonymous | S1216S | 316 | 315/1/0 | 0.16 | 210 | 210/0/0 | 0.00 |
| PTCH1 | 74555 | SNV | C/T | 23 | 4033 | 0 | Nonsynonymous | P1282L | 316 | 314/2/0 | 0.32 | 210 | 209/1/0 | 0.24 |
| PTCH1 | 74618 | SNV | G/A | 23 | 4096 | 0 | Nonsynonymous | R1303H | 316 | 315/1/0 | 0.16 | 210 | 209/1/0 | 0.24 |
| PTCH1 | 74654 | SNV | C/T | 23 | 4132 | 0 | Nonsynonymous | P1315L | 317 | 90/154/73 | 47.32 | 211 | 72/90/49 | 44.55 |
| PTCH1 | 74666 | SNV | G/A | 23 | 4144 | 0 | Nonsynonymous | R1319H | 316 | 315/1/0 | 0.16 | 210 | 210/0/0 | 0.00 |
| PTCH1 | 74737 | SNV | G/A | 23 | 4215 | 0 | Nonsynonymous | G1343R | 316 | 316/0/0 | 0.00 | 210 | 209/1/0 | 0.24 |
| PTCH1 | 74743 | SNV | C/G | 23 | 4221 | 0 | Nonsynonymous | R1345G | 316 | 315/1/0 | 0.16 | 210 | 210/0/0 | 0.00 |
| PTCH1 | 74790 | SNV | C/T | 23 | 4268 | 0 | Synonymous | S1360S | 316 | 314/2/0 | 0.32 | 210 | 209/1/0 | 0.24 |
| PTCH1 | 74847 | SNV | C/T | 23 | 4325 | 0 | Synonymous | V1379V | 316 | 315/1/0 | 0.16 | 210 | 210/0/0 | 0.00 |
| PTCH1 | 74925 | SNV | C/T | 23 | 4403 | 0 | Synonymous | D1405D | 316 | 315/1/0 | 0.16 | 210 | 210/0/0 | 0.00 |
| PTCH1 | 74962 | SNV | G/A | 23 | 4440 | 0 | Nonsynonymous | V1418I | 316 | 314/2/0 | 0.32 | 210 | 210/0/0 | 0.00 |
| PTCH1 | 74984 | SNV | C/T | 23 | 4462 | 0 | Nonsynonymous | S1425L | 317 | 317/0/0 | 0.00 | 210 | 209/1/0 | 0.24 |
| PTCH1 | 75034 | SNV | C/T | 23 | 4512 | 0 | Nonsynonymous | R1442W | 316 | 315/1/0 | 0.16 | 210 | 210/0/0 | 0.00 |
| PTCH1 | 75035 | SNV | G/A | 23 | 4513 | 0 | Nonsynonymous | R1442Q | 316 | 315/1/0 | 0.16 | 210 | 210/0/0 | 0.00 |
| NECTIN1 | 55145 | SNV | C/T | 2 | 436 | 0 | Synonymous | S88S | 316 | 315/1/0 | 0.16 | 207 | 207/0/0 | 0.00 |
| NECTIN1 | 55304 | SNV | G/A | 2 | 595 | 0 | Synonymous | T141T | 316 | 315/1/0 | 0.16 | 207 | 206/1/0 | 0.24 |
| NECTIN1 | 56034 | SNV | G/A | 3 | 768 | 0 | Nonsynonymous | R199Q | 320 | 315/5/0 | 0.78 | 210 | 209/1/0 | 0.24 |
| NECTIN1 | 56056 | SNV | G/A | 3 | 790 | 0 | Synonymous | T206T | 320 | 320/0/0 | 0.00 | 210 | 209/1/0 | 0.24 |
| NECTIN1 | 56073 | SNV | G/A | 3 | 807 | 0 | Nonsynonymous | R212H | 320 | 319/1/0 | 0.16 | 210 | 210/0/0 | 0.00 |
| NECTIN1 | 56565 | SNV | C/A | 4 | 964 | 0 | Nonsynonymous | D264E | 320 | 320/0/0 | 0.00 | 210 | 209/1/0 | 0.24 |
| NECTIN1 | 56579 | SNV | G/A | 4 | 978 | 0 | Nonsynonymous | C269Y | 320 | 319/1/0 | 0.16 | 210 | 210/0/0 | 0.00 |
| NECTIN1 | 56608 | SNV | G/A | 4 | 1007 | 0 | Nonsynonymous | E279K | 320 | 320/0/0 | 0.00 | 210 | 209/1/0 | 0.24 |

| | | | | | | | | | | | | | | |
|---|---|---|---|---|---|---|---|---|---|---|---|---|---|---|
| NECTIN1 | 58552 | SNV | G/A | 5 | 1160 | 0 | Nonsynonymous | E330K | 320 | 316/4/0 | 0.63 | 210 | 208/2/0 | 0.48 |
| SARDH | 10835 | SNV | G/A | 2 | 346 | 0 | Nonsynonymous | S18N | 317 | 317/0/0 | 0.00 | 210 | 209/1/0 | 0.24 |
| SARDH | 10846 | SNV | G/T | 2 | 357 | 0 | Nonsynonymous | G22C | 316 | 314/2/0 | 0.32 | 210 | 208/2/0 | 0.48 |
| SARDH | 10912 | SNV | C/T | 2 | 423 | 0 | Nonsynonymous | R44W | 316 | 314/2/0 | 0.32 | 210 | 210/0/0 | 0.00 |
| SARDH | 10928 | SNV | G/A | 2 | 439 | 0 | Nonsynonymous | G49E | 316 | 315/1/0 | 0.16 | 210 | 210/0/0 | 0.00 |
| SARDH | 10932 | SNV | G/A | 2 | 443 | 0 | Synonymous | Q50Q | 317 | 168/130/19 | 26.50 | 211 | 119/81/11 | 24.41 |
| SARDH | 10986 | SNV | C/T | 2 | 497 | 0 | Synonymous | N68N | 316 | 316/0/0 | 0.00 | 210 | 209/1/0 | 0.24 |
| SARDH | 11090 | SNV | C/T | 2 | 601 | 0 | Nonsynonymous | S103F | 316 | 315/1/0 | 0.16 | 210 | 210/0/0 | 0.00 |
| SARDH | 12348 | SNV | C/T | 3 | 625 | -7 | intron | | 316 | 316/0/0 | 0.00 | 210 | 209/1/0 | 0.24 |
| SARDH | 12390 | SNV | G/T | 3 | 660 | 0 | Nonsynonymous | V123L | 316 | 315/1/0 | 0.16 | 210 | 210/0/0 | 0.00 |
| SARDH | 12410 | SNV | T/A | 3 | 680 | 0 | Synonymous | T129T | 316 | 316/0/0 | 0.00 | 210 | 209/1/0 | 0.24 |
| SARDH | 12496 | SNV | A/G | 3 | 766 | 0 | Nonsynonymous | N158S | 316 | 315/1/0 | 0.16 | 210 | 208/2/0 | 0.48 |
| SARDH | 12528 | SNV | A/G | 3 | 798 | 0 | Nonsynonymous | M169V | 316 | 316/0/0 | 0.00 | 210 | 209/1/0 | 0.24 |
| SARDH | 13527 | SNV | C/T | 4 | 859 | 0 | Nonsynonymous | T189I | 320 | 317/3/0 | 0.47 | 210 | 210/0/0 | 0.00 |
| SARDH | 13556 | SNV | C/T | 4 | 888 | 0 | Nonsynonymous | L199F | 320 | 320/0/0 | 0.00 | 210 | 209/1/0 | 0.24 |
| SARDH | 13561 | SNV | C/T | 4 | 893 | 0 | Synonymous | Y200Y | 320 | 318/2/0 | 0.31 | 210 | 210/0/0 | 0.00 |
| SARDH | 13580 | SNV | C/A | 4 | 912 | 0 | Nonsynonymous | H207N | 320 | 319/1/0 | 0.16 | 210 | 209/1/0 | 0.24 |
| SARDH | 13600 | SNV | C/T | 4 | 932 | 0 | Synonymous | P213P | 320 | 320/0/0 | 0.00 | 210 | 209/1/0 | 0.24 |
| SARDH | 14825 | SNV | G/A | 5 | 1040 | 0 | Synonymous | V249V | 321 | 320/1/0 | 0.16 | 210 | 209/1/0 | 0.24 |
| SARDH | 14897 | SNV | G/A | 5 | 1107 | 5 | intron | | 321 | 320/1/0 | 0.16 | 210 | 210/0/0 | 0.00 |
| SARDH | 15086 | SNV | C/T | 6 | 1108 | -5 | intron | | 320 | 319/1/0 | 0.16 | 210 | 210/0/0 | 0.00 |
| SARDH | 15120 | SNV | G/T | 6 | 1137 | 0 | Nonsynonymous | A282S | 320 | 319/1/0 | 0.16 | 210 | 210/0/0 | 0.00 |
| SARDH | 15164 | SNV | C/T | 6 | 1181 | 0 | Synonymous | V296V | 320 | 319/1/0 | 0.16 | 210 | 210/0/0 | 0.00 |
| SARDH | 15165 | SNV | G/A | 6 | 1182 | 0 | Nonsynonymous | V297I | 320 | 318/2/0 | 0.31 | 210 | 209/1/0 | 0.24 |
| SARDH | 25946 | SNV | T/C | 7 | 1241 | 0 | Synonymous | S316S | 320 | 320/0/0 | 0.00 | 210 | 208/2/0 | 0.48 |
| SARDH | 25996 | SNV | C/T | 7 | 1291 | 0 | Nonsynonymous | A333V | 320 | 320/0/0 | 0.00 | 210 | 208/2/0 | 0.48 |
| SARDH | 27497 | SNV | G/A | 8 | 1314 | -4 | intron | | 320 | 319/1/0 | 0.16 | 210 | 210/0/0 | 0.00 |
| SARDH | 27596 | SNV | G/T | 8 | 1409 | 0 | Nonsynonymous | E372D | 320 | 314/6/0 | 0.94 | 210 | 206/4/0 | 0.95 |
| SARDH | 31841 | SNV | C/T | 9 | 1453 | 0 | Nonsynonymous | T387M | 316 | 315/1/0 | 0.16 | 210 | 210/0/0 | 0.00 |
| SARDH | 36568 | SNV | C/T | 11 | 1662 | 0 | Truncation | R457* | 316 | 315/1/0 | 0.16 | 210 | 210/0/0 | 0.00 |
| SARDH | 36624 | SNV | T/C | 11 | 1718 | 0 | Synonymous | D475D | 317 | 317/0/0 | 0.00 | 210 | 209/1/0 | 0.24 |
| SARDH | 36641 | SNV | G/A | 11 | 1735 | 0 | Nonsynonymous | R481H | 316 | 315/1/0 | 0.16 | 210 | 210/0/0 | 0.00 |
| SARDH | 36666 | SNV | C/T | 11 | 1760 | 0 | Synonymous | H489H | 316 | 94/157/65 | 45.41 | 210 | 72/96/42 | 42.86 |
| SARDH | 36667 | SNV | G/A | 11 | 1761 | 0 | Nonsynonymous | E490K | 316 | 316/0/0 | 0.00 | 210 | 209/1/0 | 0.24 |
| SARDH | 36675 | SNV | G/A | 11 | 1763 | 6 | intron | | 316 | 315/1/0 | 0.16 | 210 | 210/0/0 | 0.00 |
| SARDH | 39917 | SNV | C/T | 12 | 1764 | -8 | intron | | 316 | 313/2/1 | 0.63 | 210 | 209/1/0 | 0.24 |
| SARDH | 39994 | SNV | C/T | 12 | 1833 | 0 | Truncation | R514* | 316 | 315/1/0 | 0.16 | 210 | 210/0/0 | 0.00 |
| SARDH | 48630 | SNV | C/T | 14 | 1997 | 0 | Synonymous | A568A | 320 | 318/2/0 | 0.31 | 210 | 208/2/0 | 0.48 |
| SARDH | 48696 | SNV | C/T | 14 | 2063 | 0 | Synonymous | A590A | 320 | 319/1/0 | 0.16 | 210 | 210/0/0 | 0.00 |
| SARDH | 48711 | SNV | C/T | 14 | 2078 | 0 | Synonymous | S595S | 320 | 191/115/14 | 22.34 | 210 | 129/64/17 | 23.33 |
| SARDH | 54433 | SNV | C/T | 16 | 2219 | 0 | Synonymous | D642D | 316 | 316/0/0 | 0.00 | 210 | 209/1/0 | 0.24 |
| SARDH | 54449 | SNV | A/G | 16 | 2235 | 0 | Nonsynonymous | M648V | 318 | 91/152/75 | 47.48 | 210 | 60/98/52 | 48.10 |
| SARDH | 59740 | SNV | C/G | 17 | 2433 | 0 | Nonsynonymous | L714V | 316 | 315/1/0 | 0.16 | 210 | 210/0/0 | 0.00 |
| SARDH | 81046 | SNV | G/T | 21 | 3029 | 0 | Nonsynonymous | K912N | 320 | 319/1/0 | 0.16 | 210 | 210/0/0 | 0.00 |
| SHMT1 | 12555 | SNV | C/A | 2 | 416 | 0 | 5-UTR | | 320 | 319/1/0 | 0.16 | 210 | 210/0/0 | 0.00 |
| SHMT1 | 12563 | SNV | T/G | 2 | 213 | 0 | Nonsynonymous | M1? | 320 | 318/2/0 | 0.31 | 210 | 209/1/0 | 0.24 |
| SHMT1 | 14878 | SNV | G/A | 3 | 453 | 7 | intron | | 323 | 227/86/10 | 16.41 | 211 | 151/50/10 | 16.59 |
| SHMT1 | 14881 | SNV | T/C | 3 | 453 | 10 | intron | | 322 | 317/5/0 | 0.78 | 211 | 210/1/0 | 0.24 |
| SHMT1 | 20155 | SNV | C/T | 4 | 506 | 0 | Truncation | R99* | 320 | 319/1/0 | 0.16 | 210 | 210/0/0 | 0.00 |
| SHMT1 | 21017 | SNV | G/A | 5 | 700 | 0 | Synonymous | T163T | 322 | 322/0/0 | 0.00 | 211 | 210/1/0 | 0.24 |
| SHMT1 | 27724 | SNV | C/G | 6 | 731 | -6 | intron | | 320 | 319/1/0 | 0.16 | 210 | 210/0/0 | 0.00 |
| SHMT1 | 27800 | SNV | T/C | 6 | 801 | 0 | Nonsynonymous | L197P | 320 | 319/1/0 | 0.16 | 210 | 210/0/0 | 0.00 |
| SHMT1 | 28305 | SNV | C/T | 7 | 830 | 0 | Truncation | R207* | 320 | 319/1/0 | 0.16 | 210 | 210/0/0 | 0.00 |
| SHMT1 | 28323 | SNV | C/T | 7 | 848 | 0 | Nonsynonymous | R213W | 320 | 319/1/0 | 0.16 | 210 | 210/0/0 | 0.00 |
| SHMT1 | 28329 | SNV | C/T | 7 | 854 | 0 | Nonsynonymous | R215W | 320 | 319/1/0 | 0.16 | 210 | 210/0/0 | 0.00 |
| SHMT1 | 28333 | SNV | A/G | 7 | 858 | 0 | Nonsynonymous | K216R | 320 | 314/6/0 | 0.94 | 210 | 206/3/1 | 1.19 |
| SHMT1 | 28439 | SNV | G/C | 7 | 964 | 0 | Synonymous | V251V | 320 | 317/3/0 | 0.47 | 210 | 208/2/0 | 0.48 |
| SHMT1 | 28454 | SNV | C/T | 7 | 979 | 0 | Synonymous | H256H | 320 | 319/1/0 | 0.16 | 210 | 210/0/0 | 0.00 |
| SHMT1 | 32865 | SNV | C/T | 8 | 1026 | -3 | intron | | 322 | 322/0/0 | 0.00 | 211 | 210/1/0 | 0.24 |
| SHMT1 | 35383 | SNV | G/A | 9 | 1265 | 6 | intron | | 322 | 322/0/0 | 0.00 | 211 | 210/1/0 | 0.24 |
| SHMT1 | 37908 | SNV | G/A | 10 | 1302 | 0 | Nonsynonymous | R364H | 320 | 319/1/0 | 0.16 | 210 | 210/0/0 | 0.00 |
| SHMT1 | 39712 | SNV | C/G | 12 | 1582 | 0 | Truncation | Y457* | 317 | 317/0/0 | 0.00 | 210 | 209/1/0 | 0.24 |
| SHMT1 | 39721 | SNV | C/T | 12 | 1591 | 0 | Synonymous | A460A | 316 | 313/3/0 | 0.47 | 210 | 208/2/0 | 0.48 |
| SHMT1 | 39761 | SNV | C/T | 12 | 1631 | 0 | Nonsynonymous | L474F | 318 | 184/115/19 | 24.06 | 210 | 126/74/10 | 22.38 |
| SHMT2 | 6346 | SNV | C/T | 2 | 355 | 0 | Nonsynonymous | S50L | 320 | 311/9/0 | 1.41 | 210 | 206/4/0 | 0.95 |
| SHMT2 | 6347 | SNV | G/A | 2 | 356 | 0 | Synonymous | S50S | 320 | 320/0/0 | 0.00 | 210 | 209/1/0 | 0.24 |
| SHMT2 | 7282 | SNV | C/T | 4 | 659 | 0 | Synonymous | Y151Y | 320 | 319/1/0 | 0.16 | 210 | 210/0/0 | 0.00 |
| SHMT2 | 7330 | SNV | C/T | 4 | 707 | 0 | Synonymous | P167P | 320 | 320/0/0 | 0.00 | 210 | 209/1/0 | 0.24 |
| SHMT2 | 7663 | SNV | C/T | 5 | 743 | 0 | Synonymous | D179D | 320 | 312/8/0 | 1.25 | 210 | 203/7/0 | 1.67 |
| SHMT2 | 7720 | SNV | C/T | 5 | 800 | 0 | Synonymous | N198N | 320 | 320/0/0 | 0.00 | 210 | 209/1/0 | 0.24 |
| SHMT2 | 7906 | SNV | A/G | 6 | 826 | 0 | Nonsynonymous | N207S | 320 | 319/1/0 | 0.16 | 210 | 210/0/0 | 0.00 |
| SHMT2 | 8154 | SNV | A/G | 7 | 946 | 0 | Nonsynonymous | H247R | 320 | 320/0/0 | 0.00 | 210 | 209/1/0 | 0.24 |
| SHMT2 | 8219 | SNV | A/G | 7 | 1011 | 0 | Nonsynonymous | K269E | 320 | 320/0/0 | 0.00 | 210 | 209/1/0 | 0.24 |
| SHMT2 | 8227 | SNV | G/A | 7 | 1019 | 0 | Synonymous | A271A | 320 | 301/19/0 | 2.97 | 210 | 200/10/0 | 2.38 |
| SHMT2 | 8234 | SNV | G/A | 7 | 1026 | 0 | Nonsynonymous | V274I | 320 | 319/1/0 | 0.16 | 210 | 210/0/0 | 0.00 |
| SHMT2 | 8719 | SNV | G/T | 8 | 1175 | 0 | Synonymous | L323L | 320 | 282/35/3 | 6.41 | 211 | 188/19/4 | 6.40 |
| SHMT2 | 8982 | SNV | C/T | 9 | 1230 | -9 | intron | | 320 | 319/1/0 | 0.16 | 210 | 210/0/0 | 0.00 |
| SHMT2 | 9170 | SNV | A/G | 10 | 1330 | -7 | intron | | 320 | 312/8/0 | 1.25 | 210 | 204/6/0 | 1.43 |
| SHMT2 | 9172 | Deletion | C/- | 10 | 1330 | -5 | intron | | 320 | 319/1/0 | 0.16 | 210 | 210/0/0 | 0.00 |
| SHMT2 | 9433 | SNV | G/A | 11 | 1488 | 0 | Nonsynonymous | A428T | 322 | 320/2/0 | 0.31 | 211 | 211/0/0 | 0.00 |
| SHMT2 | 9461 | SNV | G/A | 11 | 1516 | 0 | Nonsynonymous | R437H | 322 | 322/0/0 | 0.00 | 211 | 210/1/0 | 0.24 |
| SHMT2 | 9715 | SNV | C/T | 12 | 1647 | 0 | Nonsynonymous | R481C | 322 | 321/1/0 | 0.16 | 211 | 211/0/0 | 0.00 |

| | | | | | | | | | | | | | | |
|---|---|---|---|---|---|---|---|---|---|---|---|---|---|---|
| SHMT2 | 9716 | SNV | G/A | 12 | 1648 | 0 | Nonsynonymous | R481H | 322 | 320/2/0 | 0.31 | 211 | 211/0/0 | 0.00 |
| SP8 | 6909 | SNV | C/A | 3 | 3038 | 0 | Truncation | S261* | 316 | 315/1/0 | 0.15 | 208 | 208/0/0 | 0.00 |
| TFAP2A | 22010 | SNV | T/C | 5 | 922 | 0 | Synonymous | G269G | 323 | 323/0/0 | 0.00 | 211 | 210/1/0 | 0.24 |
| TFAP2A | 24010 | SNV | C/T | 6 | 1020 | 0 | Synonymous | Y302Y | 323 | 321/2/0 | 0.31 | 211 | 210/1/0 | 0.24 |
| TFAP2A | 24011 | SNV | G/A | 6 | 1021 | 0 | Nonsynonymous | V303M | 323 | 323/0/0 | 0.00 | 211 | 210/1/0 | 0.24 |
| TFAP2A | 26022 | SNV | G/T | 7 | 1290 | 0 | Synonymous | T392T | 316 | 315/1/0 | 0.16 | 210 | 208/2/0 | 0.48 |
| TFAP2A | 26091 | SNV | C/T | 7 | 1359 | 0 | Synonymous | N415N | 317 | 259/53/5 | 9.94 | 210 | 177/32/1 | 8.10 |
| TYMS | 9644 | SNV | A/G | 3 | 520 | 0 | Synonymous | E127E | 323 | 280/42/1 | 6.81 | 211 | 185/24/2 | 6.64 |
| TYMS | 18164 | SNV | A/T | 5 | 771 | 0 | Nonsynonymous | Q211L | 323 | 322/1/0 | 0.15 | 211 | 211/0/0 | 0.00 |
| TYMS | 18255 | SNV | G/A | 5 | 862 | 0 | Synonymous | T241T | 323 | 323/0/0 | 0.00 | 211 | 210/1/0 | 0.24 |
| TYMS | 18770 | SNV | T/G | 6 | 872 | -7 | intron | | 323 | 322/1/0 | 0.15 | 211 | 211/0/0 | 0.00 |
| VANGL2 | 20276 | SNV | G/A | 2 | 498 | 0 | Synonymous | E4E | 316 | 316/0/0 | 0.00 | 210 | 209/1/0 | 0.24 |
| VANGL2 | 23492 | SNV | G/C | 4 | 742 | 0 | Nonsynonymous | D86H | 316 | 316/0/0 | 0.00 | 210 | 209/1/0 | 0.24 |
| VANGL2 | 23521 | SNV | G/A | 4 | 771 | 0 | Nonsynonymous | M95I | 316 | 315/1/0 | 0.16 | 210 | 210/0/0 | 0.00 |
| VANGL2 | 23559 | SNV | G/A | 4 | 809 | 0 | Nonsynonymous | G108D | 316 | 315/1/0 | 0.16 | 210 | 210/0/0 | 0.00 |
| VANGL2 | 23599 | SNV | C/T | 4 | 849 | 0 | Synonymous | L121L | 316 | 316/0/0 | 0.00 | 210 | 208/2/0 | 0.48 |
| VANGL2 | 23640 | SNV | G/A | 4 | 890 | 0 | Nonsynonymous | R135Q | 316 | 315/1/0 | 0.16 | 210 | 209/1/0 | 0.24 |
| VANGL2 | 23960 | SNV | C/G | 4 | 1210 | 0 | Nonsynonymous | L242V | 316 | 315/1/0 | 0.16 | 210 | 210/0/0 | 0.00 |
| VANGL2 | 23986 | SNV | C/G | 4 | 1236 | 0 | Synonymous | V250V | 316 | 315/1/0 | 0.16 | 210 | 210/0/0 | 0.00 |
| VANGL2 | 24871 | SNV | G/T | 5 | 1320 | 0 | Nonsynonymous | K278N | 323 | 322/1/0 | 0.15 | 211 | 211/0/0 | 0.00 |
| VANGL2 | 24879 | SNV | A/G | 5 | 1328 | 0 | Nonsynonymous | H281R | 323 | 323/0/0 | 0.00 | 211 | 210/1/0 | 0.24 |
| VANGL2 | 25546 | SNV | C/T | 6 | 1491 | 0 | Synonymous | D335D | 323 | 322/1/0 | 0.15 | 211 | 211/0/0 | 0.00 |
| VANGL2 | 28491 | SNV | G/A | 7 | 1572 | 0 | Synonymous | A362A | 320 | 319/1/0 | 0.16 | 210 | 210/0/0 | 0.00 |
| VANGL2 | 28524 | SNV | G/T | 7 | 1605 | 0 | Synonymous | L373L | 316 | 315/1/0 | 0.16 | 210 | 210/0/0 | 0.00 |
| VANGL2 | 28542 | SNV | A/G | 7 | 1623 | 0 | Synonymous | K379K | 321 | 104/154/63 | 43.61 | 210 | 86/90/34 | 37.62 |
| VANGL2 | 28647 | SNV | C/T | 7 | 1728 | 0 | Synonymous | Y414Y | 320 | 317/3/0 | 0.47 | 210 | 206/4/0 | 0.95 |
| VANGL2 | 29567 | SNV | C/T | 8 | 1814 | 0 | Nonsynonymous | A443V | 322 | 321/1/0 | 0.16 | 211 | 211/0/0 | 0.00 |
| VANGL2 | 29574 | SNV | A/C | 8 | 1821 | 0 | Synonymous | G445G | 322 | 188/119/15 | 23.14 | 211 | 148/52/11 | 17.54 |
| VANGL2 | 29633 | SNV | A/C | 8 | 1880 | 0 | Nonsynonymous | E465A | 322 | 321/1/0 | 0.16 | 211 | 211/0/0 | 0.00 |
| VANGL2 | 29640 | SNV | G/A | 8 | 1887 | 0 | Synonymous | P467P | 322 | 185/122/15 | 23.60 | 211 | 146/54/11 | 18.01 |
| VANGL2 | 29684 | SNV | G/A | 8 | 1931 | 0 | Nonsynonymous | R482H | 322 | 321/1/0 | 0.16 | 211 | 211/0/0 | 0.00 |
| WNT9B | 25963 | SNV | C/T | 2 | 162 | 0 | Nonsynonymous | A42V | 316 | 315/1/0 | 0.16 | 207 | 207/0/0 | 0.00 |
| WNT9B | 25964 | SNV | G/A | 2 | 163 | 0 | Synonymous | A42A | 316 | 315/1/0 | 0.16 | 207 | 206/1/0 | 0.24 |
| WNT9B | 25978 | SNV | A/G | 2 | 177 | 0 | Nonsynonymous | Q47R | 316 | 314/2/0 | 0.32 | 207 | 207/0/0 | 0.00 |
| WNT9B | 26006 | SNV | C/T | 2 | 205 | 0 | Synonymous | D56D | 316 | 314/2/0 | 0.32 | 207 | 206/1/0 | 0.24 |
| WNT9B | 26119 | SNV | G/A | 2 | 318 | 0 | Nonsynonymous | R94Q | 316 | 314/2/0 | 0.32 | 207 | 206/1/0 | 0.24 |
| WNT9B | 26129 | SNV | C/T | 2 | 328 | 0 | Synonymous | R97R | 316 | 315/1/0 | 0.16 | 207 | 207/0/0 | 0.00 |
| WNT9B | 26134 | SNV | A/G | 2 | 333 | 0 | Nonsynonymous | N99S | 316 | 315/1/0 | 0.16 | 207 | 207/0/0 | 0.00 |
| WNT9B | 26155 | SNV | C/T | 2 | 354 | 0 | Nonsynonymous | M106T | 316 | 148/131/37 | 32.44 | 207 | 87/91/29 | 35.99 |
| WNT9B | 28541 | SNV | G/A | 3 | 413 | 0 | Nonsynonymous | A126T | 317 | 314/3/0 | 0.47 | 210 | 210/0/0 | 0.00 |
| WNT9B | 28564 | SNV | G/T | 3 | 436 | 0 | Synonymous | R133R | 317 | 294/22/1 | 3.79 | 211 | 203/8/0 | 1.90 |
| WNT9B | 28717 | SNV | G/C | 3 | 589 | 0 | Nonsynonymous | K184N | 316 | 315/1/0 | 0.16 | 210 | 210/0/0 | 0.00 |
| WNT9B | 29670 | SNV | G/A | 4 | 664 | 0 | Synonymous | T209T | 316 | 313/3/0 | 0.47 | 210 | 208/2/0 | 0.48 |
| WNT9B | 29708 | SNV | G/A | 4 | 702 | 0 | Nonsynonymous | R222H | 316 | 315/1/0 | 0.16 | 210 | 210/0/0 | 0.00 |
| WNT9B | 29774 | SNV | C/T | 4 | 768 | 0 | Nonsynonymous | S244L | 316 | 314/2/0 | 0.32 | 210 | 210/0/0 | 0.00 |
| WNT9B | 29809 | SNV | T/C | 4 | 803 | 0 | Synonymous | L256L | 317 | 317/0/0 | 0.00 | 210 | 209/1/0 | 0.24 |
| WNT9B | 29889 | SNV | G/C | 4 | 883 | 0 | Synonymous | V282V | 317 | 299/17/1 | 3.00 | 210 | 199/11/0 | 2.62 |
| WNT9B | 30027 | SNV | C/T | 4 | 1021 | 0 | Synonymous | A328A | 316 | 315/1/0 | 0.16 | 210 | 210/0/0 | 0.00 |
